# A parallel glycolysis supports rapid adaptation in dynamic environments

**DOI:** 10.1101/2022.08.19.504590

**Authors:** Richard C. Law, Glenn Nurwono, Junyoung O. Park

## Abstract

Glycolysis is a universal metabolic process that breaks down glucose to produce cellular energy currency ATP and biomass precursors^1^. The Entner-Doudoroff pathway is a glycolytic pathway that parallels the textbook glycolysis but yields half as many ATP^2^. In organisms that possess both glycolytic pathways, such as *Escherichia coli*, inactivating the less energy-efficient Entner-Doudoroff pathway does not alter growth rates^3^. The benefit of the Entner-Doudoroff pathway has instead been hypothesized to be metabolic flexibility as an auxiliary enzyme-efficient catabolic route^4^. However, its *raison d’être* remains incompletely understood. Here we identify the advantage of employing parallel glycolytic pathways under dynamic nutrient environments. Upon carbon and nitrogen upshifts, wild-type cells accelerate growth faster than those with the Entner-Doudoroff pathway knocked out. Using stable isotope tracers and mass spectrometry, we find that the Entner-Doudoroff pathway flux increases disproportionately faster than that of the textbook glycolysis during nutrient upshifts. We attribute the fast response time of the Entner-Doudoroff pathway to its strong thermodynamic driving force and concerted regulation facilitating glucose uptake. Intermittent supply of nutrients manifests this evolutionary advantage of the parallel glycolysis. Thus, the dynamic nature of an ostensibly redundant pathway’s role in promoting rapid adaptation constitutes a metabolic design principle.

---

By survival of the fittest, modern organisms efficiently utilize available nutrients and rapidly adapt growth to changing environments^5^. Organisms across the domains of life use glycolysis, or the Embden-Meyerhof-Parnas (EMP) pathway, for efficient generation of biomass precursors and energy^1,6^. Glycolysis is regulated to respond to environmental perturbations and changing cellular demands by rapidly modulating its flux while maintaining metabolic homeostasis^7^.

Most organisms possess alternative glycolytic routes, such as the phosphoketolase pathway^8^, the pentose phosphate pathway, and, common to ∼25% of prokaryotes, the Enter-Doudoroff (ED) pathway (**Fig. 1a**)^2^. The EMP textbook glycolysis and the ED glycolysis are parallel from a carbon-centric perspective as both pathways convert one glucose into two pyruvate molecules. The difference is that compared to the textbook glycolysis, which yields two ATP and two NADH, the ED pathway yields one ATP, one NADH, and one NADPH (**Fig. 1b**). Despite the less efficient ATP generation, some microorganisms exclusively use the ED pathway due to its lower protein burden (i.e., fewer reaction steps) and higher exergonicity (i.e., greater thermodynamic driving force; **Fig. 1b**) compared to the textbook glycolysis^4,9,10^.

**Fig. 1:**
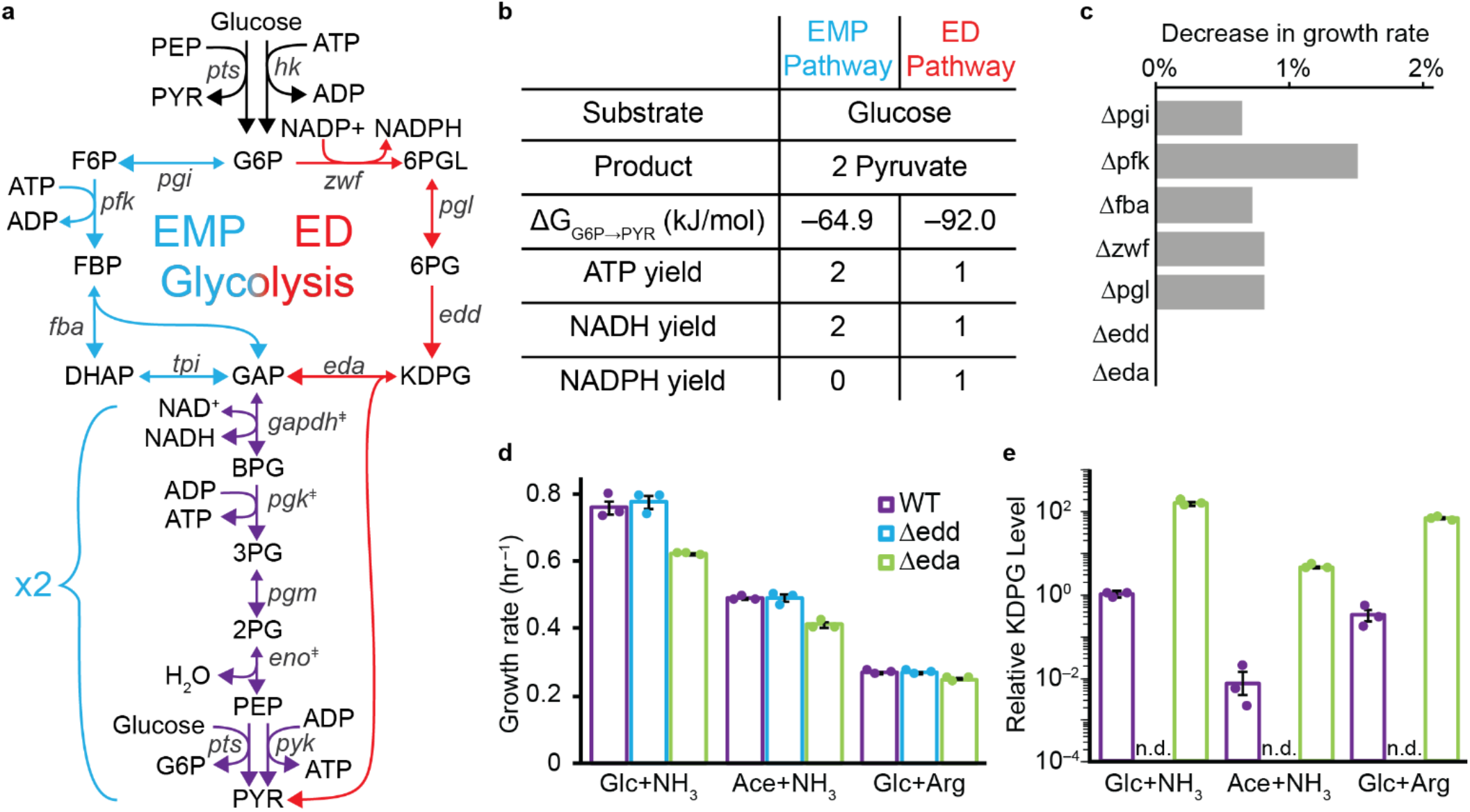
The ED pathway performs parallel glycolysis but does not affect cell growth rate. (**a**) The EMP glycolysis (red) and the ED glycolysis (blue) are parallel, and they share the steps of lower glycolysis (purple). Essential genes are marked with ‡. (**b**) The EMP and ED pathways are parallel carbon-wise with a bioenergetic difference. The ED pathway is thermodynamically more forward-driven (i.e., ΔG is more negative). (**c**) Flux balance analysis (FBA) of single non-essential gene knockouts predicted negligible impact of the ED pathway on cell growth. (**d**) The growth rates of WT and *Δedd* strains on different carbon and nitrogen sources were similar, but *Δeda* consistently grew slower. (**e**) The slow growth of Δ*eda* was due to the buildup of the ED pathway intermediate 2-keto-3-deoxy-6-phosphogluconate (KDPG). Error bars represent the standard error of the mean (s.e.m.) with n=3 biological replicates. G6P stands for glucose-6-phosphate; F6P, fructose-6-phosphate; FBP, fructose-1,6-bisphosphate; DHAP, dihydroxyacetone phosphate; GAP, glyceraldehyde-3-phosphate; 6PGL, 6-phosphogluconolactone; 6PG, 6-phosphogluconate; KDPG, 2-dehydro-3-deoxy-phosphogluconate; BPG, 1,3-bisphosphoglycerate; 3PG, 3-phosphoglycerate; 2PG, 2-phosphoglycerate; PEP, phosphoenolpyruvate; and PYR, pyruvate. *pts* represents the gene(s) for phosphotransferase system; *hk*, hexokinase; *pgi*, phosphoglucoisomerase; *pfk*, phosphofructokinase; *fba*, fructose bisphosphate aldolase; *tpi* triose phosphate isomerase; *zwf*, glucose-6-phosphate dehydrogenase; *pgl*, 6-phosphogluconolactonase; *edd*, phosphogluconate dehydratase; *eda*, 2-dehydro-3-deoxy-phosphogluconate aldolase; *gapdh*, glyceraldehyde-3-phosphate dehydrogenase; *pgk*, phosphoglycerate kinase; *pgm*, phosphoglycerate mutase; *eno*, enolase; and *pyk*, pyruvate kinase.

Interestingly, 14% of genome-annotated bacteria, including *E. coli*, possess both the EMP and the ED pathways^4^. Despite the apparent redundancy, the ED pathway facilitates sugar acid catabolism^11^ and genetic resilience^12–14^. The two glycolytic pathways additionally have different transcriptional regulation mechanisms^15^. The unique intermediate of the ED pathway 2-keto-3-deoxy-6-phosphogluconate (KDPG) deactivates transcriptional repressor YebK, which is implicated in determining the duration of the lag phase during nutrient downshift^16^. However, our understanding of the ED pathway is still limited compared to that of the EMP pathway. Furthermore, there is no clear experimental support for an evolutionary advantage of concurrent utilization of the parallel glycolytic pathways.

We were curious why an organism would retain and operate the ED pathway when it already has the EMP pathway that generates ATP more efficiently. We sought to identify conditions in which the ED pathway’s contribution to cells becomes meaningful. In stable nutrient environments, knocking out the ED pathway in *E. coli* had a negligible effect on cell growth. When we subjected the cells to nutrient upshift, however, the initial acceleration of growth of the ED-pathway-capable strain was significantly faster than that of the knockout strain. We found this difference to occur on a fast timescale (<10 mins.) during which cells would exhibit a metabolic response without substantial proteome reallocation.

To gain a mechanistic insight into the observed growth dynamics, we developed a strategy for discerning the fluxes through the two glycolytic pathways. Using stable isotope tracers and liquid chromatography-mass spectrometry (LC-MS), we observed that the ratio of ED to EMP pathway fluxes dynamically increased to 20% upon nitrogen upshift and 130% upon carbon upshift. Thus, the ED pathway aided cells in meeting the rapidly increasing energetic and carbon demand. This transitory metabolic benefit underlies the evolutionary advantage of parallel glycolytic pathways in intermittent nutrient availability over the short and long term. We surmise that organisms may employ parallel pathways elsewhere in metabolism to support rapid adaptation in dynamic environments.

## Results

### The ED pathway does not affect growth rates

To assess the contribution of the EMP and ED glycolytic enzymes on growth rates, we performed flux balance analysis (FBA) on the genome-scale metabolic model of *E. coli* (iAF1260)^17^. We simulated knocking out individual nonessential genes of the EMP and ED glycolysis and determined the maximum growth rates within the feasible flux space (**Fig. 1c**). Unlike the EMP pathway genes, which slowed growth upon knockout, the ED pathway genes had no effect on growth rates. We validated FBA predictions experimentally by comparing the growth rates of *E. coli* with and without the ED pathway genes (i.e., WT vs. Δ*edd* vs. *Δeda*) under various nutrient conditions (**Fig. 1d**). The WT and Δ*edd* strains had indistinguishable growth rates, corroborating our FBA simulations. We observed a 17% slowdown of growth in the Δ*eda* strain. This difference was attributable to a 100-fold buildup of the ED pathway intermediate KDPG in the Δ*eda* mutant (**Fig. 1e**), whose accumulation is correlated with bacteriostasis^18^.

The Δ*eda* strain accumulated KDPG to as much as 34 mM (**Supplementary Fig. 1**). Nonetheless, clean elimination of the ED pathway in the Δ*edd* strain showed that the ED pathway itself exerted a negligible direct effect on exponential growth rates.

### Parallel glycolysis accelerates growth

A key trait of modern organisms is their ability to quickly detect and utilize scarce nutrient resources once they become available^19,20^. Since WT and Δ*edd* strains showed no differences in stable environments, we investigated the role of the ED pathway during dynamic adaptation. We subjected these two strains to carbon or nitrogen upshift, a transition from a nutrient-limited to a replete state. Carbon limitation was achieved by using minimal media with a less favorable carbon source acetate in lieu of glucose. Acetate forces cells to use the glyoxylate shunt and gluconeogenesis to produce larger carbon backbones and supports slow growth^21,22^. Nitrogen limitation was introduced by culturing cells in a low initial NH_4_Cl concentration (2 mM), which gets depleted and stalls cell growth, or replacing NH_4_Cl with arginine, a less favorable nitrogen source. We induced upshift by spiking in glucose and NH_4_Cl to carbon- and nitrogen-limited cultures, respectively.

Upon upshift, both WT and Δ*edd* strains immediately increased growth rates (**Fig. 2**). However, the transition paths from slow to fast growth differed between the two strains. Upon carbon upshift, WT cells increased growth faster than Δ*edd* cells and maintained faster growth for 15 minutes until they both reached stable growth (**Fig. 2a** and **Supplementary Fig. 2**). This trend was similar in the arginine-to-NH_4_Cl nitrogen upshift, in which WT cells accelerated growth and maintained faster growth than Δ*edd* cells for an hour until the two strains reached the same stable growth rate (**Fig. 2b** and **Supplementary Fig. 3**). For NH_4_Cl depletion to repletion, WT displayed a higher initial acceleration of growth in the first five minutes, but the Δ*edd* strain outpaced WT in the subsequent 10 minutes (**Fig. 2c**). While the nitrogen upshifts brought both strains to the stable growth rate of ∼0.8 hr^−1^ measured in the nutrient-replete state, the carbon upshift did not. The incomplete recovery of the growth rate is due to the presence of high acetate (**Supplementary Fig. 4**). These observations suggested that the ED pathway may contribute to the faster growth acceleration in dynamic conditions.

**Fig. 2:**
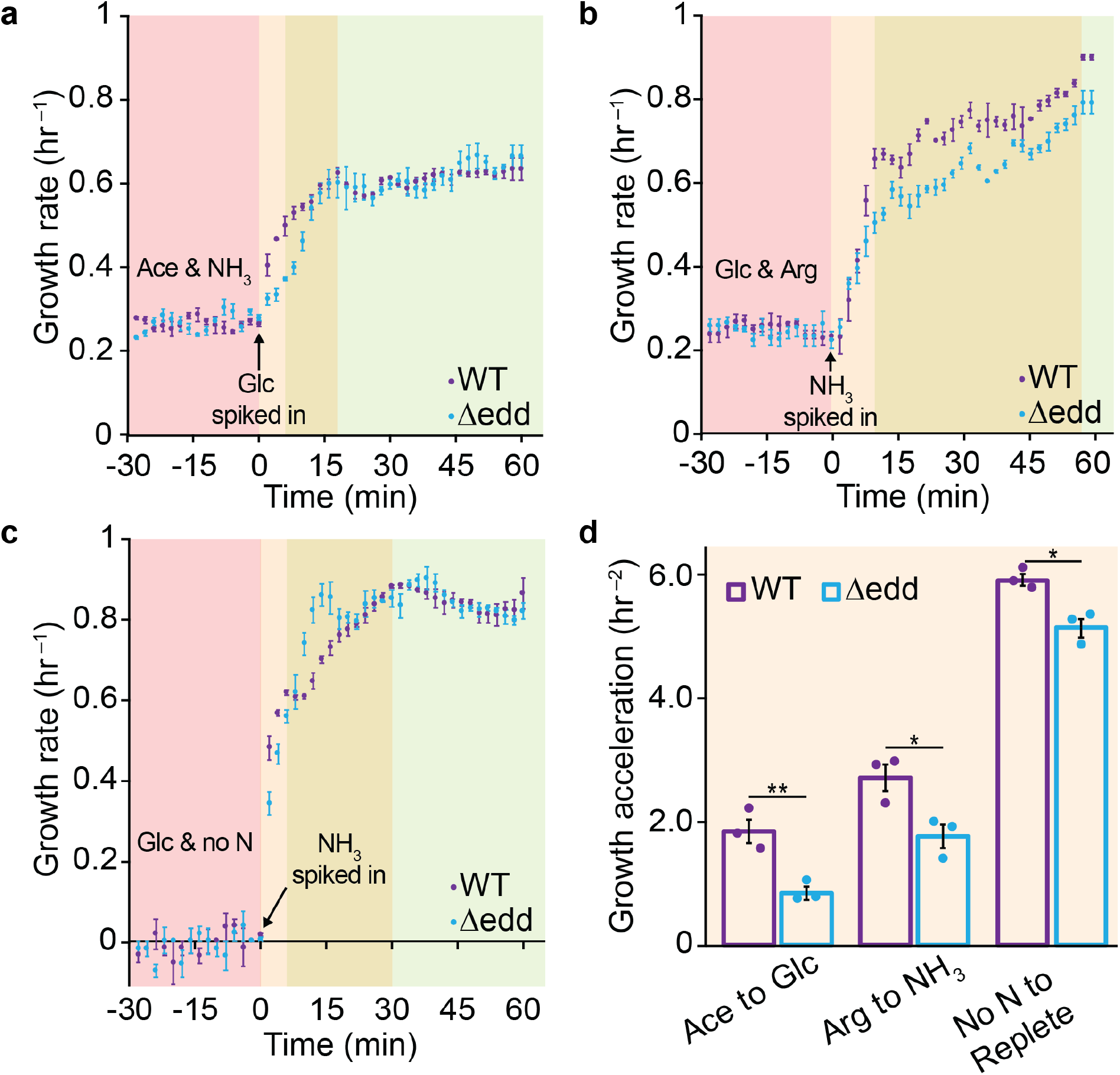
The ED glycolysis contributes to faster growth acceleration. (**a**) On acetate, WT and *Δedd* grow at the same rate. However, WT cells with the ED pathway increase growth rates faster than *Δedd* cells upon glucose addition. Both strains reach the same stable growth rate. Similarly, WT increases growth rate faster than *Δedd* upon arginine-to-ammonia nitrogen upshift. Both strains reach the same stable growth rate after 80 minutes (**Supplementary Fig. 3**). (**c**) Nitrogen upshift from ammonia depletion resulted in WT initially accelerating growth faster, but Δ*edd* overtook WT in the subsequent 10 minutes achieving stable growth sooner. (**d**) The maximum acceleration of growth occurred in the first few minutes of nutrient upshift. WT cells consistently accelerated faster than *Δedd* cells. Error bars represent the s.e.m. (n=3 biological replicates). * represents *P*<0.05 and ** represents *P*<0.01 by a two-tailed t-test.

To quantify this difference in growth dynamics, we computed growth acceleration, which is the time derivative of the specific growth rate (µ), which is the time derivative of log culture density. The maximal growth acceleration occurred in the first few minutes after upshift, and WT consistently outpaced Δ*edd* during this period (**Fig. 2d**). This rapid growth acceleration reflects metabolic rewiring that is driven by changes in metabolite levels rather than enzyme levels.

### Parallel glycolytic pathways activate upon upshift

We sought to gain mechanistic insights into the disparate growth acceleration with and without the ED pathway during nutrient upshift. To this end, we measured the levels of glycolytic intermediates and cofactors before and shortly after upshift using LC-MS (**Fig. 3a**). All three nutrient upshifts resulted in rapid changes in metabolite levels with the carbon upshift inducing the greatest overall change.

**Fig. 3:**
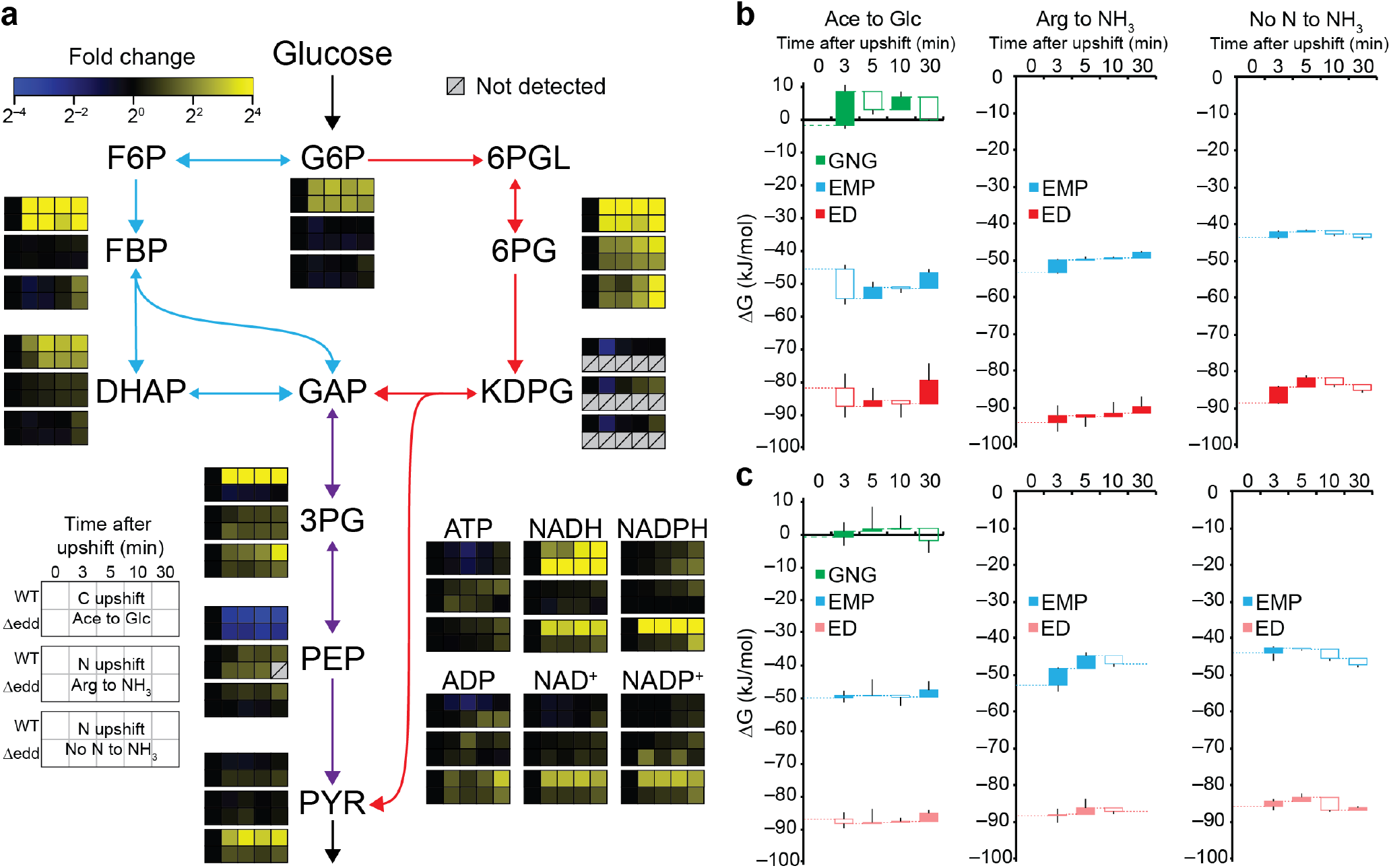
Both EMP and ED glycolytic pathways are activated upon nutrient upshift. (**a**) Most metabolites rapidly increased in WT and Δ*edd* strains following nutrient upshift. KDPG, an ED pathway specific intermediate, transiently decreased in the first five minutes of upshift and was not detected in the Δ*edd* strain. PEP decreased following glucose addition. (**b, c**) Gibbs free energy (ΔG) rapidly changed across the EMP pathway, the ED pathway, and gluconeogenesis (GNG) during upshift in (**b**) WT and (**c**) Δ*edd*. The ΔG of GNG immediately increased to positive value upon upshift in both strains. The EMP and ED glycolysis both remained thermodynamically forward-driven (ΔG<<0) during the transition. Since the ED pathway was inactive in Δ*edd*, its ΔG is shown in light red. The dotted lines represent the ΔG at the previous time points. Filled boxes indicate an increase in ΔG from the previous state and empty boxes indicate a decrease. Whiskers represent the s.e.m. (n = 3-6 biological replicates).

In both WT and Δ*edd*, carbon upshift increased the upper EMP glycolytic intermediates and 6-phosphogluconate (6PG) while lowering phosphoenolpyruvate (PEP), which had accumulated in the absence of glucose (**Fig. 3a**). The decrease in PEP levels reflected the unblocking of the phosphotransferase system (PTS), which phosphorylates glucose using PEP while transporting glucose into the cell, upon glucose addition^23^. In the WT strain, Gibbs free energy change (ΔG) across the EMP and the ED glycolysis became more forward-driven (i.e., ΔΔG<0) to –54.8 kJ/mol and –87.9 kJ/mol, respectively, while ΔG of gluconeogenesis increased to 8.6 kJ/mol (ΔΔG=+10.8 kJ/mol) in three minutes of carbon upshift (**Fig. 3b**). Thus, the rapid metabolic shifts rendered glycolysis thermodynamically more favorable and gluconeogenesis unfavorable in WT. On the other hand, thermodynamic shifts in Δ*edd* were less pronounced (**Fig. 3c**).

Nitrogen upshift increases glycolytic flux through a combination of upregulation of glucose transport by PTS by decreasing α-ketoglutarate (αKG), which inhibits Enzyme I of the PTS, and decreasing the reversibility of glycolysis^24^. Upon nitrogen upshift, the intermediates of glycolysis changed to a smaller extent in WT compared to carbon upshift (**Fig. 3a**), and as a result, we observed smaller ΔΔG in both the EMP and the ED glycolysis (**Fig. 3b**). ΔΔG of glycolytic pathways for Δ*edd* in nitrogen upshift was larger than carbon upshift but otherwise comparable to nitrogen upshift in WT (**Fig. 3c**). While the initial values of overall ΔG of the EMP and ED pathways were substantially different, their ΔG changed similarly upon carbon and nitrogen upshift due to the similarity in the initial substrates and final products of the parallel pathways. However, the intermediate metabolites unique to each pathway (e.g., fructose-1,6-bisphosphate and KDPG), which do not contribute to overall pathway thermodynamics, displayed disparate dynamics (**Fig. 3a**), suggesting different timescales at play in modulating the two glycolytic pathways.

### Asymmetric ^13^C-glucose reveals glycolytic fluxes

To quantify fluxes through the parallel glycolytic pathways, we needed a glucose tracer that differentially labels lower glycolytic metabolites depending on the pathway taken. Unlike the EMP glycolysis, in which each glucose produces a pair of triose phosphates (glyceraldehyde-3-phosphate, GAP) by cleaving fructose bisphosphate down the middle, the ED glycolysis shunts only the fourth, fifth, and sixth carbons of glucose to the shared lower glycolysis as a triose phosphate (**Fig. 4a**). With [1,2-^13^C_2_]glucose, the EMP pathway generates unlabeled (M+0) and doubly labeled (M+2) triose phosphate at a one-to-one ratio while the ED pathway generates only M+0 triose phosphate. The EMP versus ED glycolysis also result in unique positional isotope labeling (**Fig. 4a**) that can be distinguished by MS/MS fragmentation of valine (**Fig. 4b** and **Supplementary Fig. 5**). Furthermore, tracing [1,2-^13^C_2_]glucose informs us of the oxidative pentose phosphate pathway (OxPPP) flux, which uniquely generates singly labeled (M+1) triose phosphate (**Supplementary Fig. 6**). In *E. coli* exponentially growing in a glucose minimal medium, 3-phosphoglycerate (3PG) and valine labeling measurement indicated a slow ED pathway flux relative to the EMP glycolysis (**Fig. 4a,b**).

**Fig. 4:**
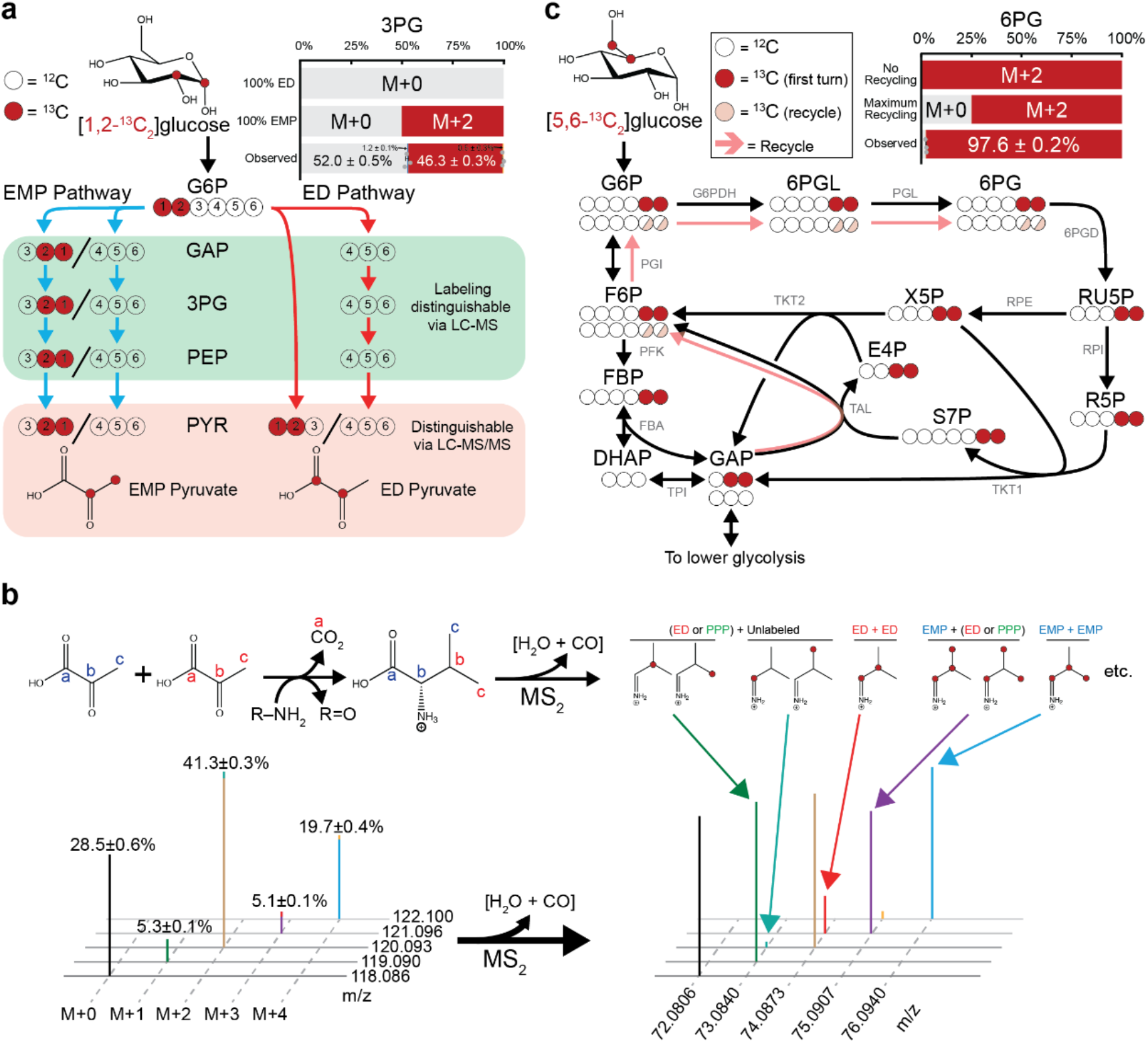
Asymmetrically ^13^C-labeled glucose tracers reveal glycolytic fluxes. (**a**) With [1,2-^13^C_2_]glucose, only the EMP glycolysis generates labeled lower glycolytic intermediates. Both the EMP and the ED pathways produced labeled pyruvate but at different positions. The labeling fraction of 3PG indicated the ED pathway activity, albeit much smaller than that of the EMP pathway, in cells without any nutrient limitation. (b) Positional labeling of pyruvate can be distinguished using LC-MS/MS. Each of the isotopologues of valine, which is synthesized from two pyruvate molecules, were fragmented. The MS2 spectra revealed the positional labeling that is indicative of the ED pathway activity. (c) The unlabeled fraction of 6PG from [5,6-13C2]glucose tracing revealed a small degree of recursive pentose phosphate pathway (PPP) usage, which we used to more accurately compute glycolytic fluxes. Unlabeled 6PG is only generated from S7P and unlabeled GAP through the reactions in light red arrows. Error bars represent the s.e.m. (n=3 biological replicates). RU5P stands for ribulose-5-phosphate; X5P, xylulose-5-phosphate; R5P, ribose-5-phosphate; E4P, erythrose-4-phosphate; S7P, sedoheptulose-7-phosphate; PGI, phosphoglucoisomerase; PFK, phosphofructokinase; FBA, fructose-1,6-bisphosphate aldolase; TPI, triose phosphate isomerase; G6PDH, glucose-6-phosphate dehydrogenase; PGL, phosphogluconolactonase; 6PGD, 6-phoshogluconate dehydrogenase; RPE, ribulose phosphate epimerase; RPI, ribose phosphate isomerase; TKT1 and TKT2, transketolase; and TAL, transaldolase.

The non-oxidative pentose phosphate pathway (non-OxPPP) and the reversibility of phosphoglucoisomerase (PGI) potentially obscures our flux quantitation by opening the door for carbons to recursively go through the PPP and produce various isotopomers of triose phosphate^25,26^. To disambiguate the central carbon metabolism fluxes, we also traced [5,6-^13^C_2_]glucose (**Fig. 4c**). Similar to [1,2-^13^C_2_]glucose, [5,6-^13^C_2_]glucose indicates the relative glycolytic fluxes based on the skewed labeling ratios but with the ED pathway flux contributing to M+2 triose phosphate (**Supplementary Fig. 7**). Nonetheless, [5,6-^13^C_2_]glucose divulges the extent of the PPP recycling by uniquely generating unlabeled 6PG. Our 6PG labeling measurement indicated a small recursive PPP flux (**Fig. 4c**). Additionally, since [5,6-^13^C_2_]glucose does not undergo labeling rearrangement in the PPP, balances on the two isotopomers it generates can also be used to measure glycolytic fluxes (**Supplementary Fig. 8**). Thus, [1,2-^13^C_2_]- and [5,6-^13^C_2_]glucose, which are labeled asymmetrically around the cleavage site, reveal the central carbon metabolism fluxes (**Supplementary Notes 2-4**). We determined the ED-to-EMP flux ratio using the three approaches ([1,2-^13^C_2_]glucose labeling of 3PG, correction with [5,6-^13^C_2_]glucose, and [1,2-^13^C_2_]glucose fragmentation of valine. From these methods, we found the nutrient replete ED-to-EMP flux ratio to be in the range of 0.06 to 0.12.

### The ED pathway accelerates faster upon upshift

We hypothesized that the adaptive (but not stable) growth advantage of the WT over Δ*edd* was due to the ED pathway enabling higher glycolytic flux in transitional periods. Using [1,2-^13^C_2_] and [5,6-^13^C_2_]glucose, we set out to investigate how the use of the parallel glycolytic pathways changed during carbon and nitrogen upshift. Glycolytic intermediates typically reach a stable labeling within a minute (**Supplementary Fig. 9**) due their fast turnover (∼1 s^−1^) (**Supplementary Fig. 10**). Thus, their isotopic labeling can be approximated as a series of minute-scale pseudo-steady states. We measured the isotope labeling of 3PG and 6PG from [1,2-^13^C_2_]- and [5,6-^13^C_2_]-glucose tracers before and 3, 5, 10, and 30 minutes after upshift (**Fig. 5a,b** and **Supplementary Fig. 11**).

**Fig. 5:**
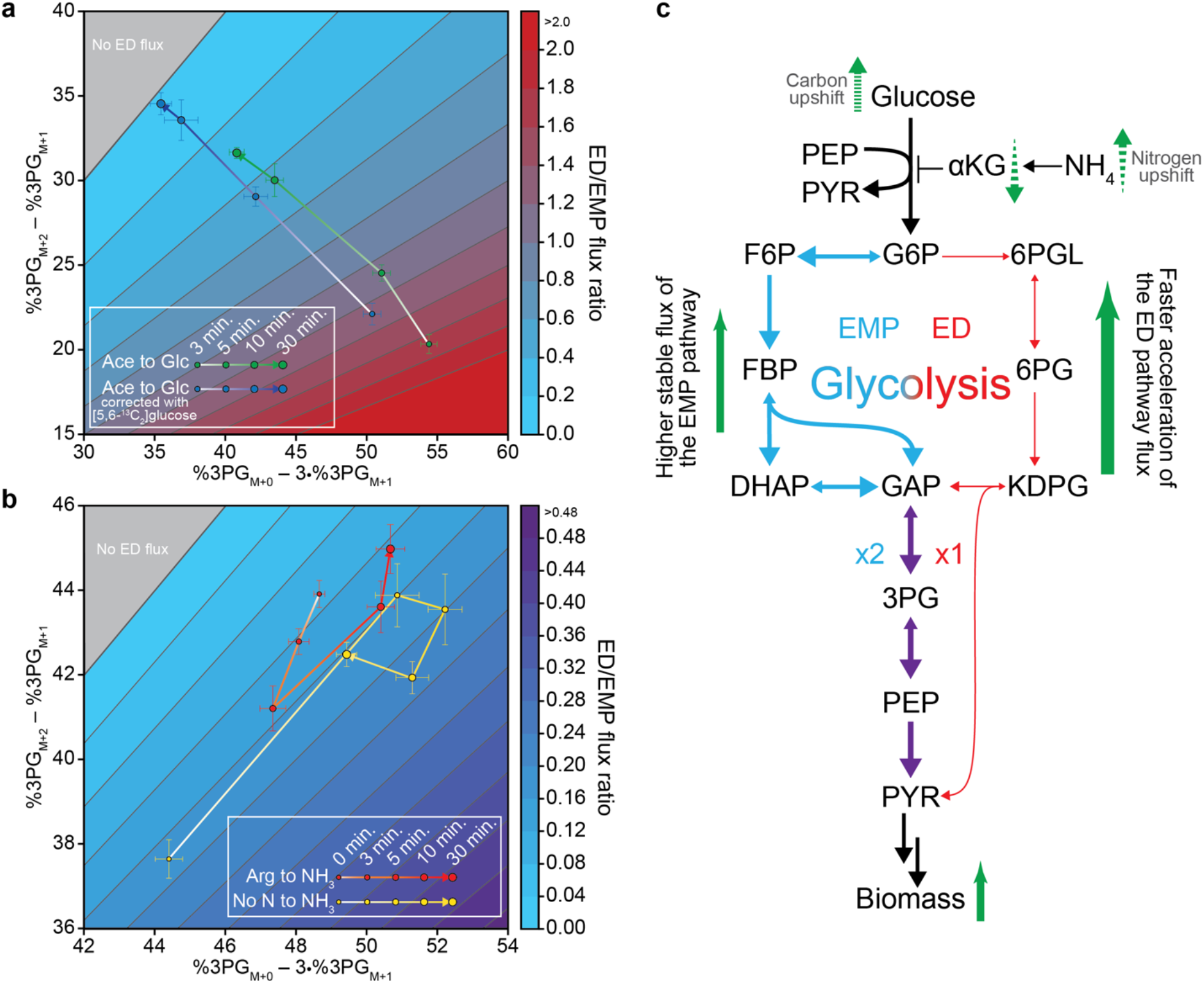
ED pathway flux accelerates faster than the EMP glycolytic flux. (**a**) The ED-to-EMP flux ratios were obtained using the 3PG labeling from [1,2-^13^C_2_]glucose (green) upon carbon upshift. Due to some residual gluconeogenic activity, the 3PG labeling fractions were corrected using the mirror-image tracer [5,6-^13^C_2_]glucose (blue) (see **Methods** and **Supplementary Note 4**). The ED pathway is the dominant glycolysis immediately following glucose addition. (**b**) Upon nitrogen upshift from arginine (red) and no nitrogen (yellow), the ED pathway flux accelerated faster than the EMP pathway within five minutes. Their flux ratios returned to the initial ratios after 30 minutes. (**c**) The EMP and the ED pathways have complementary roles. The EMP pathway excels in maintaining high homeostatic glycolytic flux while the ED pathway excels in rapidly bolstering glycolytic flux upon increased biomass and energy demand. Error bars represent the s.e.m. (n=3 biological replicates).

The ED pathway’s contribution to overall glycolysis rapidly increased upon carbon and nitrogen upshift. Upon carbon upshift, the ED pathway carried the major glycolytic flux (**Fig. 5a**). The relative EMP glycolytic flux increased over time. Although we could not quantify the ED pathway flux in the carbon-limited condition due to the absence of ^13^C glucose tracers, we found that even on acetate *E. coli* expressed the ED pathway enzymes (**Supplementary Fig. 12**). Since any unlabeled glycolytic intermediates that had been derived from unlabeled acetate affect our ability to compute glycolytic fluxes accurately, we took extra caution by performing acetate-to-[U-^13^C_6_]glucose upshift to account for residual acetate-sourced intermediates (**Supplementary Fig. 13** and **Supplementary Note 4**). Both nitrogen upshifts from arginine and low NH_4_Cl rapidly increased the relative ED pathway flux (**Fig. 5b**). The extent of the ED pathway utilization during nitrogen upshift was smaller than that of carbon upshift, demonstrating the benefit of a reliable glycolytic route even in thermodynamically less favorable times of early carbon upshift.

Taken together, our flux quantitation revealed that while the EMP pathway is the main stable glycolytic route, the ED pathway responds faster to upshift glycolysis (**Fig. 5c**). This faster acceleration of the ED pathway flux was consistent with our earlier observation that the ED pathway-capable WT accelerated growth faster than the Δ*edd* ED pathway knockout strain even though they grew equally well under stable nutrient conditions.

### Intermittent feeding favors parallel glycolysis

We were curious about the evolutionary benefit that the transiently faster growth acceleration may confer on *E. coli* with both EMP and ED glycolysis. In nature, access to favorable nutrients is often limited because their supply is intermittent and subject to competition within microecosystems^27^. We cultured WT and Δ*edd* strains on low glucose (**Fig. 6a**) or low ammonia (**Fig. 6b**) until the limiting nutrient was depleted and spiked in a low dose of the nutrient repeatedly after nutrient depletion.

**Fig. 6:**
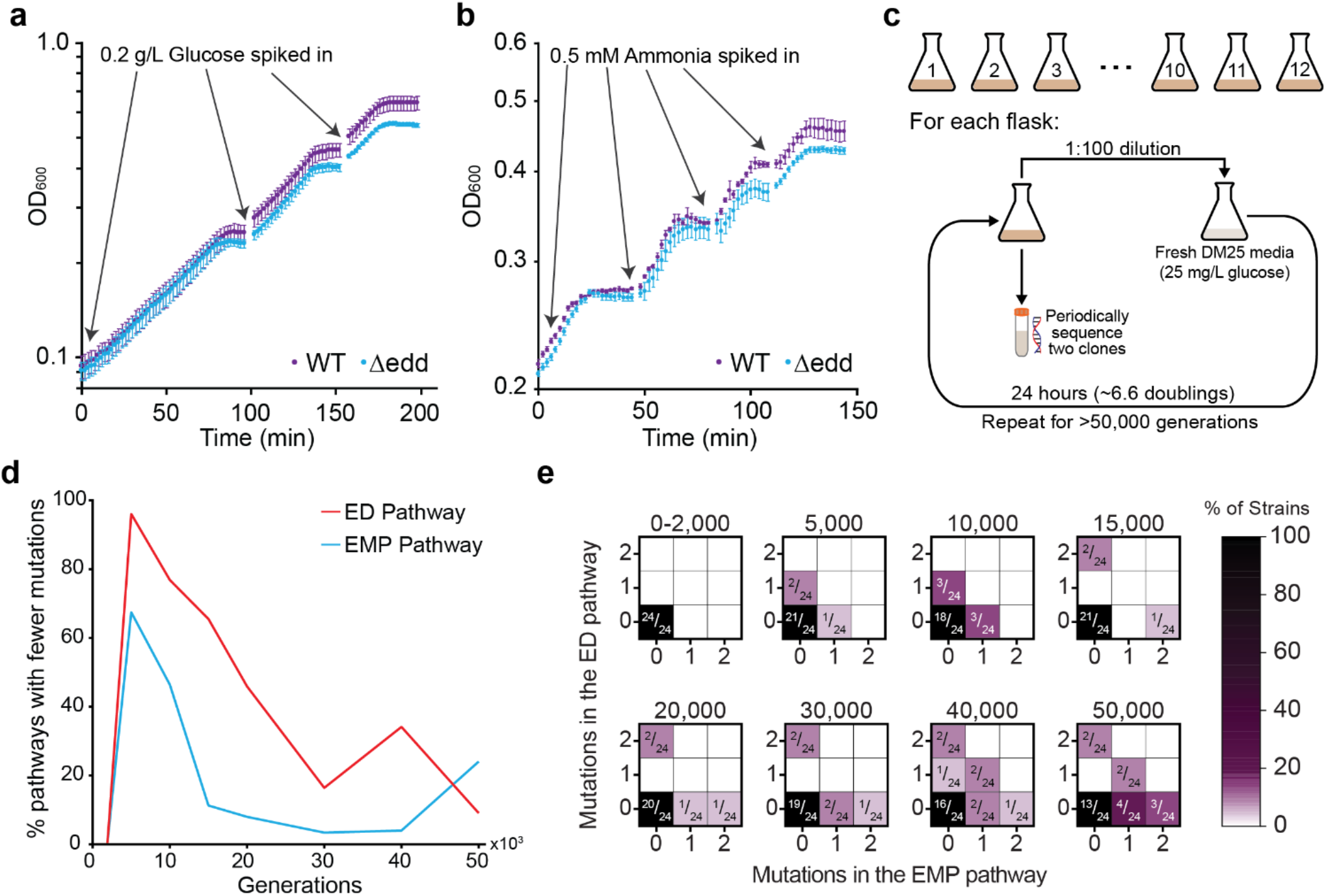
Importance of the ED pathway persists in short- and long-term intermittent nutrient feeding. (**a**) WT outperformed the Δ*edd* strain under intermittent supply of glucose. With each successive upshift, WT further pulled away from Δ*edd* in terms of culture density. (**b**) Similarly, intermittent feeding of ammonia led to higher growth in WT culture density. (**c**) The long-term evolution experiment mirrored intermittent glucose supply. 12 starter cultures of *E. coli* were transferred to fresh low-glucose media daily, inducing carbon upshift for >50,000 generations. From each culture, two clones were periodically sampled and sequenced. (**d**) The EMP and ED pathways accumulated mutations early on compared to other pathways. At generation 5,000, the ED pathway had a higher frequency of mutations than 96% of pathways. (see **Methods**). (**e**) The EMP and ED pathway mutations develop over time in parallel. No strains developed mutations in both pathways until generation 40,000. Error bars represent the s.e.m. (n=3 biological replicates).

Under these intermittent nutrient feeding conditions, cells possessing both glycolytic pathways achieved increasingly higher culture densities than Δ*edd* cells (**Fig. 6a,b**). In three cycles of glucose nutrient feeding, WT achieved a cell density of ∼13 % higher than Δ*edd*. Similarly, four cycles of ammonia upshift widened the growth gap between the two strains to ∼5 %.

We further investigated the evolutionary relevance of the ED pathway using the *E. coli* long-term evolution experiment (LTEE) by Lenski et al^28^. In the LTEE, twelve initial populations derived from REL606 and REL607 strains have been intermittently fed limiting amounts (25 mg/L) of glucose over 70,000 generations. The low glucose concentration led to glucose depletion, and daily subculturing into fresh media introduced carbon upshift every 6.6 generations (**Fig. 6c**).

We analyzed the genome sequences of the twelve populations over 50,000 generations, which is the latest generation with readily available genomic data. The mutation history was suggestive of increasing adaptability as we observed more than expected genes with two or more mutations (**Supplementary Fig. 14**). Some genes even accumulated as many as 50 mutations (**Supplementary Fig. 15**). If random mutations accumulated, the distribution of total number of mutations for a gene would closely match the Poisson distribution.

We compared the mutation history of the EMP and the ED glycolytic pathways relative to other pathways. The ED pathway was highly mutated early on, reaching the 96^th^ percentile of pathways by 5,000 generations, while the EMP pathway was at the 67^th^ percentile (**Fig. 6d** and **Supplementary Fig. 16**). At 5,000 generations, none of the mutations were silent (**Supplementary Fig. 17**). In subsequent generations, the percentile ranks of both glycolytic pathways decreased, but the fractions of nonsynonymous and indel mutations remained above 50%. With mutations in earlier generations generally leading to a greater increase in the fitness of an organism^29,30^, the early mutations in the ED pathway manifested its importance under intermittent glucose supply. Up to 30,000 generations, mutations occurred in either of the two glycolytic pathways (**Fig. 6e**). Furthermore, by 50,000 generations, more than 80% of those strains that had mutations had only one of the two glycolytic pathways mutated (**Supplementary Fig. 18**). Taken together, these observations hint at the distinct utility of the EMP and the ED glycolysis under stable and transitional periods.

## Discussion

Parallel reactions and pathways are prevalent in metabolism, yet the evolutionary advantage of their concurrent utilization remains incompletely understood. In bacteria, the phosphotransferase system (PTS) carries out the same function as the glucose transporter, hexokinase, and pyruvate kinase^31^. In eukaryotes, parallel pathways are commonly found in different organelles (e.g., one-carbon metabolism and partially the TCA cycle in the cytosol and mitochondria)^32–34^. Genetic redundancy is commonplace in diploids^14^. In all organisms, multiple isoforms of various enzymes exist.

The EMP and the ED pathways are parallel glycolytic pathways that perform perhaps the most foundational task in metabolism. In an organism with both glycolytic pathways, it is expected that they should both play important roles, lest one of them would have been deprecated. Interestingly, in stable environmental conditions, the ED pathway flux in *E. coli* is a small fraction of the EMP glycolysis, and when we knocked out the ED pathway, we did not find any growth defect under steady-state culture conditions.

Instead, we found the benefit of the ED pathway to manifest during dynamic and transitory responses upon nutrient upshift. In the first few minutes of carbon and nitrogen upshift, cells with the ED pathway accelerated growth faster than those without. The quantitation of ED-to-EMP pathway flux ratios revealed the rapid increase of the relative ED pathway flux concurrently with the growth acceleration. The stronger thermodynamic driving force and fewer enzymatic steps of the ED pathway facilitate its rapid preferential upregulation compared to the EMP pathway. These observations suggest that the ED glycolysis is designed to jump once limiting nutrients become available, whereas the EMP glycolysis, although slower to respond to nutrient upshift, is designed to efficiently generate ATP under steady state. The parallel glycolytic pathways are thus complementary and ensure that cells are evolutionarily competitive in both stable and dynamic environments.

The dynamics of growth acceleration differed slightly across our nutrient upshift experiments. We attribute the differences to disparate thermodynamic driving forces and regulatory mechanisms at play^35^. In the carbon upshift experiment, the absence of glucose initially forces cells to halt glycolysis and use acetate for gluconeogenesis. The addition of glucose initiates a switch from gluconeogenesis to glycolysis, which requires flipping the sign of ΔG of every EMP glycolysis reaction step that participates in gluconeogenesis^7^. On the other hand, the ED pathway steps do not participate in gluconeogenesis due to highly negative ΔG°’ and thus can more rapidly increase its flux upon glucose addition. The ED pathway allows NADPH generation and oxidative stress response like the OxPPP^36^ but without decarboxylation. Thus, in addition to facilitating rapid glycolytic response, another role of the ED pathway is carbon-efficient NADPH generation (**Supplementary Fig. 19**). By feeding [1-^2^H]- and [3-^2^H]-glucose to cells, we observed NADPH production through glucose-6-phosphate dehydrogenase (shared between the ED pathway and the OxPPP) that is three times as high as that of the OxPPP exclusive step 6-phosphogluconate dehydrogenase (**Supplementary Fig. 20**). This deuterated glucose labeling experiment revealed that the contributions of the ED pathway and the OxPPP to NADPH production are comparable.

The mechanism of increasing glycolytic flux upon nitrogen upshift is different from that of carbon upshift. In nitrogen limitation, cells voluntarily limit their glucose import via competitive inhibition of the PTS EI by αKG. The PTS system is the main glucose importer and requires PEP as a substrate to transport. In glucose limitation, PEP accumulates and acts as a reserve pool ready to increase the PTS flux as soon as glucose becomes available^23^. This is not the case for nitrogen limitation and upshift. Our observations suggest that cells may procure PEP by rapid increase of the ED pathway flux. The ED pathway converts glucose to pyruvate in five reaction steps, and pyruvate can be converted to PEP via phosphoenolpyruvate synthase (PpsA). PpsA is also regulated by nitrogen availability with αKG acting as its inhibitor that blocks phosphotransferase^37^. While PpsA sacrifices ATP, which is not a concern for nitrogen-limited cells, the concerted regulation of PpsA and PTS facilitates streamlined glucose transport that supports a rapid increase of glycolytic flux. The ED pathway plays a crucial role in bridging PTS and PpsA via a productive shortcut (**Supplementary Fig. 21**).

The short- and long-term intermittent feeding experiments corroborated the benefit of parallel glycolysis. Repeated nutrient upshifts manifested the transitory yet evolutionary benefit of the ED pathway in rapid acceleration of glycolysis for energy and biomass precursor generation. The Lenski long-term evolution experiment, which provided the accumulation of beneficial and neutral mutations over 50,000 generations of glucose upshift, demonstrated the comparable importance of the EMP and the ED pathways. Had the ED pathway not played an important role in repeated glucose upshifts, its mutation history would be unlikely to see the high activity in early generations.

In nature, rapid nutrient upshift is a common occurrence and nutrients are gradually depleted by organisms. Bacteria readily adapt to these nutrient fluctuations^38^ and have evolved to cope with short-term changes^39^. In addition to already discovered strategies, our results indicated the manifold benefits of the parallel glycolysis for recovering from nutrient depletion. More generally, parallel pathways afford flexibility. By employing different cofactors and substrate binding affinities attuned to different environments, parallel reaction steps contribute to metabolic homeostasis under various conditions and stresses^40^. This may also help mitigate the impacts of when flux of one of the two pathways may become misregulated^41^. Enzymes catalyzing the same reactions may also possess different regulatory mechanisms as in the case of self-resistance enzymes that protect cells from natural product inhibitors they themselves produce^42,43^.

Glycolysis is often the highest flux-carrying metabolic pathway in heterotrophic organisms. The co-existence of the EMP and the ED glycolytic pathways acts as structural support for rapid glycolytic flux control. Even though the ED pathway flux in *E. coli* was relatively small compared to the EMP glycolysis flux, the ED pathway still carried a substantial flux comparable to other pathways that branch off of central carbon metabolism. Such “hardwired” flux control mechanisms add to the arsenal of rapid adaptation strategies that include small-molecule-based regulation, thermodynamic shift, and post-translational modification^44,45^. Alternative glycolytic pathways are not unique to prokaryotes^46^, and parallel pathways are commonly found in metabolic maps. Thus, we postulate that dynamic pathway responses underlie a metabolic design principle.

## Methods

### Strains and culture conditions

*E. coli* K-12 strain NCM3722 was the wild type (WT) in this study. The ED pathway knockout strains Δ*edd* and Δ*eda* with the NCM3722 background were produced by P1 phage transduction^47^ of a deletion allele from the Keio collection^48^. *E. coli* were grown in Gutnick minimal media^49^ at 37 °C. Media contained either 0.2% (w/v) glucose or 0.273% (w/v) sodium acetate as the carbon source such that the same molar availability of elemental carbon was achieved. For the nitrogen source, media contained 10 mM NH_4_Cl for the nitrogen-replete condition and either 2 mM NH_4_Cl or 2.5 mM of arginine for the nitrogen-limited conditions. Culture density (OD_600_) was monitored by spectrophotometer or plate reader.

For carbon upshift, cells were initially grown on acetate. When cultures reached the mid-log phase of growth (OD_600_ ≈ 0.3), carbon upshift was performed by spiking in a concentrated glucose stock solution into the culture to a final glucose concentration of 0.2% (w/v). For nitrogen upshift, cells were initially cultured in media containing 2 mM NH_4_Cl or 2.5 mM arginine until mid-log phase. At OD_600_ ≈ 0.3, cells consumed most of the nitrogen from NH_4_Cl, slowing down cell growth, and arginine cultures were in the mid-log phase. To induce upshift, NH_4_Cl was spiked into cultures to a final concentration of 10 mM. Metabolism was quenched and metabolites were extracted immediately prior to upshift (0 minutes) as well as 3, 5, 10, and 30 minutes after upshift. For isotope labeling experiments, unlabeled glucose was replaced with [1,2-^13^C_2_]-, [5,6-^13^C_2_]-, or [U-^13^C_6_]-glucose.

### Flux balance analysis of single gene glycolysis knockouts

Flux balance analysis was performed using the COnstraint-Based Reconstruction and Analysis Toolbox (COBRA) on MATLAB with the *E. coli* genome-scale reconstruction iAF1260^50^. The objective function was set to maximize biomass production while satisfying the mass balance constraints and carbon uptake rates specified by:

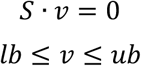

*S* is the stoichiometric matrix and *v* is the vector corresponding to reaction fluxes. *lb* and *ub* are the lower and upper bounds of *v* based on biochemical and thermodynamic considerations. Individual genes of the EMP or ED glycolysis were silenced by constraining the respective reaction’s flux bounds to 0 mmol/h/gDCW. For simulations of growth on glucose, the glucose uptake rate was set to 8 mmol/h/gDCW to reflect typical substrate uptake rates^50^.

### Nutrient upshift growth assays

To measure growth rates and accelerations during nutrient upshift, cells were grown in 96-well plates in a plate reader (BioTek) with shaking at 37 °C. Culture density (OD_600_) measured every two minutes. For each biological replicate, the median value of technical replicates across multiple wells was taken for growth analysis. Growth acceleration was computed as the time derivative of the specific growth rate (µ), which is the time derivative of log culture density (ln(C)):

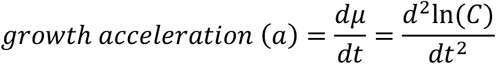

Intermittent nutrient upshift growth curves were performed by culturing cells in a 96-well plate in the plate reader. Cultures were first inoculated into plates with either glucose- or NH_4_Cl-depleted media before 0.02% (w/v) glucose or 0.5 mM NH_4_Cl was added at t = 0 minutes. Upshift was monitored until the limiting nutrient was depleted and cell growth ceased for no longer than 15 minutes. For nutrient upshift, the plate was taken out from the plate reader for spiking in 0.02% (w/v) glucose or 0.5 mM NH_4_Cl. These nutrient spike-in and growth monitoring steps were repeated a few times. All growth assays were performed in biological triplicates and, for each biological replicate, 12 technical replicates.

### Metabolite extraction and measurement

Metabolite extraction was conducted as quickly as possible to minimize perturbations in metabolism. To quickly quench metabolism and extract metabolites, 1 mL of cultures was vacuum-filtered onto nylon membrane filters (0.45 µm; Millipore) and flipped cell-side down into 400 µL of 40:40:20 HPLC-grade acetonitrile/methanol/water that was pre-cooled to –20 °C in a 6-well plate. Extraction continued at –20 °C for 20 minutes before the filter was flipped cell-side up and washed with the extraction solvent in the well. The extract was collected in an Eppendorf tube and centrifuged at 4 °C. The supernatant was dried under nitrogen flow and reconstituted in HPLC-grade water for LC-MS analysis.

The metabolite extract samples were analyzed by high performance liquid chromatography (Vanquish Duo UHPLC, Thermo) coupled to a high-resolution orbitrap mass spectrometer (Q Exactive Plus, Thermo). The LC separation was achieved using a hydrophilic interaction chromatography column (XBridge BEH Amide XP Column, 130 Å, 2.5 µm, 2.1 mm × 150 mm, Waters). Mass spectrometry was performed in both positive and negative mode using a mass resolution of 140,000 at 200 m/z. The resulting LC-MS data was processed using the Metabolomic Analysis and Visualization Engine (MAVEN)^51^ with peaks identified by both the known retention times and mass-to-charge ratios (m/z)^52^.

For LC-MS/MS analysis of valine, the same LC method was used with a modified MS protocol with a full MS/data-dependent MS^2^ scan using a normalized collision energy (NCE) of 35. The positive parent ions of all valine isotopologues were inputted for MS^2^ fragmentation. The resulting LC-MS/MS data was analyzed on MAVEN by first identifying the parent m/z for each valine isotopologue and then extracting its fragment spectra^51^.

### Quantitation of Gibbs free energy of reaction

Absolute metabolite concentrations in different conditions and time points were obtained by comparing peak areas to the known reference points in which the absolute concentrations of central carbon metabolites of *E. coli* had been measured^53,54^. The KDPG concentration was measured using an isotope-ratio-based approach^55^. Cellular metabolites were labeled by culturing *E. coli* on [U-^13^C_6_]glucose for multiple generations and extracted using the extraction solvent containing known concentrations of the unlabeled KDPG internal standard.

Using absolute metabolite concentrations (**Supplementary Table 1**), Gibbs free energy of reaction (ΔG) was computed using the following equation.

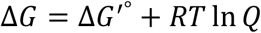

ΔG’° is ΔG at standard biochemical conditions, R is the universal gas constant, T is the temperature in kelvins, Q is the reaction quotient (i.e., the ratio of product-to-substrate activities, which are effective concentrations in a non-ideal solution). ΔG and changes in ΔG from one state to another (ΔΔG) were computed for the EMP pathway, the ED pathway, and gluconeogenesis (**Supplementary Note 1**).

### Cell lysate assay

Measurement of the ED pathway activity in acetate cultures was conducted by monitoring KDPG production from cell lysates upon adding 6PG. Cells were grown on acetate until cultures reached ∼0.4 OD_600_. Cells were pelleted, washed twice in cold phosphate-buffered saline (PBS), and resuspended in PBS. Cells were lysed by addition of 20 mg/mL lysozyme and sonication before subsequent centrifugation for 10 minutes. All lysis steps were conducted at 4 °C. The supernatant was moved to an Eppendorf tube and heated to 37 °C. 5 mM 6PG was added to the cell lysate and the reaction mixture incubated at 37°C with continuous shaking. Small aliquots of the reaction mixture were sampled over time. The reaction in those aliquots was quenched by the addition of cold 40:40:20 methanol:acetonitrile:water at a 1:4 ratio of the aliquot to the quenching solution. The mixture was centrifuged, and the supernatant was taken for the measurement of KDPG by LC-MS.

### Glycolytic pathway flux quantitation via labeling in lower glycolysis metabolites

Glycolytic fluxes were obtained using the intracellular metabolite labeling from [1,2-^13^C_2_]-, [5,6-^13^C_2_]-, or [U-^13^C_6_]-glucose tracers (**Supplementary Tables 2-10**). [1,2-^13^C_2_]- and [5,6-^13^C_2_]-glucose tracers provided the necessary metabolite labeling (e.g., 3PG, 6PG, and Val) for determination of central carbon metabolism fluxes (**Supplementary Notes 2 and 3**). For the carbon upshift case, [U-^13^C_6_]-glucose tracer provided information necessary to correct for incomplete turnover of 3PG in early time points (**Supplementary Note 4**). Briefly, since the isotope labeling of 3PG from [1,2-^13^C_2_]glucose depends on which route glucose took (the EMP pathway, ED pathway, or pentose phosphate pathway), the following relationships between pathway fluxes and 3PG isotopologues were derived.

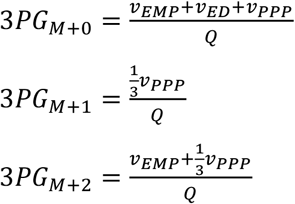

Q is the normalization factor that ensures the sum of 3PG mass isotopomer fractions is 1. These equations were rearranged to solve for the ED-to-EMP flux ratios in **Fig. 5**:

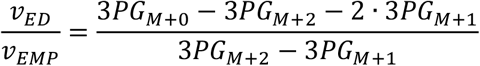

### Analysis of long-term evolution experiment

The long-term evolution experiment (LTEE) by Lenski et al. encompassed six cultures from each of the ancestral strains REL606 and REL607 (12 cultures total) in a DM medium with 25 mg/L glucose (citrate was also included in this medium as a chelating agent, which the ancestral strain does not grow on)^28^.

Cultures were diluted 1:100 into the fresh medium daily, and two clones from each culture were sampled periodically for sequencing. The mutation history of the 12 populations was downloaded from https://barricklab.org/shiny/LTEE-Ecoli/ and analyzed using MATLAB and Python programs.

Nonsynonymous (missense and nonsense) mutations and indels were considered for all analyses, and synonymous mutations were also included where stated (e.g., **Supplementary Figs. 15 and 17**).

The mutation history of the ED and the EMP pathways were compared to other pathways. Since the assignment of genes to metabolic pathways is not always one-to-one or clear-cut, pseudo pathways were formed from randomly generated groups of genes. Each pseudo pathway was generated by randomly selecting 10 genes from the *E. coli* genome. Mutations in the pathway genes were counted and normalized by the number of genes in the pathway at each generation. This process was repeated for 1,000 pathways as well as the EMP and the ED pathways, and the percentile ranks of the EMP and the ED pathways’ per-gene mutations were obtained. The EMP and the ED pathways included the following mutually exclusive sets of glycolytic genes: *pgi, pfka, pfkb, fbaa, fbab*, and *tpia* were included in the EMP pathway; and *zwf, pgl, edd*, and *eda* were included in the ED pathway. These gene sets were also used to track the number of strains with mutations in the EMP and the ED pathways.

To assess the randomness of the mutation history, mutation data from generation 50,000 were compared to the Poisson distribution. Without considering their identity, individual genes’ mutations were counted for all 24 clones. Since each of the populations begot two clones, which are thus not independent of each other, 2^12^ sets of 12 independent clones were generated using only one of the two sequenced clones from each population. Each set generated a distribution of frequencies of mutations in a gene. The mean and the standard deviation of these distributions over the 2^12^ sets were obtained. The resulting distribution was fit to a Poisson distribution using the MATLAB curve fitting toolbox (**Supplementary Fig.14**).

## Data availability

Source data for Figures 1-6 are provided in Supplementary Tables 1-17 and the GitHub public repository: https://github.com/richardlaw517/Parallel_Glycolysis

## Code availability

The code for the analysis of metabolic fluxes and LTEE is available on the GitHub public repository: https://github.com/richardlaw517/Parallel_Glycolysis

## Acknowledgements

The authors would like to thank the members of the Park lab, the UCLA Metabolomics Center, and the UCLA Molecular Instrumentation Center for helpful discussion. This work was supported by the National Institute Of General Medical Sciences of the National Institutes of Health under Award Number R35GM143127, the National Institutes of Health Instrumentation Grant Number 1S10OD016387-01, the BioPACIFIC Materials Innovation Platform of the National Science Foundation under Award Number DMR-1933487, and the Hellman Fellowship. The content is solely the responsibility of the authors and does not necessarily represent the official views of the National Institutes of Health or the National Science Foundation.

## Author Contributions

R.C.L. and J.O.P. designed the study and wrote the paper. R.C.L. carried out the experiments. R.C.L. and J.O.P. analyzed the metabolomic and isotope-labeling results. R.C.L., G.N., and J.O.P. analyzed the sequencing data from the long-term evolution experiment.

