## Supplemental Information for "A parallel glycolysis supports rapid adaptation in dynamic environments"

Supplementary Notes 1-4

Supplementary Figures 1-22

Supplementary Tables 1-17

Supplementary References

### Supplementary Notes

#### 1. Calculation of Gibbs free energies

Gibbs free energy of reaction (ΔG) is defined by the following equation.

$$\Delta G = \Delta G'^{\circ} + RT \ln Q$$

ΔG'° is ΔG at standard biochemical conditions, R is the universal gas constant, T is the temperature in kelvins, Q is the reaction quotient, i.e., the ratio of product-to-substrate activities, which are effective concentrations in a non-ideal solution.

##### *1a. Calculation of changes in Gibbs free energy of reaction from relative metabolite concentrations*

The difference in the Gibbs free energy between two metabolic states can be obtained using relative metabolite levels:

$$\Delta(\Delta G) = \left(\Delta G'^{\circ} + RT \ln Q\right)_2 - \left(\Delta G'^{\circ} + RT \ln Q\right)_1 = \Delta G'^{\circ} - \Delta G'^{\circ} + RT \ln \frac{Q_2}{Q_1} = RT \ln \frac{\left({P_2}/{P_1}\right)}{\left({R_2}/{R_1}\right)}$$

We computed the changes in thermodynamic driving force for the overall EMP and ED pathways using the following overall reactions.

EMP: G6P + ADP + 2 $NAD^+$ + 2 $P_i$ → 2 PEP + ATP + 2 NADH + 2 $H_2O$

ED: G6P + ADP + $NAD^+$ + $NADP^+$ + $P_i$ → PYR + PEP + ATP + NADH + NADPH + $H_2O$

ED + PpsA: G6P + ADP + $NAD^+$ + $NADP^+$ → 2 PEP + NADH + NADPH + AMP

GNG: 2 PEP + 2 ATP + 2 NADH + 3 $H_2O$ → G6P + 2 ADP + 2 $NAD^+$ + 3 $P_i$

In the first and the last step of glycolysis, the phosphotransferase system (PTS) carries out a reaction that involves both intracellular and extracellular metabolites (glucose). PTS imports glucose while phosphorylating it with phosphoenolpyruvate (PEP) to form G6P and pyruvate (PYR). Thus, we considered the pathway steps starting from G6P and ending with either PEP or PYR so that the reaction free energies are based solely on intracellular metabolite levels.

##### *1b. Calculation of Gibbs free energy of reaction from absolute metabolite concentrations*

Using absolute metabolite concentrations and ΔG'°, we obtained Gibbs free energy of reaction. We used the eQuilibrator[1] to calculate ΔG'° at physiological pH 7.7 and ionic strength 0.25 M[2].

The ΔG values in **Fig. 1b** were obtained using the following overall pathway reactions.

EMP: G6P + 2 $NAD^+$ + 3 ADP + 2 $P_i$ → 2 PYR + 2 NADH + 3 ATP + 2 $H_2O$

ED: G6P + $NAD^+$ + $NADP^+$ + 2 ADP + $P_i$ → 2 PYR + NADH + NADPH + 2 ATP + $H_2O$

It should be noted that these two lumped reactions included the pyruvate kinase (*pyk*) reaction for PEP-to-PYR conversion to highlight the parallel nature of the EMP and the ED glycolysis. However, this step is carried out mainly by the PTS, which transfers the phosphoryl group of PEP to glucose as part of glucose transport. When calculating ΔΔG in **Fig. 3**, we were interested in thermodynamic shifts as a result of intracellular metabolic rewiring, obviating the need for *pyk*.

### 2. Simple analytical solution to glycolytic flux distributions using the isotope labeling measurement of lower glycolytic intermediates

The EMP and the ED glycolytic fluxes were discerned by using an asymmetric $^{13}C$-glucose tracer. Depending on the glycolytic pathway taken, [1,2-$^{13}C_2$]glucose yields different labeling in lower glycolysis (**Fig. 4a**). The EMP glycolysis generates one-to-one ratio of labeled (M+2 or $M_2$) and unlabeled (M+0 or $M_0$) glyceraldehyde-3-phosphate (GAP) and subsequent lower glycolytic intermediates.

$$\text{EMP:} \quad 1\,M_0 \,\&\, 1\,M_2$$

The ED pathway generates one GAP using the unlabeled carbons (as well as one pyruvate using the labeled carbons).

$$\text{ED:} \quad 1\,M_0$$

Therefore, based on how far the $M_0$:$M_2$ ratio of GAP is from 50:50, the relative fluxes through the ED and the EMP pathways ($v_{ED}$ and $v_{EMP}$, respectively) can be obtained.

$$\frac{v_{ED}}{v_{EMP}} = \frac{M_0}{M_2} - 1$$

Using this equation, we obtained the ED-to-EMP flux ratio of 0.154 ± 0.006 in the nutrient replete condition. While it is a simple solution, it overestimates the ED pathway flux due to lack of consideration of the pentose phosphate pathway.

### 3. Analytical solution to fluxes through central carbon metabolism including the EMP pathway, the ED pathway, and the pentose phosphate pathway

Besides the EMP and the ED glycolysis, glucose can pass through the pentose phosphate pathway (PPP) in which three glucose molecules generate two fructose-6-phosphate (F6P) and one GAP (**Supplementary Fig. 6**).

$$\text{PPP:} \quad 3\,Glucose \rightarrow 2\,F6P + 1\,GAP$$

The resulting GAP is downstream of phosphofructokinase (PFK) reaction, a committed step of the EMP glycolysis, and is sent to lower glycolysis. However, F6P may either go to lower glycolysis via PFK, fructose-1,6-bisphosphate aldolase (FBA), and triose phosphate isomerase (TPI) or back to the ED pathway or the PPP due to the reversibility of phosphoglucoisomerase (PGI).

$$\text{R\_EMP:} F6P \rightarrow via\ PFK \,\&\, FBA \,\&\, TPI \rightarrow 2\,GAP$$

$$\text{R\_ED:} \quad F6P \rightarrow via\ reverse\ PGI \,\&\, ED \rightarrow PYR + GAP$$

$$\text{R\_PPP:} \ F6P \rightarrow via\ reverse\ PGI \,\&\, PPP \rightarrow \tfrac{2}{3}\ F6P + \tfrac{1}{3}\ GAP$$

Here we derive solutions to fluxes through central carbon metabolism by considering non-recursive (**Section 3a**) and recursive (**Section 3b**) usage of the PPP. We also utilized positional labeling of pyruvate from LC-MS/MS fragmentation of valine (**Section 3c**).

***3a. Case with no recycling of the PPP-derived F6P into the ED pathway or the PPP***

In the oxidative pentose phosphate pathway (OxPPP), the first carbon of [1,2-$^{13}C_2$]glucose is decarboxylated to generate singly labeled (M+1) pentose phosphate. Three hexose molecules generate three pentose phosphates, which become one doubly labeled (M+2) F6P, one singly labeled (M+1) F6P, and one unlabeled (M+0) GAP through the non-oxidative PPP. In the case where there is no recycling of F6P and all PPP-derived F6P molecules go through the EMP pathway, the resulting GAP labeling ratio M+0:M+1:M+2 becomes 3:1:1.

$$\text{PPP:} \quad \frac{3}{3}M_0,\ \frac{1}{3}M_1,\ \frac{1}{3}M_2$$

Here we mass balanced 3-phosphoglycerate (3PG) isotopomers because 3PG is reliably measured by LC-MS due to its abundance and it has the same isotope labeling as GAP because it is directly downstream of GAP. The fractions of the 3PG mass isotopomers resulting from [1,2-$^{13}C_2$]glucose are determined by the fluxes through the EMP pathway, the ED pathway, and the PPP.

$$3PG_{M+0} = \frac{v_{EMP}+v_{ED}+v_{PPP}}{Q} \qquad (1)$$

$$3PG_{M+1} = \frac{\frac{1}{3}v_{PPP}}{Q} \qquad (2)$$

$$3PG_{M+2} = \frac{v_{EMP}+\frac{1}{3}v_{PPP}}{Q} \qquad (3)$$

Q is the normalization factor that ensures the sum of 3PG mass isotopomer fractions to be 1.

$$Q = 2 \cdot v_{EMP} + v_{ED} + \frac{5}{3} \cdot v_{PPP}$$

We solved for the three pathway fluxes, $v_{EMP}$, $v_{ED}$, and $v_{PPP}$.

By rearranging **eqn. 2**, we obtained:

$$v_{PPP} = 3Q \cdot 3PG_{M+1}$$

By rearranging **eqn. 3** and using the solution for $v_{PPP}$, we obtained:

$$v_{EMP} = Q \cdot 3PG_{M+2} - \frac{1}{3}v_{PPP} = Q(3PG_{M+2} - 3PG_{M+1})$$

By rearranging **eqn. 1** and using the solutions for $v_{PPP}$ and $v_{EMP}$, we obtained:

$$v_{ED} = Q \cdot 3PG_{M+0} - v_{EMP} - v_{PPP} = Q(3PG_{M+0} - 3PG_{M+2} - 2 \cdot 3PG_{M+1})$$

Q is cancelled when we look at flux ratios. The flux ratio of the ED pathway to the EMP pathway is:

$$\frac{v_{ED}}{v_{EMP}} = \frac{3PG_{M+0} - 3PG_{M+2} - 2 \cdot 3PG_{M+1}}{3PG_{M+2} - 3PG_{M+1}}$$

We used this solution to calculate the ED and the EMP pathway fluxes in exponentially growing *E. coli* in replete media, finding a flux ratio of 0.089 ± 0.004.

***3b. Recycling of the PPP-derived F6P into the ED pathway or the PPP as informed by [5,6-$^{13}C_2$]glucose***

The OxPPP leads to [1,2-$^{13}C_2$]glucose generating singly labeled pentose phosphates and odd-numbered $^{13}$C-labeled pentose phosphates via non-OxPPP carbon shuffling. The complex labeling undermined our ability to quantify the extent of recycling of pentose phosphate pathway intermediates, thereby reducing the accuracy of the ED/EMP flux ratio calculation.

To circumvent this, we utilized [5,6-$^{13}C_2$]glucose, whose $^{13}$C atoms are neither lost nor shuffled by the PPP. Unlabeled (M+0) F6P arises only as a result of a transaldolase reaction that combines the first three (unlabeled) carbons of sedoheptulose-7-phosphate (S7P) and unlabeled GAP, which comes from the EMP pathway. As a result, the presence of M+0 OxPPP intermediates (e.g., 6-phosphogluconate (6PG)) implies the recycling of PPP-derived F6P. Therefore, we obtain the extent of recycling (C) using the unlabeled fraction of 6PG and use the C to refine the ED/EMP flux ratio.

<u>Derivation of the relationship between C, fluxes, and unlabeled metabolite fractions from [5,6-$^{13}C_2$]glucose:</u>

Recycle factor (C) is the fraction of all 6PG generated that comes from hexose phosphates that are derived from the PPP intermediates. This is achieved by multiplying the fraction of G6P derived by reverse PGI by the fraction of F6P derived from PPP flux (**Supplementary Fig. 22**):

$$C = \left(\frac{\text{6PG from G6PDH}}{\text{All incoming 6PG}}\right) \cdot \left(\frac{\text{6PG from reverse pgi}}{\text{All incoming G6P}}\right) \cdot \left(\frac{\text{F6P from PPP}}{\text{All incoming F6P}}\right)$$

$$= 1 \cdot \left(\frac{v_{reverse_{pgi}}}{v_{glc\_uptake} + v_{reverse_{pgi}}}\right) \cdot \left(\frac{\frac{2}{3} v_{PPP}}{v_{forward_{pgi}} + \frac{2}{3} v_{PPP}}\right)$$

A balance on G6P can be used to define C in terms of only intracellular fluxes:

$$\frac{dG6P}{dt} = 0 = v_{glc\_uptake} + v_{reverse_{pgi}} - v_{g6pd} - v_{forward_{pgi}}$$

$$C = \left(\frac{v_{reverse_{pgi}}}{v_{forward_{pgi}} + v_{g6pd}}\right) \cdot \left(\frac{\frac{2}{3} v_{PPP}}{v_{forward_{pgi}} + \frac{2}{3} v_{PPP}}\right)$$

The recycle factor is 0 when there is no reverse PGI flux. On the extreme with no forward PGI flux, the recycle factor reaches a maximum of $^{2}/_{3}$. To solve for C between these two extremes, we first balance M+0 6PG under steady state:

$$\frac{d6PG_{M+0}}{dt} = 0 = v_{g6pd} \cdot G6P_{M+0} - (v_{PPP} + v_{ED}) \cdot 6PG_{M+0} = v_{g6pd} \cdot G6P_{M+0} - v_{g6pd} \cdot 6PG_{M+0}$$

$$6PG_{M+0} = G6P_{M+0}$$

Furthermore, balancing M+0 G6P yields:

$$\frac{dG6P_{M+0}}{dt} = 0 = v_{reverse_{pgi}} \cdot F6P_{M+0} - v_{forward_{pgi}} \cdot G6P_{M+0} - v_{g6pd} \cdot G6P_{M+0}$$

$$G6P_{M+0} = \frac{v_{reverse_{pgi}} \cdot F6P_{M+0}}{v_{forward_{pgi}} + v_{g6pd}} = 6PG_{M+0}$$

M+0 F6P is generated from the portion of PPP flux that uses unlabeled GAP and unlabeled 6PG as well as the portion of forward PGI flux that uses unlabeled G6P:

$$\frac{dF6P_{M+0}}{dt} = 0 = v_{forward_{pgi}} \cdot G6P_{M+0} + \frac{1}{3}v_{PPP} \cdot 6PG_{M+0} + \frac{1}{3}v_{PPP} \cdot GAP_{M+0} - v_{reverse_{pgi}} \cdot F6P_{M+0} - v_{pfk} \cdot F6P_{M+0}$$

Balancing the whole F6P metabolite pool yields:

$$\frac{dF6P}{dt} = 0 = v_{forward_{pgi}} - v_{reverse_{pgi}} + \frac{2}{3}v_{PPP} - v_{pfk} = v_{EMP} + \frac{2}{3}v_{PPP} - v_{pfk}$$

The whole F6P balance can be used to simplify the M+0 F6P balance to:

$$0 = v_{forward_{pgi}} \cdot 6PG_{M+0} + \frac{1}{3}v_{PPP} \cdot 6PG_{M+0} + \frac{1}{3}v_{PPP} \cdot GAP_{M+0} - \left(v_{forward_{pgi}} + \frac{2}{3}v_{PPP}\right) \cdot F6P_{M+0}$$

$$\left(v_{forward_{pgi}} + \frac{2}{3}v_{PPP}\right) \cdot F6P_{M+0} = v_{forward_{pgi}} \cdot 6PG_{M+0} + \frac{1}{3}v_{PPP} \cdot 6PG_{M+0} + \frac{1}{3}v_{PPP} \cdot GAP_{M+0}$$

$$F6P_{M+0} = \frac{v_{forward_{pgi}} \cdot 6PG_{M+0} + \frac{1}{3}v_{PPP} \cdot 6PG_{M+0} + \frac{1}{3}v_{PPP} \cdot GAP_{M+0}}{v_{forward_{pgi}} + \frac{2}{3}v_{PPP}}$$

Inputting this solution for M+0 F6P into the M+0 6PG balance yields:

$$6PG_{M+0} = \left(\frac{v_{reverse_{pgi}}}{v_{forward_{pgi}} + v_{g6pd}}\right) \cdot \left(\frac{v_{forward_{pgi}} \cdot 6PG_{M+0} + \frac{1}{3}v_{PPP} \cdot 6PG_{M+0} + \frac{1}{3}v_{PPP} \cdot GAP_{M+0}}{v_{forward_{pgi}} + \frac{2}{3}v_{PPP}}\right)$$

We can then rearrange this equation to solve for C:

$$6PG_{M+0} = \left(\frac{v_{reverse_{pgi}}}{v_{forward_{pgi}} + v_{g6pd}}\right) \cdot \left(\frac{\frac{2}{3}v_{PPP}}{\frac{2}{3}v_{PPP}}\right) \cdot \left(\frac{v_{forward_{pgi}} \cdot 6PG_{M+0} + \frac{1}{3}v_{PPP} \cdot (6PG_{M+0} + GAP_{M+0})}{v_{forward_{pgi}} + \frac{2}{3}v_{PPP}}\right)$$

$$6PG_{M+0} = C \cdot \left(\frac{v_{forward_{pgi}} \cdot 6PG_{M+0} + \frac{1}{3}v_{PPP} \cdot (6PG_{M+0} + GAP_{M+0})}{\frac{2}{3}v_{PPP}}\right)$$

$$C = \frac{2 \cdot 6PG_{M+0}}{3 \cdot \frac{v_{forward_{pgi}}}{v_{PPP}} \cdot 6PG_{M+0} + 6PG_{M+0} + 3PG_{M+0}}$$

where we have taken M+0 GAP to be equal to M+0 3PG.

<u>Estimating the forward PGI to PPP flux ratio by solving balances on upper and whole glycolysis:</u>

To solve for C, the flux ratio between the forward PGI and PPP is needed. The forward PGI flux depends on the reversibility of the PGI reaction. Here we used the relationship between the net and forward fluxes, which states that the forward flux is always greater than or equal to the net flux. Therefore, the flux ratio satisfies the following inequality:

$$\frac{v_{EMP}}{v_{PPP}} \leq \frac{v_{forward_{pgi}}}{v_{PPP}}$$

We solved for the flux ratio between the EMP pathway and the PPP by first determining the fraction of M+0 GAP and 3PG through a carbon balance on upper glycolysis and the PPP (**Supplementary Fig. 8**):

$$GAP_{M+0} = \frac{1 - v_{ED} - \frac{1}{3} \cdot v_{PPP}}{2 - v_{ED} - \frac{1}{3} \cdot v_{PPP}} = \frac{3 - 3 \cdot v_{ED} - v_{PPP}}{6 - 3 \cdot v_{ED} - v_{PPP}} = 3PG_{M+0}$$

We performed a carbon balance also with lower glycolysis to the control volume to determine the fraction of M+0 pyruvate:

$$PYR_{M+0} = \frac{v_{EMP} + v_{ED} + \frac{2}{3} \cdot v_{PPP}}{2 \cdot v_{EMP} + 2 \cdot v_{ED} + \frac{5}{3} \cdot v_{PPP}} = \frac{1 - \frac{1}{3} \cdot v_{PPP}}{2 - \frac{1}{3} \cdot v_{PPP}} = \frac{3 - v_{PPP}}{6 - v_{PPP}}$$

These two balances can be used to solve for central carbon metabolism fluxes. For the PPP, we get:

$$v_{PPP} = \frac{3 - 6 \cdot PYR_{M+0}}{1 - PYR_{M+0}}$$

To solve for ED flux, we begin with the solution for M+0 3PG:

$$3PG_{M+0} = \frac{3 - 3 \cdot v_{ED} - v_{PPP}}{6 - 3 \cdot v_{ED} - v_{PPP}}$$

Multiplying both sides by the denominator:

$$6 \cdot 3PG_{M+0} - 3 \cdot v_{ED} \cdot 3PG_{M+0} - v_{PPP} \cdot 3PG_{M+0} = 3 - 3 \cdot v_{ED} - v_{PPP}$$

And subsequently isolating ED flux and inputting the solution to the PPP flux:

$$3 \cdot v_{ED} \cdot (1 - 3PG_{M+0}) = 3 - 6 \cdot 3PG_{M+0} - v_{PPP} \cdot (1 - 3PG_{M+0})$$

$$v_{ED} = \frac{3 - 6 \cdot 3PG_{M+0} - v_{PPP} \cdot (1 - 3PG_{M+0})}{3 \cdot (1 - 3PG_{M+0})}$$

$$v_{ED} = \frac{3 - 6 \cdot 3PG_{M+0} - \left(\frac{3 - 6 \cdot PYR_{M+0}}{1 - PYR_{M+0}}\right) \cdot (1 - 3PG_{M+0})}{3 \cdot (1 - 3PG_{M+0})}$$

Given these two solutions for PPP flux and ED flux and normalizing all fluxes to glucose uptake rate (i.e., $v_{glc\_uptake}$=1), we can solve for EMP flux:

$$v_{EMP} = 1 - v_{ED} - v_{PPP} = 1 - \frac{3 - 6 \cdot 3PG_{M+0} - \left(\frac{3 - 6 \cdot PYR_{M+0}}{1 - PYR_{M+0}}\right) \cdot (1 - 3PG_{M+0})}{3 \cdot (1 - 3PG_{M+0})} - \frac{3 - 6 \cdot PYR_{M+0}}{1 - PYR_{M+0}}$$

From these solutions to glycolytic fluxes, we obtained the flux ratio by inputting formulas for the EMP and the PPP fluxes:

$$\frac{v_{EMP}}{v_{PPP}} = \frac{1 - \frac{3 - 6 \cdot 3PG_{M+0} - \left(\frac{3 - 6 \cdot PYR_{M+0}}{1 - PYR_{M+0}}\right) \cdot (1 - 3PG_{M+0})}{3 \cdot (1 - 3PG_{M+0})} - \frac{3 - 6 \cdot PYR_{M+0}}{1 - PYR_{M+0}}}{\frac{3 - 6 \cdot PYR_{M+0}}{1 - PYR_{M+0}}}$$

We simplified the fractions by canceling out the lowest denominator where appropriate:

$$\frac{v_{EMP}}{v_{PPP}} = \frac{1 - PYR_{M+0}}{3 - 6 \cdot 3PG_{M+0}} - \frac{\frac{(3 - 6 \cdot 3PG_{M+0}) \cdot (1 - PYR_{M+0})}{3 - 6 \cdot PYR_{M+0}} - (1 - 3PG_{M+0})}{3 \cdot (1 - 3PG_{M+0})} - 1$$

$$\frac{v_{EMP}}{v_{PPP}} = \frac{1 - PYR_{M+0}}{3 - 6 \cdot 3PG_{M+0}} - \frac{(3 - 6 \cdot 3PG_{M+0}) \cdot (1 - PYR_{M+0})}{3 \cdot (3 - 6 \cdot PYR_{M+0}) \cdot (1 - 3PG_{M+0})} - \frac{2}{3} \leq \frac{v_{forward_{pgi}}}{v_{PPP}}$$

This expression gives us a lower bound on the ratio of forward PGI to PPP flux. We take this ratio to be equal to its lower bound so as to get an upper bound for the recycle factor C. This leads to a solution considering the highest possible recycling:

$$C_{max} = \frac{2 \cdot 6PG_{M+0}}{3 \cdot \frac{v_{forward_{pgi}}}{v_{PPP}} \cdot 6PG_{M+0} + 6PG_{M+0} + 3PG_{M+0}}$$

where $\frac{v_{forward_{pgi}}}{v_{PPP}} = \frac{1-PYR_{M+0}}{3-6 \cdot 3PG_{M+0}} - \frac{(3-6 \cdot 3PG_{M+0}) \cdot (1-PYR_{M+0})}{3 \cdot (3-6 \cdot PYR_{M+0}) \cdot (1-3PG_{M+0})} - \frac{2}{3}$

In the nutrient replete condition, we used labeling data for 3PG, 6PG, and pyruvate and found $C_{max} = 0.088 \pm 0.007$.

Utilizing C to estimate pentose phosphate pathway isotopomers from [1,2-$^{13}C_2$]glucose tracing:

Returning to [1,2-$^{13}C_2$]glucose tracing now including the recycle factor $C=C_{max}$, we consider that a fraction of the PPP flux (i.e., 1-C) directly generates the original isotopomers in **Section 3a** while the rest (i.e., C) generates complex labeling forms from the recursive use of the PPP.

$$3PG_{M+0} = \frac{v_{EMP}([G6P_{000}] + (1 - C) + C \cdot 3PG_{M+0}) + v_{ED}\left((1 - C) + C \cdot 3PG_{M+0}\right) + v_{PPP}\left((1 - C) + C \cdot 3PG_{M+0} + \frac{2C}{3} \cdot [6PG_{000} + 6PG_{100}]\right)}{Q}$$

$$3PG_{M+1} = \frac{v_{EMP}([G6P_{100} + G6P_{010} + G6P_{001}] + C \cdot 3PG_{M+1}) + v_{ED}(C \cdot 3PG_{M+1}) + v_{PPP}\left(\frac{1}{3}(1 - C) + C \cdot 3PG_{M+1} + \frac{2C}{3} \cdot \left[\frac{6PG_{010} + 6PG_{001} + 6PG_{110} + 6PG_{101}}{2}\right]\right)}{Q}$$

$$3PG_{M+2} = \frac{v_{EMP}([G6P_{011} + G6P_{101} + G6P_{110}] + C \cdot 3PG_{M+2}) + v_{ED}(C \cdot 3PG_{M+2}) + v_{PPP}\left(\frac{1}{3}(1 - C) + C \cdot 3PG_{M+2} + \frac{2C}{3} \cdot \left[\frac{6PG_{010} + 6PG_{001} + 6PG_{110} + 6PG_{101}}{2}\right]\right)}{Q}$$

$$3PG_{M+3} = \frac{v_{EMP}([G6P_{111}] + C \cdot 3PG_{M+3}) + v_{ED}(C \cdot 3PG_{M+3}) + v_{PPP}\left(C \cdot 3PG_{M+3} + \frac{2C}{3} \cdot [6PG_{011} + 6PG_{111}]\right)}{Q}$$

where $G6P_{xxx}$ and $6PG_{xxx}$ are the simulated fraction for the labeling state of the first three carbons in G6P and 6PG (x=0 means unlabeled and x=1 means labeled for each of carbon position) and Q is a normalization that ensures the isotopomer fractions sum up to 1. These isotopomer fractions were generated by iteratively simulating atom transitions in the PPP (**Supplementary Fig. 6**) to get the labeling of the product F6P from [1,2-$^{13}C_2$]glucose. The distribution of mass isotopomers for F6P and G6P are represented as an array list of vectors describing the binary labeling state of individual atoms. Based on the recycle factor C, a proportion of these vectors from the F6P vector pool is added to the pool for G6P, which can then be used for the next iteration of PPP simulation. Successive iteration (>1,000) of this process ensures the stochastic nature by which hexose phosphates are chosen for the PPP and thus simulates all possible isotopomers generated by the pentose phosphate pathway after a sustained period of activity (i.e., a steady state).

We solved for 3PG labeling in terms of simulated hexose isotopomers and fluxes.

$$3PG_{M+0} = \frac{v_{EMP} \cdot ([G6P_{000}] + 1 - C) + v_{ED} \cdot (1 - C) + v_{PPP} \cdot \left(1 - C + \frac{2C}{3} \cdot [6PG_{000} + 6PG_{100}]\right)}{Q - C}$$

$$3PG_{M+1} = \frac{v_{EMP} \cdot [G6P_{100} + G6P_{010} + G6P_{001}] + \frac{v_{PPP}}{3} \cdot \left(1 - C + 2C \cdot \left[\frac{6PG_{010} + 6PG_{001} + 6PG_{110} + 6PG_{101}}{2}\right]\right)}{Q - C}$$

$$3PG_{M+2} = \frac{v_{EMP} \cdot [G6P_{011} + G6P_{101} + G6P_{110}] + \frac{v_{PPP}}{3} \cdot \left(1 - C + 2C \cdot \left[\frac{6PG_{010} + 6PG_{001} + 6PG_{110} + 6PG_{101}}{2}\right]\right)}{Q - C}$$

$$3PG_{M+3} = \frac{v_{EMP} \cdot [G6P_{111}] + \frac{2C}{3} v_{PPP} \cdot [6PG_{011} + 6PG_{111}]}{Q - C}$$

Using these equations and the labeling data for 3PG, the simulated labeling for hexose phosphates, and $C_{max}$, we obtained values for the three glycolytic fluxes by mathematical optimization that minimized the variance-weighted sum of squared residuals between the simulated and measured labeling isotopomer abundances of 3PG.

$$\min_{v} \sum_{n=0}^{3} \left( \frac{3PG_{obs,M+n} - 3PG_{M+n}(v)}{SD_{3PG_{M+n}}} \right)^2$$

subject to $v_{EMP} \in [-2,1]$, $v_{ED}, v_{PPP} \in [0,3]$, and $v_{EMP} + v_{ED} + v_{PPP} = 1$

$v$ is the vector of glycolytic fluxes, $3PG_{obs}$ is the experimentally observed mass isotopomer distributions, 3PG($v$) is the labeling simulated by solving each expression for the 3PG isotopomers given $v$, and $SD_{MID}$ is the standard deviation of the measurements. M+n denotes the number of carbons labeled with $^{13}C$ in 3PG. This process was iterated 1,000 times to find the global minimum. Using this approach, we found the normalized EMP and ED fluxes to be 0.895 ± 0.007, 0.105 ± 0.007. These obtained fluxes were used to calculate the ED-to-EMP flux ratio: 0.118 ± 0.009.

#### *3c. Validation of the fluxes by MS/MS fragmentation of valine*

We further validated our flux results using the positional labeling information obtained from MS/MS fragmentation of valine, which is synthesized by combining two pyruvate molecules (**Fig. 4b**, **Supplementary Fig. 5**). The ED pathway, EMP pathway, and the PPP produce pyruvate isotopologues that contain $^{13}C$ in different positions (**Fig. 4a, Supplementary Fig. 6**).

We mapped out all possible isotopomers pyruvate and noted that there are three possible isotopomers of M+2 pyruvate can come from the three metabolic pathways:

$$PYR_{M+2} = x + y + z$$

x represents the fraction of [1,2-$^{13}C_2$]pyruvate derived from the ED pathway, y represents the fraction of [2,3-$^{13}C_2$]pyruvate derived from the EMP glycolysis, and z represents the fraction of [1,3-$^{13}C_2$]pyruvate derived from the PPP. We can connect glycolytic fluxes to x, y, and z by the stoichiometry in which these pathways create the isotopologues of pyruvate.

$$PYR_{M+2} = \frac{v_{ED} + v_{EMP} + \frac{1}{3} v_{PPP}}{Q} = x + y + z$$

$$x = \frac{v_{ED}}{Q} \qquad y = \frac{v_{EMP}}{Q} \qquad z = \frac{v_{PPP}}{3Q}$$

Q is a normalization factor to ensure the sum of all pyruvate isotopomers (i.e., $PYR_{M+0}$ + $PYR_{M+1}$ + $PYR_{M+2}$ + $PYR_{M+3}$) equals to 1 and cancels out when solving for the ED-to-EMP flux ratio.

Including M+0 and M+1 as well as the three M+2 forms, pyruvate has five possible isotopologues. Because valine is synthesized from two pyruvate molecules, there are a total of 25 possible isotopologues of valine (**Supplementary Fig. 5**). These resulting isotopologues generate distinct parent-fragment ion pairs when run through LC-MS/MS, which decarboxylates the first carbon of valine during fragmentation. The proportion of each possible isotopologue observed reflects the labeling fractions of the precursor pyruvates:

$$Val_{M+0,M+0} = (PYR_{M+0}) \cdot (PYR_{M+0})$$

$$Val_{M+1,M+1} = (PYR_{M+0}) \cdot (x) + (PYR_{M+0}) \cdot (z) + 2 \cdot (PYR_{M+0}) \cdot (PYR_{M+1})$$

$$Val_{M+2,M+2} = 2 \cdot (y) \cdot (PYR_{M+0}) + (PYR_{M+1}) \cdot (x) + (PYR_{M+1}) \cdot (z) + (PYR_{M+1}) \cdot (PYR_{M+1})$$

$$Val_{M+2,M+1} = (x) \cdot (PYR_{M+0}) + (z) \cdot (PYR_{M+0})$$

$$Val_{M+3,M+3} = (y) \cdot (x) + (y) \cdot (z) + 2 \cdot (y) \cdot (PYR_{M+1})$$

$$Val_{M+3,M+2} = (x) \cdot (x) + 2 \cdot (x) \cdot (z) + (x) \cdot (PYR_{M+1}) + (z) \cdot (PYR_{M+1}) + (z) \cdot (z)$$

$$Val_{M+4,M+4} = (y) \cdot (y)$$

$$Val_{M+4,M+3} = (x) \cdot (y) + (z) \cdot (y)$$

where the first subscript of Val refers to the isotopologue of the parent valine ion and the second subscript refers to the isotopologue of the resulting fragment. We obtained the values for x, y, and z by mathematical optimization that minimized the variance-weighted sum of squared residuals between all the simulated and measured valine parent-fragment ion pair abundances and pyruvate isotopomer abundances.

$$\min_{x,y,z,PYR_{M+0},PYR_{M+1}} \sum_{l=0}^{l=5} \sum_{n=0}^{n=4} \left( \frac{Val_{obs,M+l,M+n} - Val_{M+l,M+n}(x,y,z,PYR_{M+0},PYR_{M+1})}{SD_{Val_{M+l,M+n}}} \right)^2 + \sum_{n=0}^{3} \left( \frac{PYR_{obs,M+n} - PYR_{M+n}(x,y,z,PYR_{M+0},PYR_{M+1})}{SD_{PYR_{M+n}}} \right)^2$$

subject to $x, y, z, PYR_{M+0}, PYR_{M+1} \in [0,1]$, and $x + y + z + PYR_{M+0} + PYR_{M+1} = 1$

$Val_{obs}$ is the experimentally observed fraction of each valine parent-fragment pair, Val(x,y,z,$PYR_{M+0}$,$PYR_{M+1}$) is the simulated fraction by solving each expression for the valine isotopologues, and $SD_{Val}$ is the standard deviation of the measurements. l and n denote the number of $^{13}C$ of the valine parent (M+l) and fragment

(M+n) ions, respectively. $PYR_{obs}$ is the experimentally observed fraction of pyruvate isotopologue, $PYR(x,y,z,PYR_{M+0},PYR_{M+1})$ is the simulated fraction, and $SD_{PYR}$ is the standard deviation of the measurements. M+n denotes number of carbons labeled in pyruvate. To obtain the global minimum, this process was iterated 1,000 times with different initial values for x, y, and z. Using this approach, we found *E. coli* growing exponentially on a replete medium to have an x, y, and z of 0.023 ± 0.003, 0.399 ± 0.001, and 0.003 ± 0.003, respectively. The ED-to-EMP flux ratio was solved by taking dividing x by y, yielding a ratio of 0.059 ± 0.008.

**4. Correction of isotopic labeling data for pseudo-steady-state labeling in carbon upshift**

In acetate to glucose carbon upshift, the metabolite pools of glycolytic intermediates were incompletely labeled even though they reached stable pseudo-steady state labeling. Some residual acetate-sourced metabolites in lower glycolysis remained unlabeled and were uninformative for computing glycolytic fluxes (**Supplementary Fig. 13**). We corrected for this incomplete labeling by considering the total glycolysis-sourced pool of a metabolite to be the fraction of labeled metabolite from the [U-$^{13}C_6$]glucose upshift experiment at a given time. The total pool of glucose-sourced 3PG is the fraction of its carbons that are $^{13}C$ labeled:

$$3PG_{Glc} = \frac{0 \cdot 3PG_{M+0,U} + 1 \cdot 3PG_{M+1,U} + 2 \cdot 3PG_{M+2,U} + 3 \cdot 3PG_{M+3,U}}{3}$$

The corrected fractions are:

$$3PG_{M+0,A,corrected} = \frac{3PG_{M+0,A} - (1 - 3PG_{Glc})}{3PG_{Glc}}$$

$$3PG_{M+1,A,corrected} = \frac{3PG_{M+1,A}}{3PG_{Glc}}$$

$$3PG_{M+2,A,corrected} = \frac{3PG_{M+2,A}}{3PG_{Glc}}$$

$$3PG_{M+3,A,corrected} = \frac{3PG_{M+3,A}}{3PG_{Glc}}$$

The subscripts U and A denote the uniformly labeled and asymmetrically labeled glucose experiments, respectively.

Our two asymmetric tracers (i.e., [1,2-$^{13}C_2$]- and [5,6-$^{13}C_2$]-glucose) did not yield 'mirror-image' labeling. The M+0 labeling of 3PG from [1,2-$^{13}C_2$]glucose is analogous to M+2 3PG from [5,6-$^{13}C_2$]glucose; however, these labeling fractions disagreed. We attributed this difference to possible residual reverse flux through lower glycolysis, resulting in the labeling of 3PG to be partially impacted by that of pyruvate. The effect of the reverse glycolytic flux from pyruvate to 3PG cancels out when we average the mirror-image 3PG products from the [1,2-$^{13}C_2$]- and [5,6-$^{13}C_2$]-glucose tracing experiments. As a result, the further corrected fractions from [1,2-$^{13}C_2$]glucose (denoted by $*$) are:

$$3PG_{M+0,[1,2],corrected*} = \frac{3PG_{M+0,[1,2],corrected} + 3PG_{M+2,[5,6],corrected}}{2}$$

$$3PG_{M+1,[1,2],corrected*} = \frac{3PG_{M+1,[1,2],corrected} + 3PG_{M+2,[1,2],corrected} + 3PG_{M+0,[5,6],corrected}}{2} \cdot \left(\frac{3PG_{M+1,[1,2],corrected}}{3PG_{M+1,[1,2],corrected} + 3PG_{M+2,[1,2],corrected}}\right)$$

$$3PG_{M+2,[1,2],corrected*} = \frac{3PG_{M+1,[1,2],corrected} + 3PG_{M+2,[1,2],corrected} + 3PG_{M+0,[5,6],corrected}}{2} \cdot \left(\frac{3PG_{M+2,[1,2],corrected}}{3PG_{M+1,[1,2],corrected} + 3PG_{M+2,[1,2],corrected}}\right)$$

$$3PG_{M+3,[1,2],corrected*} = \frac{3PG_{M+3,[1,2],corrected} + 3PG_{M+3,[5,6],corrected}}{2}$$

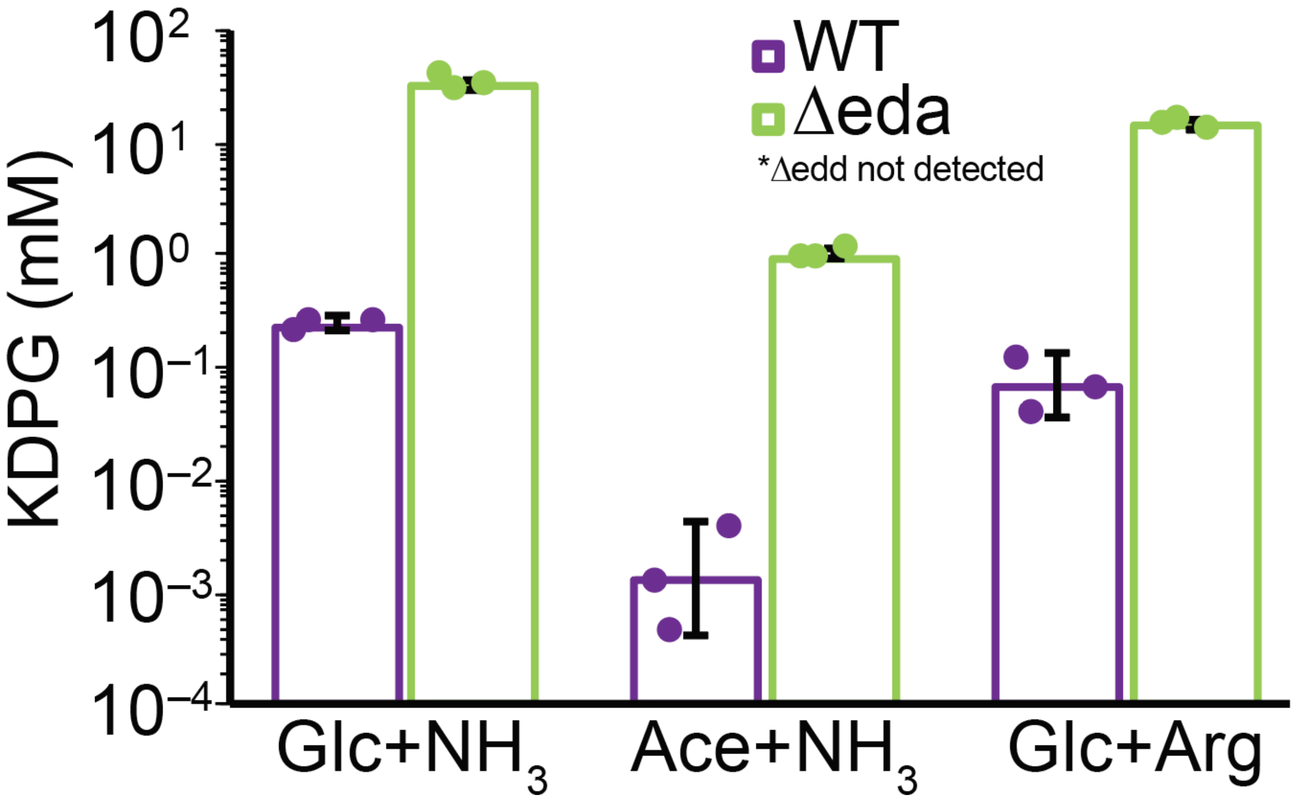


**Supplementary Figure 1. 2-keto-3-deoxy-6-phosphogluconate (KDPG) accumulates in Δ*eda*.** KDPG, a unique intermediate of the ED pathway, accumulated to high concentrations in the Δ*eda* strain. The absolute KDPG concentrations were determined using an isotope labeling ratio approach with an authenticated KDPG standard. Error bars represent the s.e.m. (3 biological replicates).

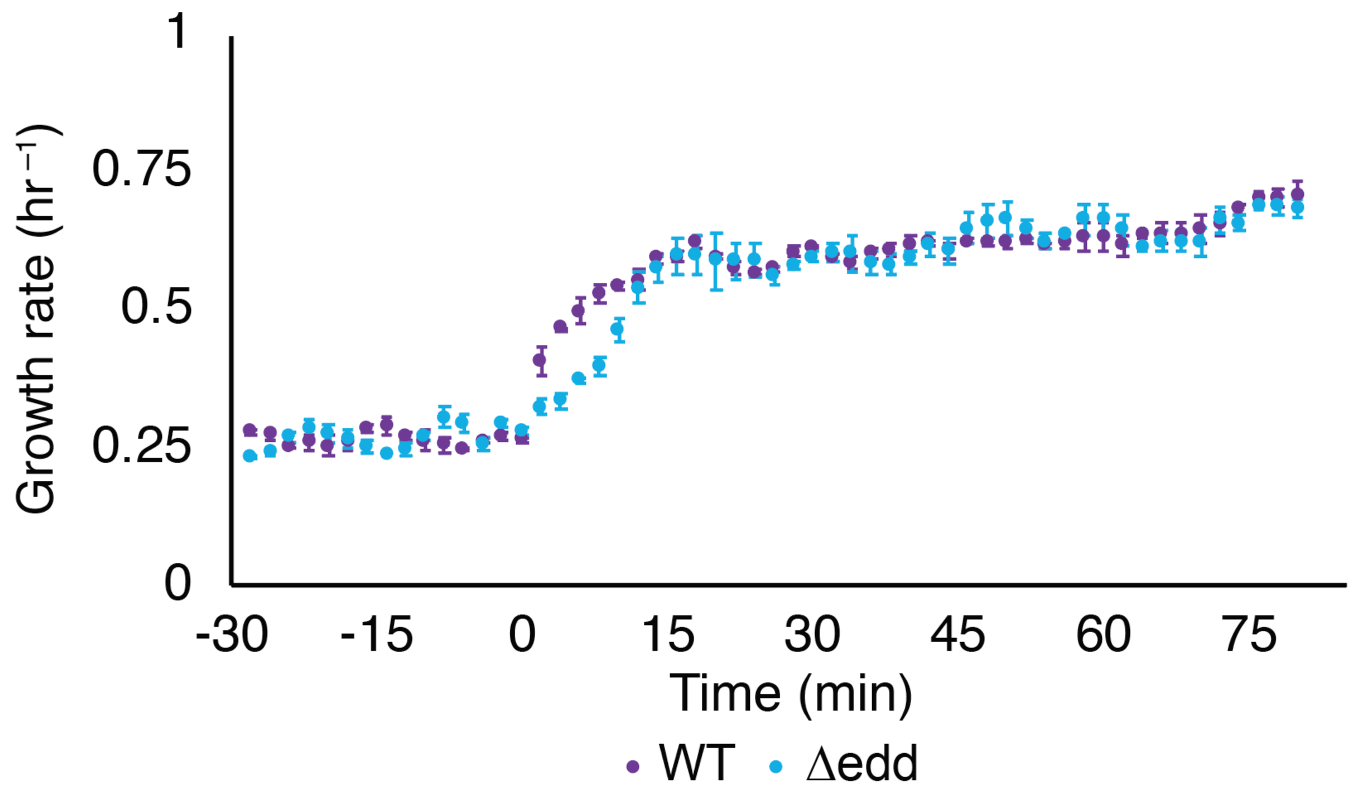


**Supplementary Figure 2.** ***E. coli*** **growth rate recovers after carbon upshift.** WT and Δ*edd* strains were cultured on acetate and spiked with glucose at t=0 min. Cells initially rapidly accelerated growth before proceeding to slowly increase its growth rate over the next hour. A steady growth rate of ~0.7 $hr^{-1}$ was achieved 80 minutes after upshift. Error bars represent the s.e.m. (n=3 biological replicates).

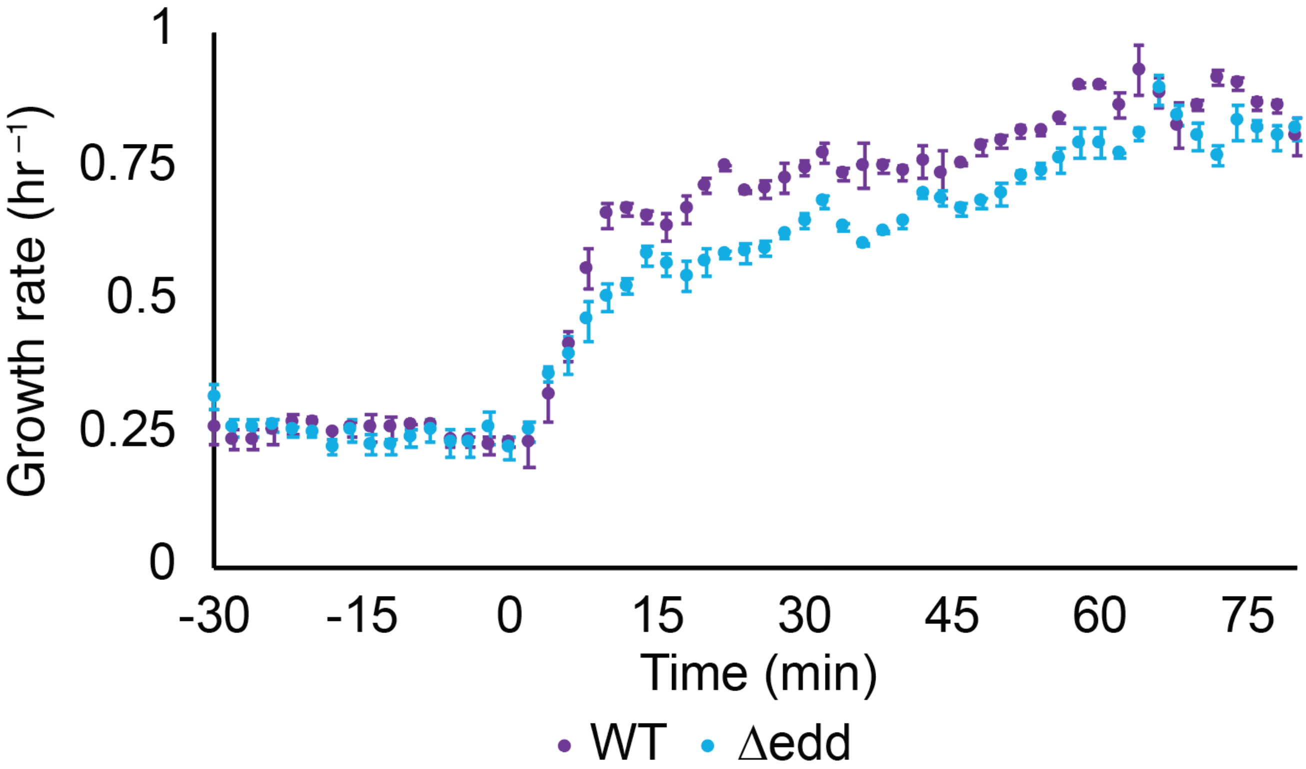


**Supplementary Figure 3. *E. coli* growth recovers after nitrogen upshift.** WT and Δ*edd* strains were cultured on arginine before ammonia was spiked in at t=0 min. Cells rapidly accelerated growth in the first 15 minutes of nitrogen upshift, and both reached a stable growth rate of ~0.75 $hr^{-1}$ 80 minutes after upshift. Error bars represent the s.e.m. (n=3 biological replicates).

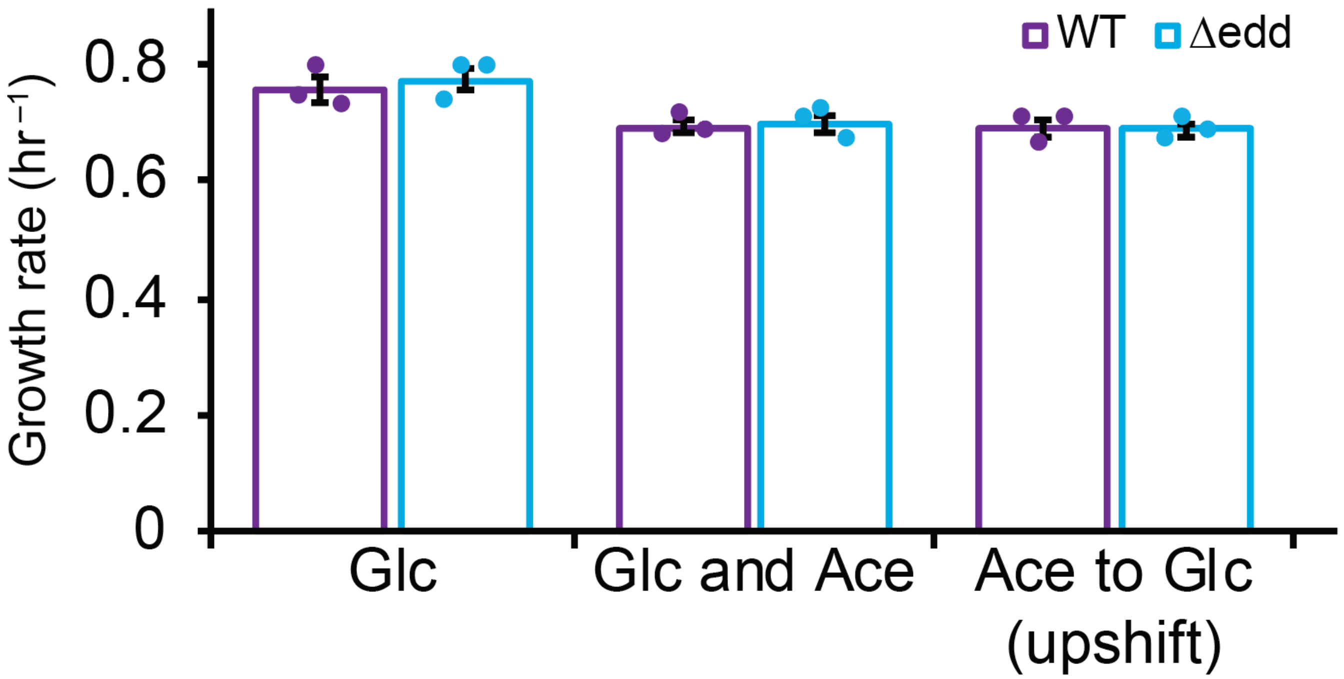


**Supplementary Figure 4. *E. coli* growth is diminished in high acetate cultures.** *E. coli* grow slower on glucose and acetate than on glucose alone. *E. coli* following glucose upshift has a similar stable growth rate to glucose and acetate cultures after upshift. Error bars represent the s.e.m. (n=3 biological replicates).

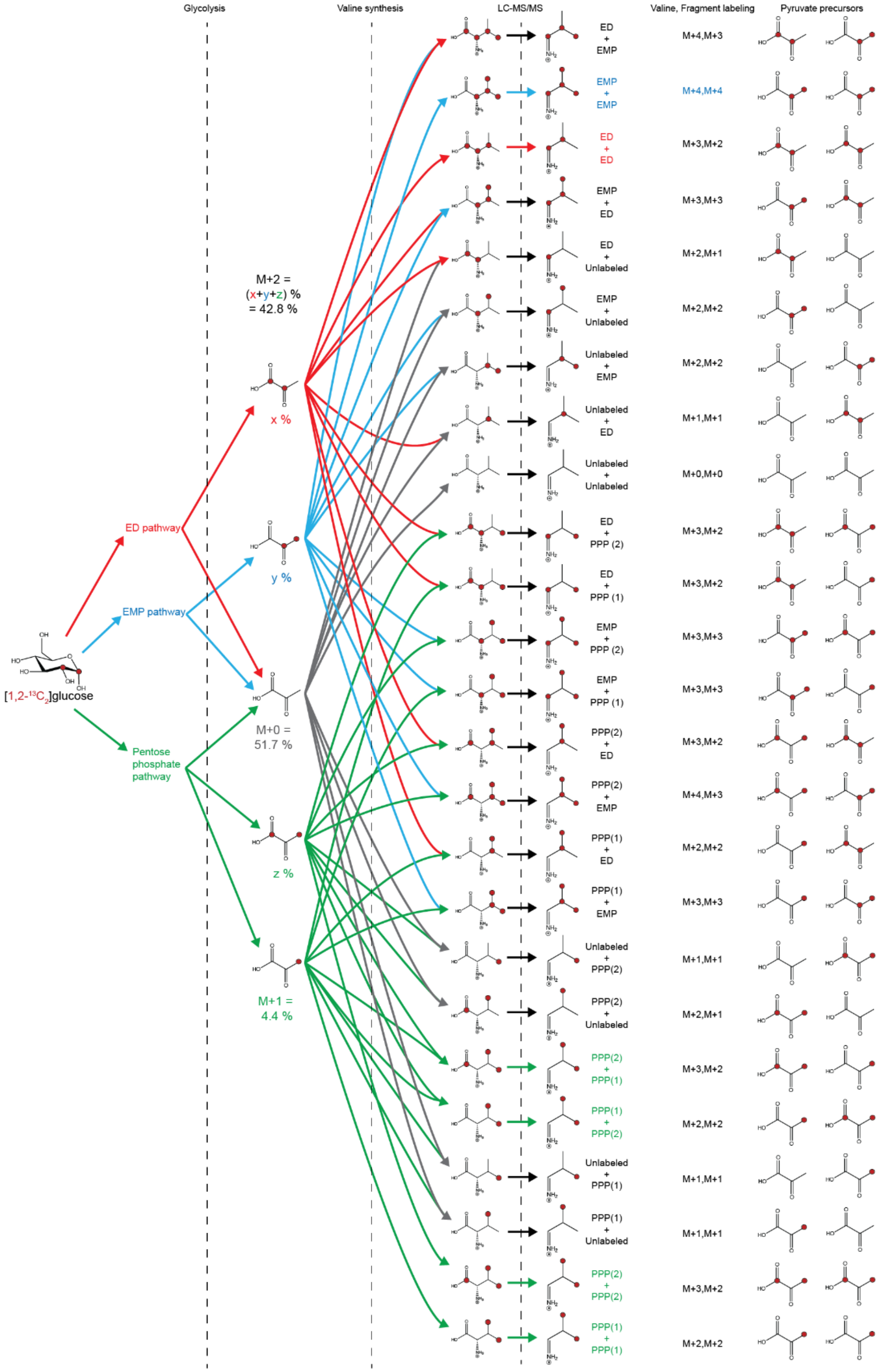


**Supplementary Figure 5. Valine isotopologues reveal glycolytic fluxes.** Using [1,2-$^{13}C_2$]glucose, 25 isotopologues of valine are produced. The information on the positional labeling of valine revealed the EMP pathway, the ED pathway, and the PPP fluxes.

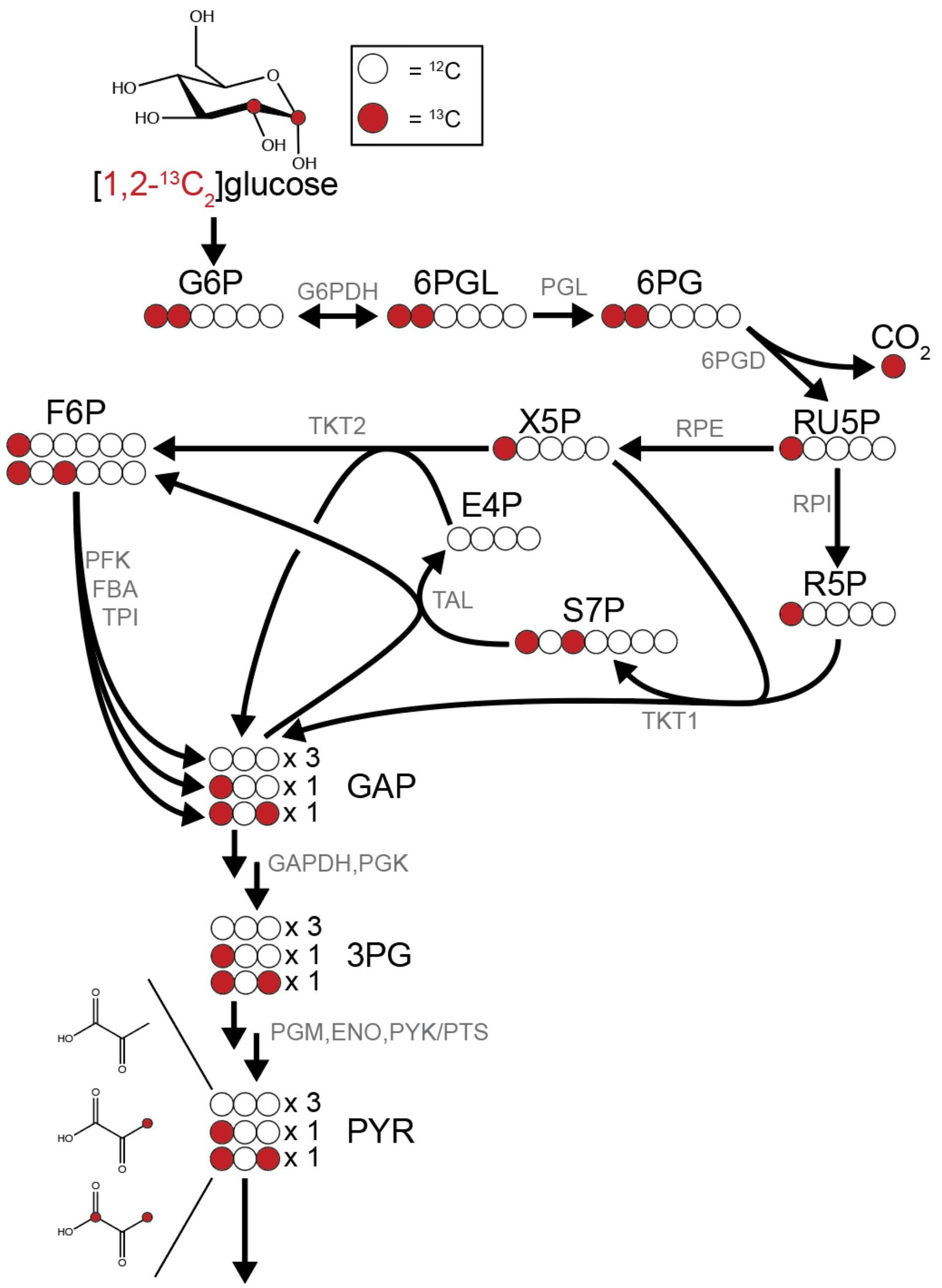


**Supplementary Figure 6. The pentose phosphate pathway generates singly labeled lower glycolytic intermediates.** [1,2-$^{13}C_2$]glucose uniquely generates M+1 labeled triose phosphates through the oxidative pentose phosphate pathway (OxPPP). For every three glucose molecules going through the PPP, three molecules of M+0, one molecule of M+1, and one molecule of M+2 triose phosphate are generated.

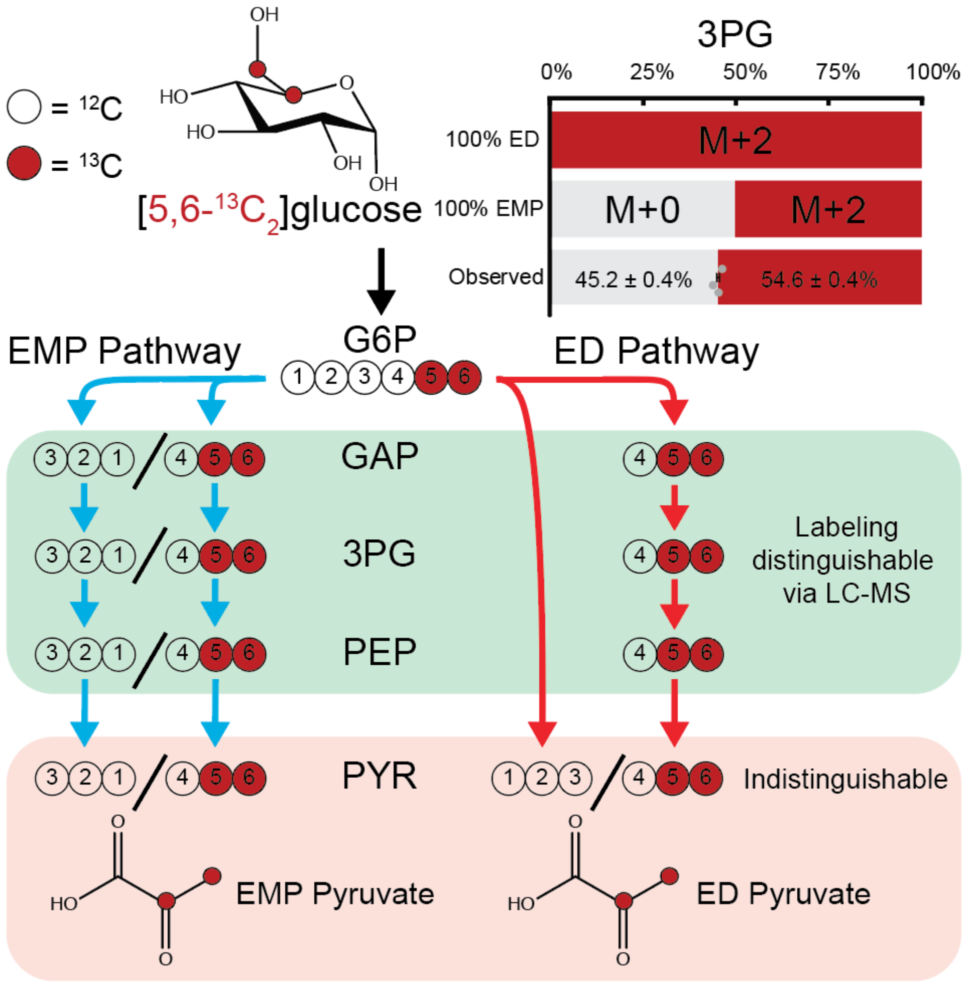


**Supplementary Figure 7. [5,6-$^{13}C_2$]glucose tracing indicates the ED pathway activity.** [5,6-$^{13}C_2$]glucose introduces unlabeled lower glycolytic intermediates only through the EMP pathway. Unlike in the case of [1,2-$^{13}C_2$]glucose, the EMP and the ED pathways produce the same positionally labeled pyruvate. The major fraction of M+2 3PG indicated the ED pathway activity in the nutrient replete condition.

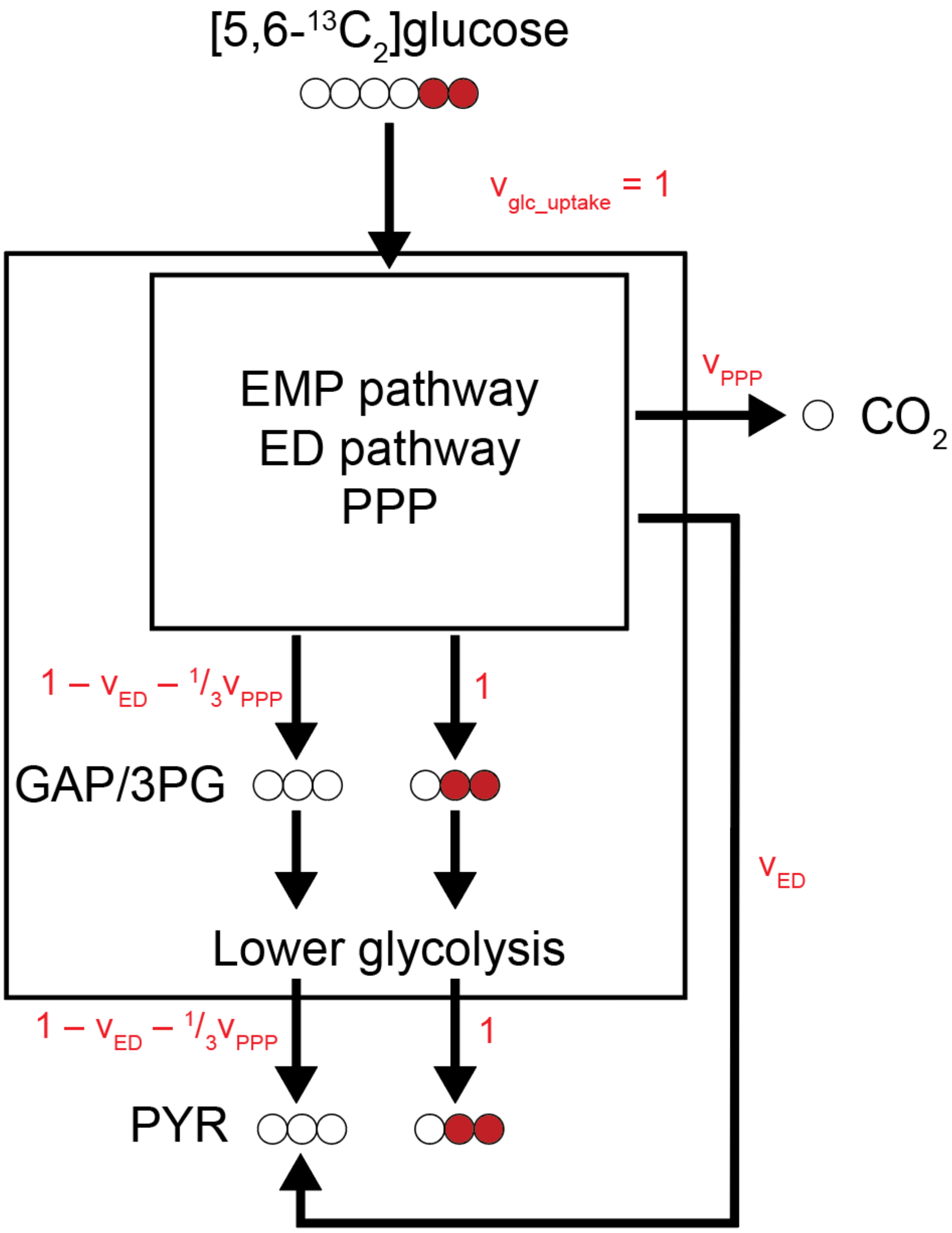


**Supplementary Figure 8. Glycolytic fluxes are solved using 3PG and pyruvate labeling from [5,6-$^{13}C_2$]glucose.** [5,6-$^{13}C_2$]glucose tracing is not convoluted by carbon shuffling in the PPP, so isotopomer distributions of 3PG and pyruvate (PYR) can be used to solve for the ED pathway activity. Mass balance of carbons in the EMP pathway, the ED pathway, and the PPP (the inner control volume) as well as lower glycolysis (the outer control volumes) revealed the relationship between 3PG and PYR labeling and the central carbon metabolism fluxes.

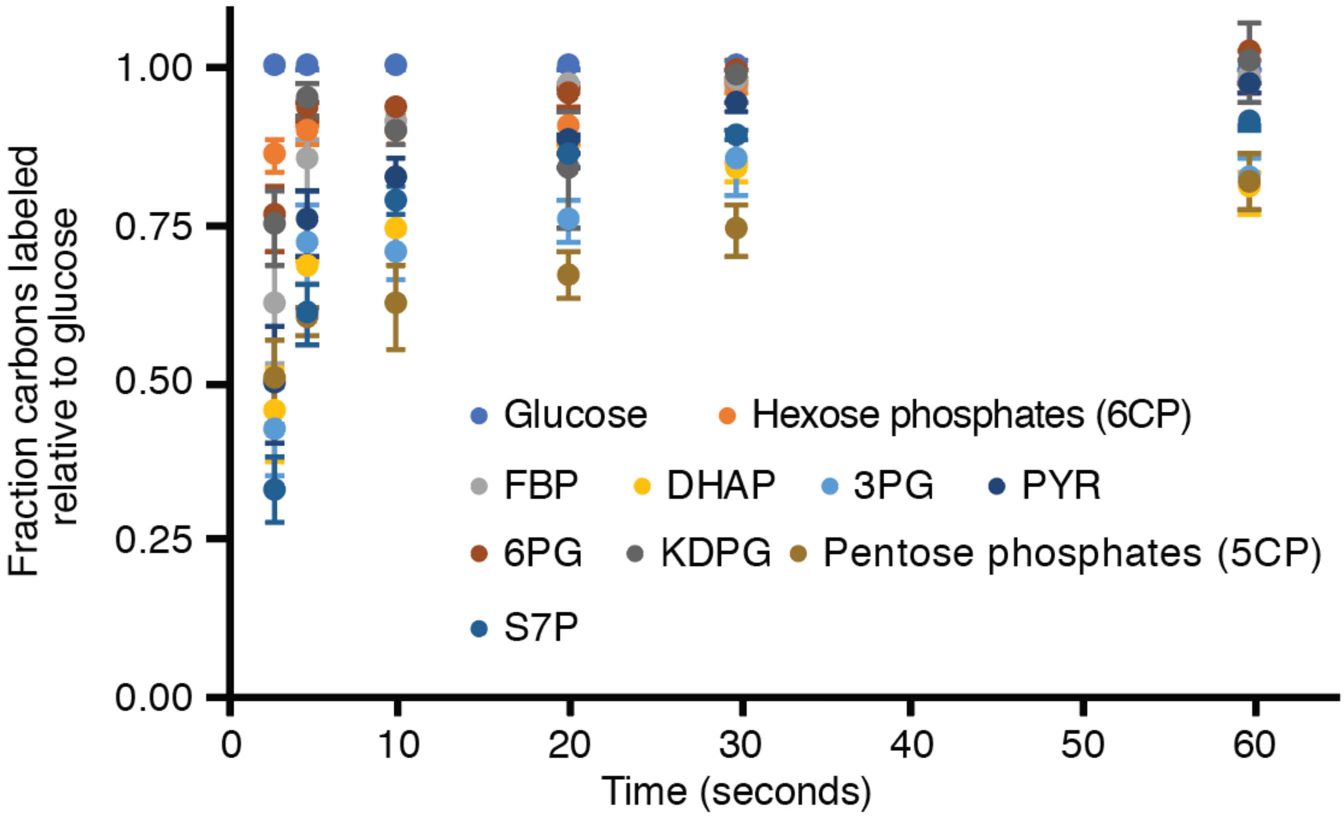


**Supplementary Figure 9. The labeling of glycolytic intermediates quickly reach a pseudo-steady state.** *E. coli* grown on an unlabeled medium were rapidly switched to the same medium containing [U-$^{13}C_6$]glucose. Glycolytic intermediates were substantially labeled within seconds. Error bars represent the s.e.m. (n=3 biological replicates).

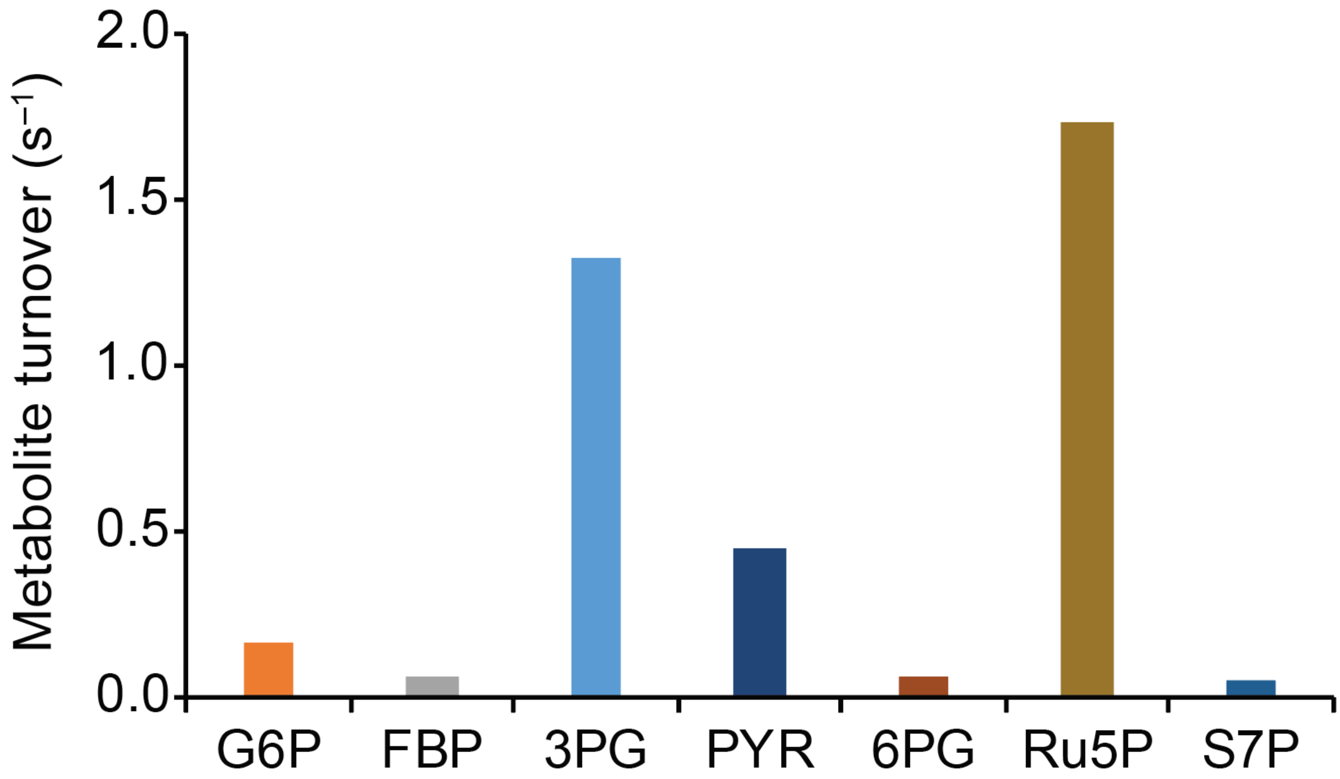


**Supplementary Figure 10. Glycolytic intermediates turn over quickly.** The turnover of individual metabolites was calculated as the flux through that metabolite divided by the metabolite pool size using previously reported fluxes and concentrations[3].

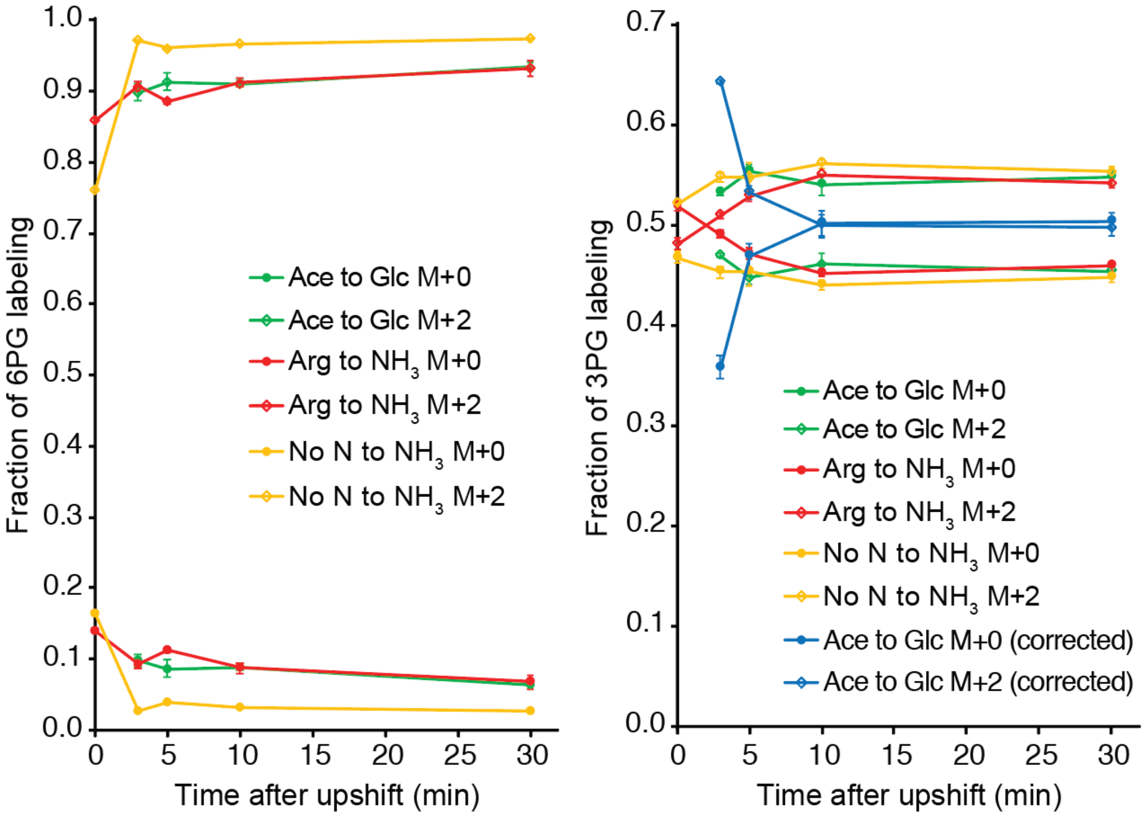


**Supplementary Figure 11. Labeling of 6PG and 3PG in [5,6-$^{13}C_2$]glucose reveals an increasing contribution of the ED pathway to glycolytic flux upon carbon and nitrogen upshift.** Nutrient upshift experiments analogous to those conducted with [1,2-$^{13}C_2$]glucose were replicated using [5,6-$^{13}C_2$]glucose and the resulting labeling of 6PG and 3PG was measured via LC-MS. The labeling of M+0 and M+2 isotopomers revealed both the ED pathway flux and recursive PPP usage (**Supplementary Note 2**). Error bars represent the s.e.m. (n=3 biological replicates).

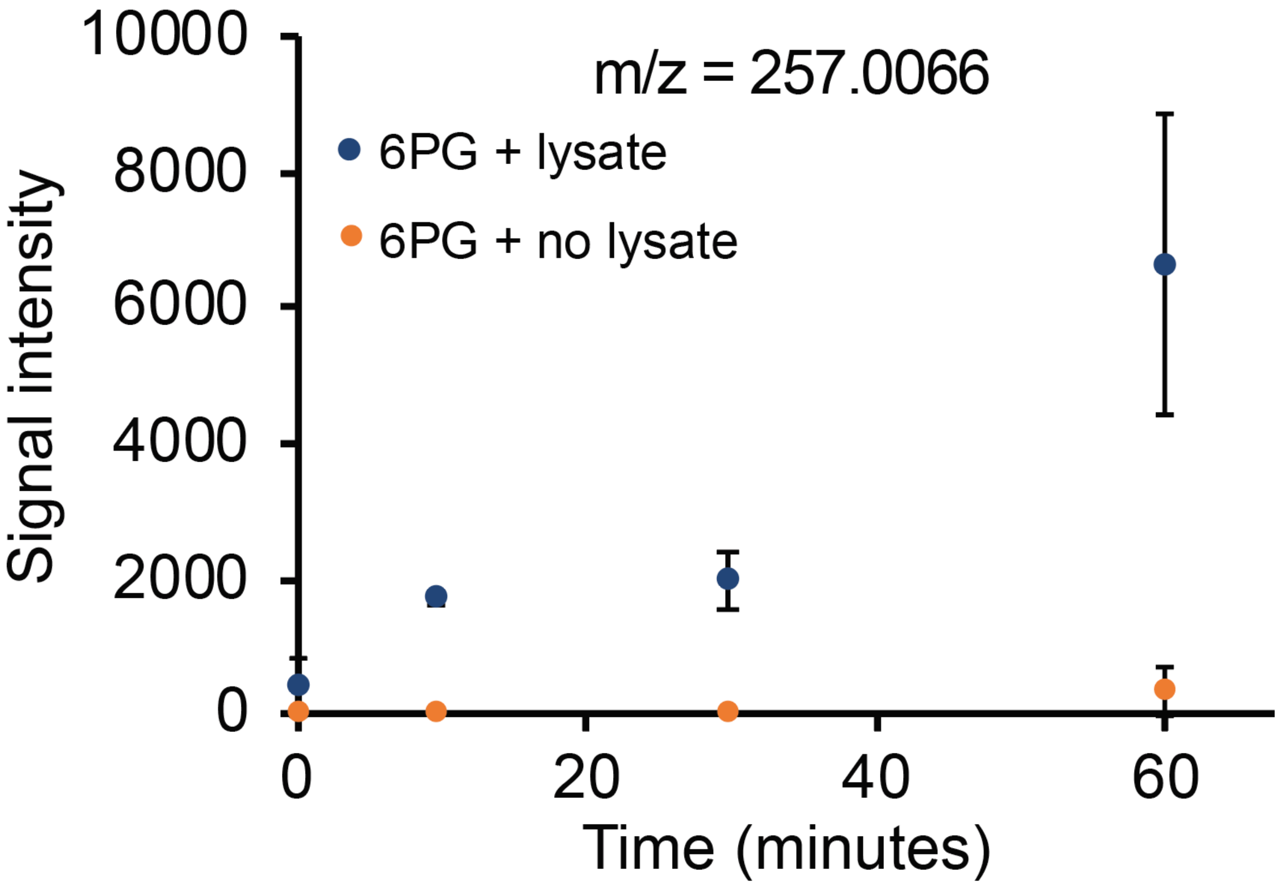


**Supplementary Figure 12. The ED pathway enzymes are present during growth on acetate.** Cell lysates from *E. coli* cultures grown on acetate were collected for an enzyme activity assay and incubated with 6PG, the direct precursor to KDPG (m/z = 257.0066) in the ED pathway. The increase in KDPG over time indicated the presence of the ED pathway enzymes in cells operating on gluconeogenesis. Error bars represent the s.e.m. (n=3 biological replicates).

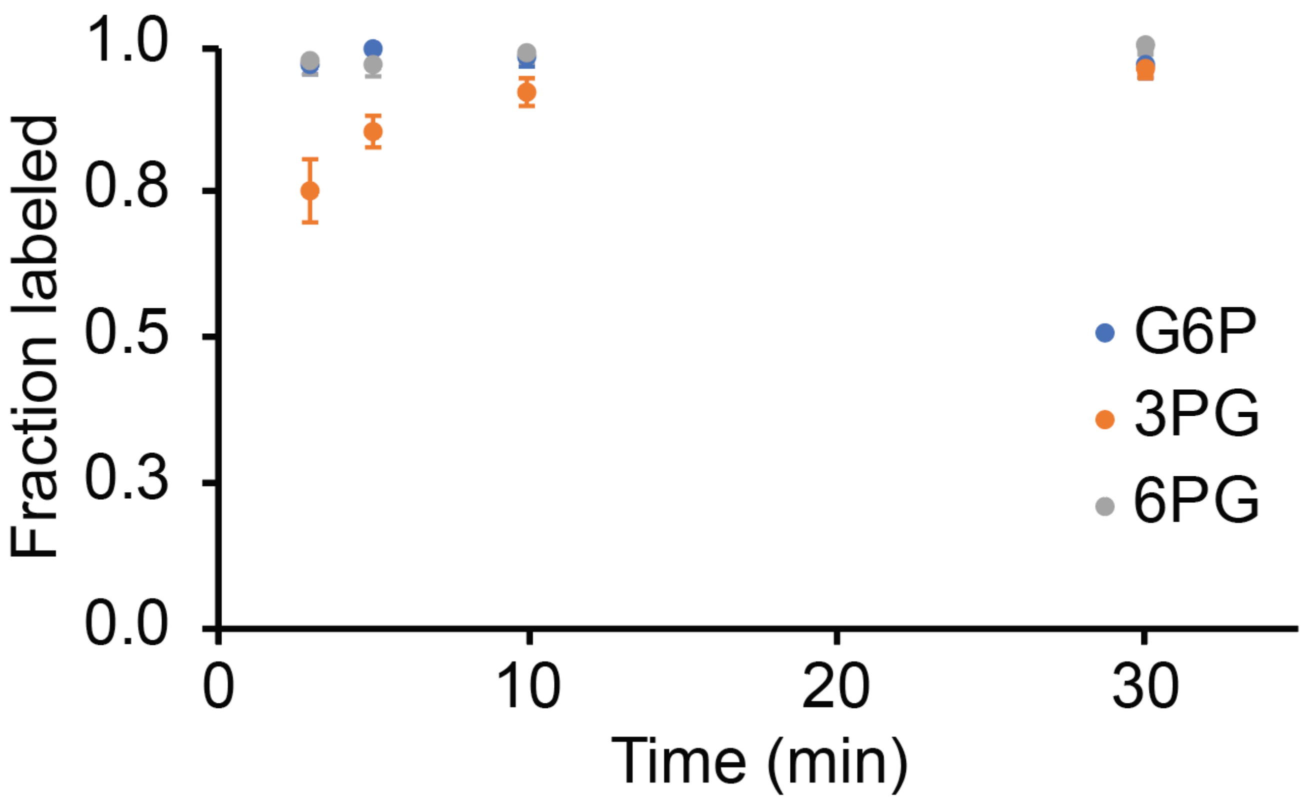


**Supplementary Figure 13. Carbon upshift using [U-$^{13}C_6$]glucose reveals incomplete turnover of lower glycolytic intermediates in early time points.** *E. coli* in the carbon-limited acetate medium underwent carbon upshift by [U-$^{13}C_6$]glucose addition. The labeling of glycolytic intermediates following the upshift informed us of the fractions of 3PG pool coming from the labeled glucose and from the pre-upshift unlabeled acetate. This information was used to accurately compute glycolytic fluxes from [1,2-$^{13}C_2$]glucose tracing. Error bars represent the s.e.m. (n=3 biological replicates).

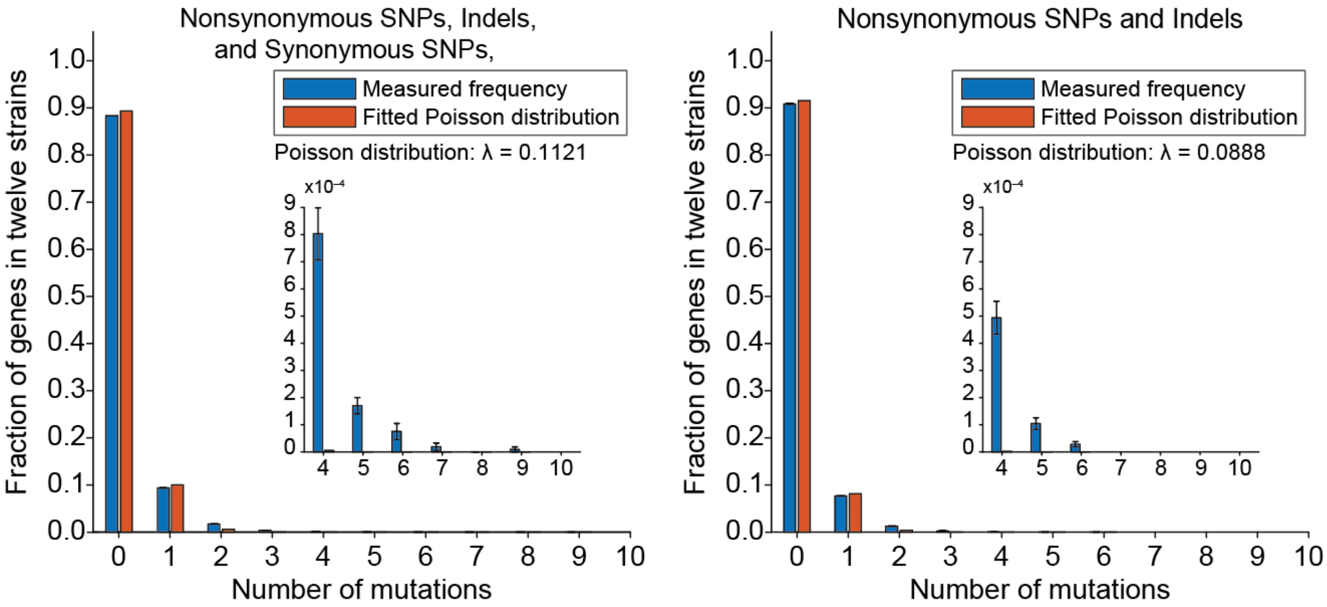


**Supplementary Figure 14. Mutation frequencies in genes do not form a Poisson distribution.** We counted the genes with the same number of mutations, and the normalized counts were fit to a Poisson distribution (see **Methods**). Poor fitting for genes with 2 or more mutations suggested a driving force toward improved fitness.

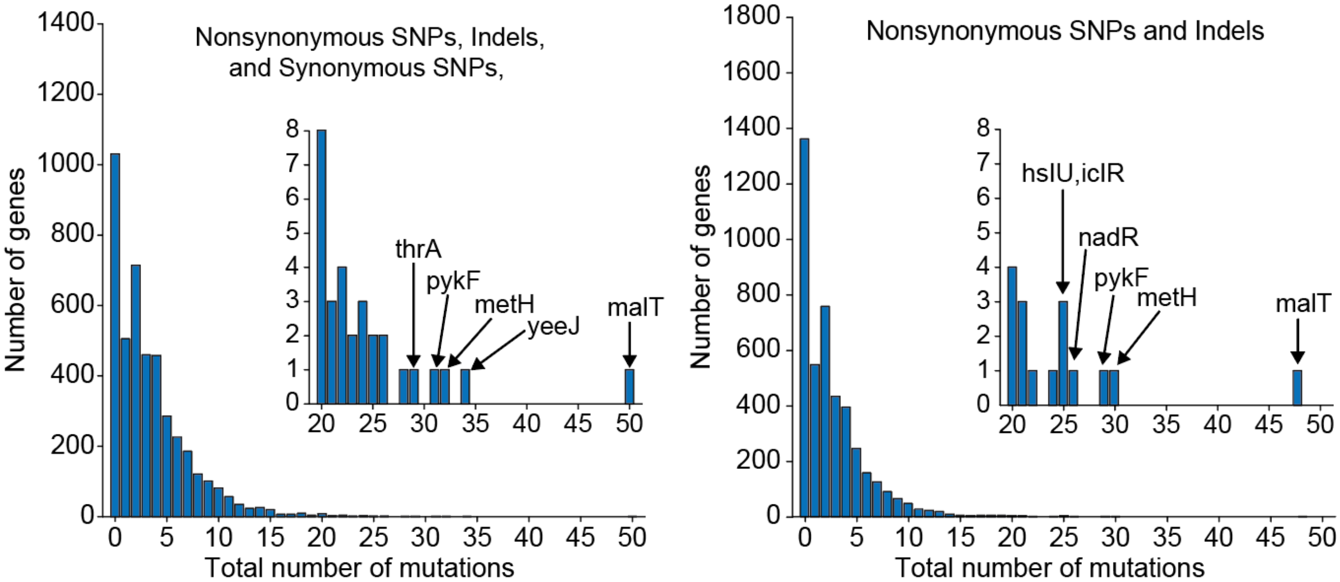


**Supplementary Figure 15. The total number of mutations in individual genes across all 24 clones at the 50,000th generation.** For each gene, we counted the number of mutations in all 24 clones. Few genes were consistently and heavily mutated: malT is a regulator for the maltose regulon that is also involved in λ phage infection; yeeJ is an autotransporter protein; metH is the gene for methionine synthase; pykF is pyruvate kinase, the last step of glycolysis; thrA is a bifunctional aspartokinase and homoserine dehydrogenase; nadR is a regulator for NAD biosynthesis; hsIU is a putative ATP dependent protease; and iclR is a regulator for isocitrate lyase of the glyoxylate cycle.

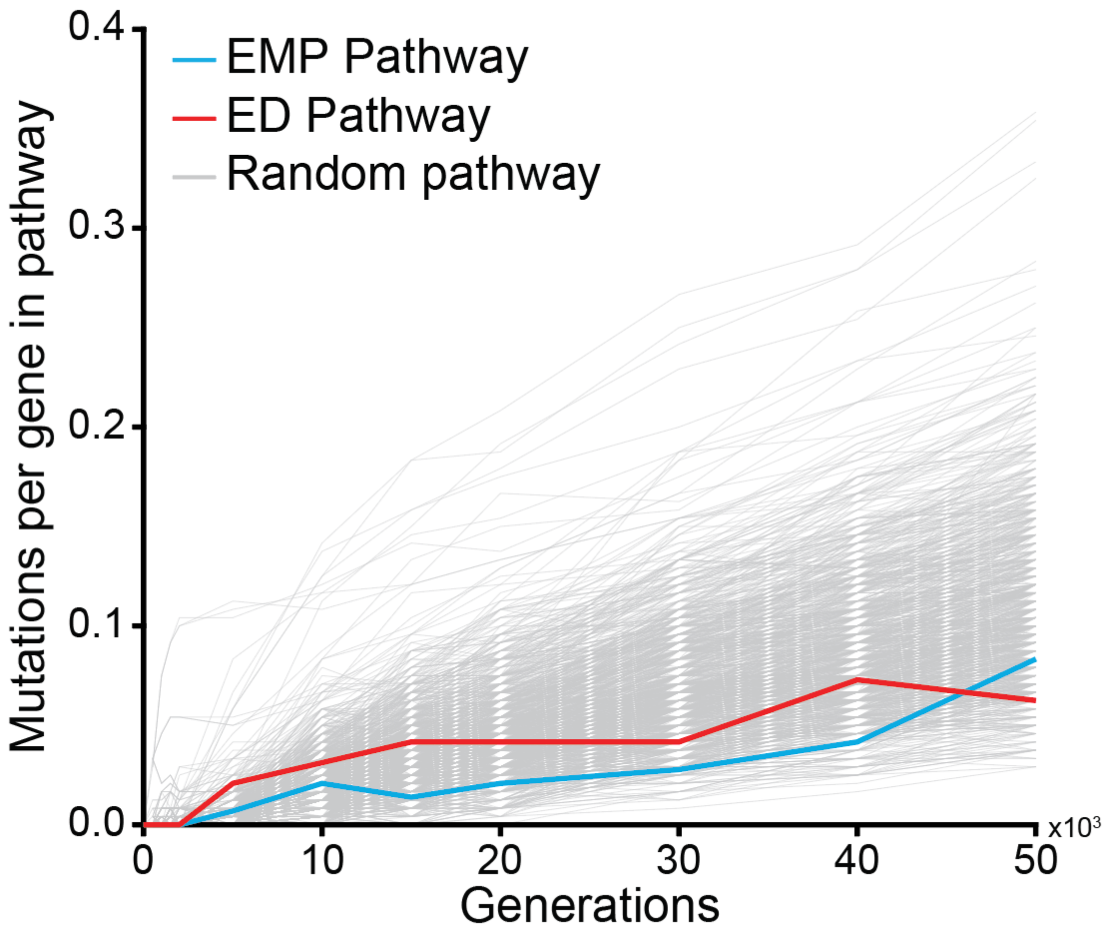


**Supplementary Figure 16. Development of nonsynonymous mutations and indels in randomly sampled gene groups and the parallel glycolytic pathways.** Mutations in randomly generated groups of genes as pseudo-pathways were monitored over 50,000 generations of the long-term evolution experiment. The ED and the EMP pathways initially accumulated more mutations than most other pathways.

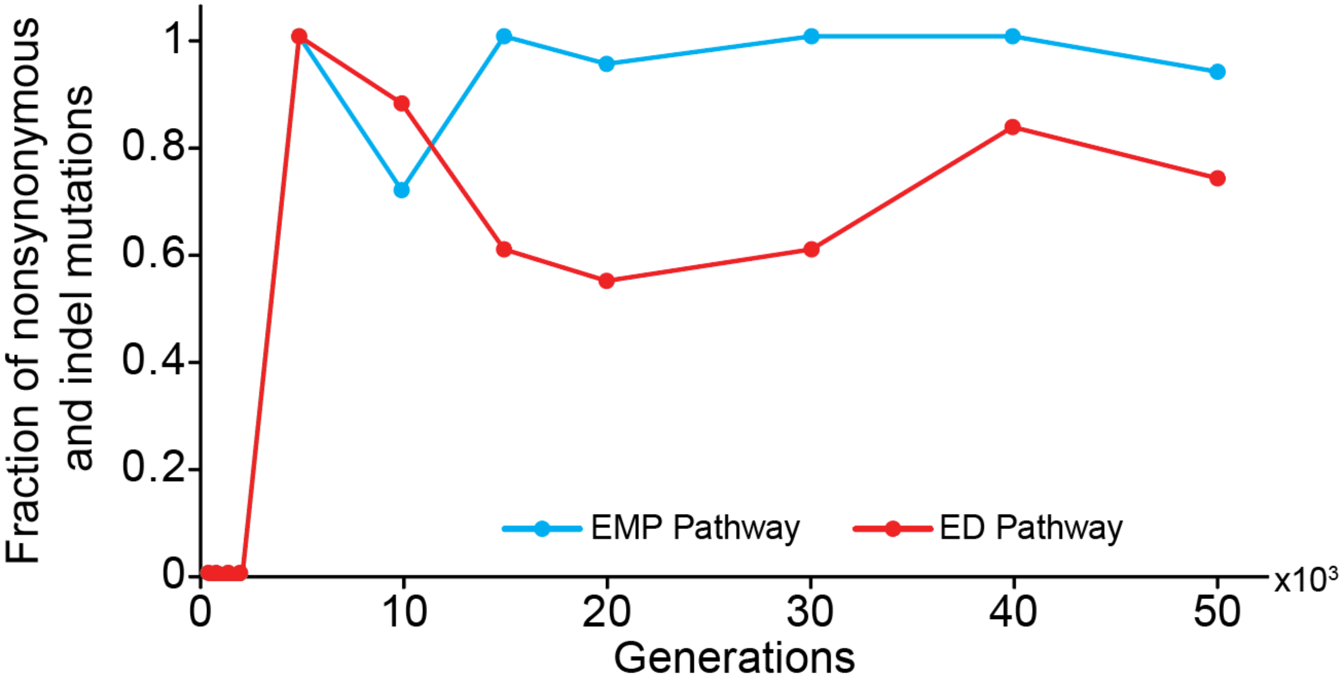


**Supplementary Figure 17. The ED and the EMP pathways have high proportions of nonsynonymous mutations and indels.** A minority of mutations in the two glycolytic pathways were synonymous mutations that do not alter amino acid sequences.

Generation 50,000

| | EMP | | | | | | ED | | | |
|---|---|---|---|---|---|---|---|---|---|---|
| | pgi | pfka | pfkb | fbaa | fbab | tpia | zwf | pgl | edd | eda |
| Ara-1 A | | SNP | | | | | | SNP | | |
| Ara-1 B | | SNP | | | | | | SNP | | |
| Ara-2 A | | | FS | SNP | SNP | | SNP | | SNP | |
| Ara-2 B | | | | | | | | | | |
| Ara-3 A | | | | | 2 SNP | | | | | |
| Ara-3 B | SNP | | | | SNP | | | | | |
| Ara-4 A | | | | | | | | | | |
| Ara-4 B | | | | | | | | | | |
| Ara-5 A | | | | | | | | | | |
| Ara-5 B | | | | | | | | | | |
| Ara-6 A | | SNP | | | | | | | | |
| Ara-6 B | | | | | | | | | | |
| Ara+1 A | | | | | | | | | | |
| Ara+1 B | | | | | | | | | | |
| Ara+2 A | | | | | | | | | | |
| Ara+2 B | | | | | | | | | | |
| Ara+3 A | | | | | SNP | | 2 SNP | | 2 SNP | |
| Ara+3 B | | | | | | | 2 SNP | | 2 SNP | |
| Ara+4 A | | | | | | | | | | |
| Ara+4 B | | | | | | | | | | |
| Ara+5 A | | | | | | | | | | |
| Ara+5 B | | | | | | | | | | |
| Ara+6 A | | | 2 SNP | | | | SNP | | | |
| Ara+6 B | SNP | | | | SNP | | | | | |

**Supplementary Figure 18. List of mutations in the ED and the EMP pathways**. The 24 genome sequences from 12 populations were analyzed at generation 50,000 for mutations in the two glycolytic pathways. SNP denotes a single nucleotide polymorphism and FS denotes a frameshift indel.

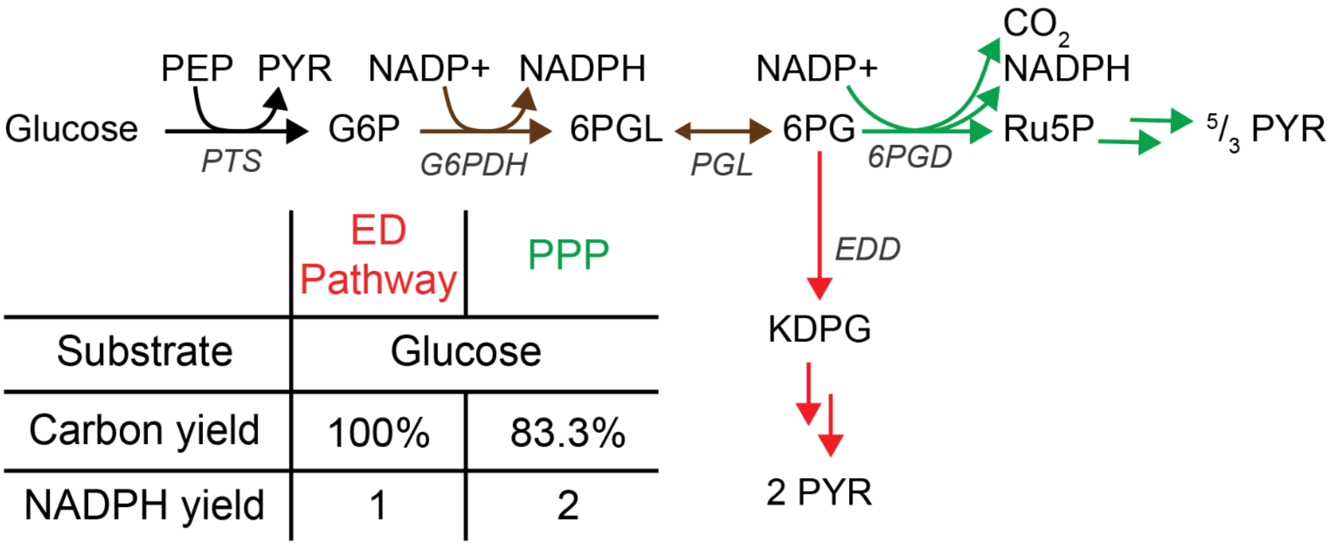


**Supplementary Figure 19. The ED pathway enables carbon-efficient NADPH production**. The OxPPP produces two NADPH, but a carbon is lost as $CO_2$ as 6PG enters the non-oxidative pentose phosphate pathway. On the other hand, the ED pathway produces NADPH without decarboxylation.

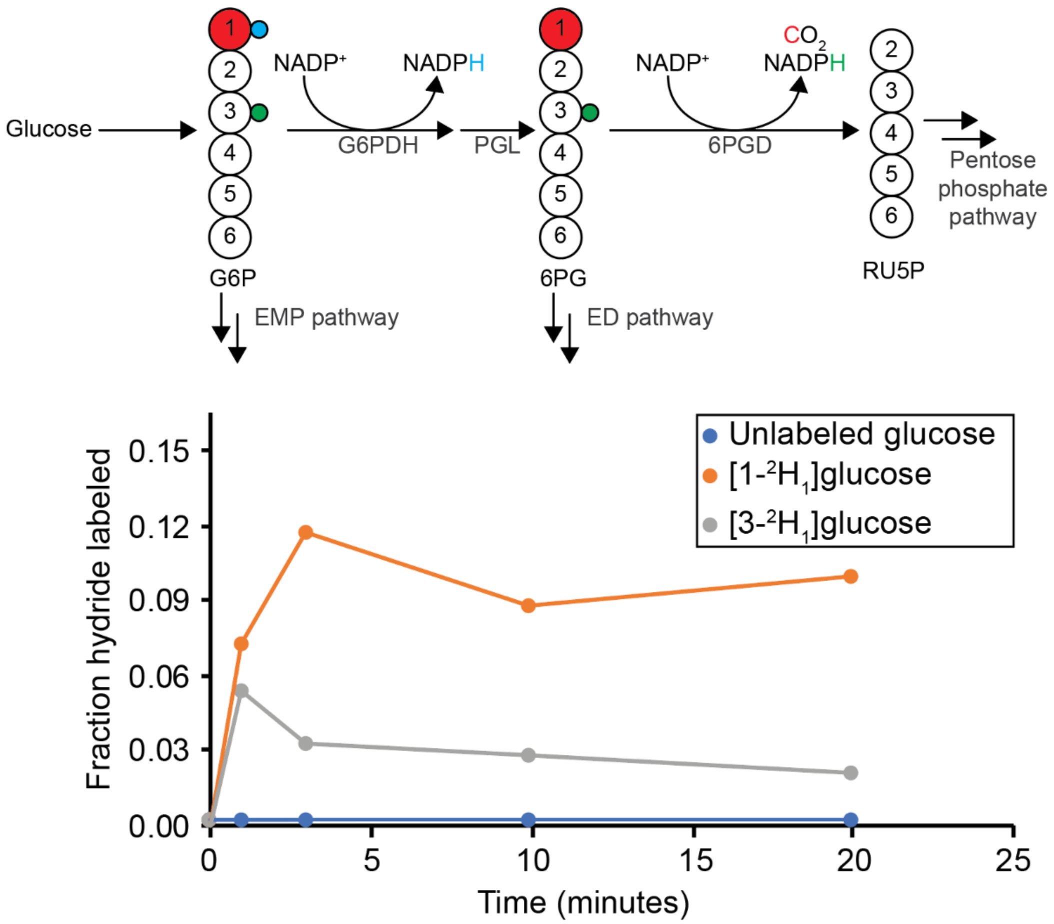


**Supplementary Figure 20. Labeling of NADPH hydride implies substantial contribution of the ED pathway to NADPH production.** Exponentially growing *E. coli* in unlabeled glucose minimal medium were rapidly switched to either [1-$^2H_1$] or [3-$^2H_2$]glucose. Labeling of the hydride of NADPH from [1-$^2H_1$]glucose indicated flux through G6PDH, whereas labeling from [3-$^2H_1$]glucose indicated flux through 6PGD. The ~3-fold labeling of NADPH from G6PDH compared to 6PGD indicated the comparable contributions of the ED pathway and the OxPPP to NADPH generation.

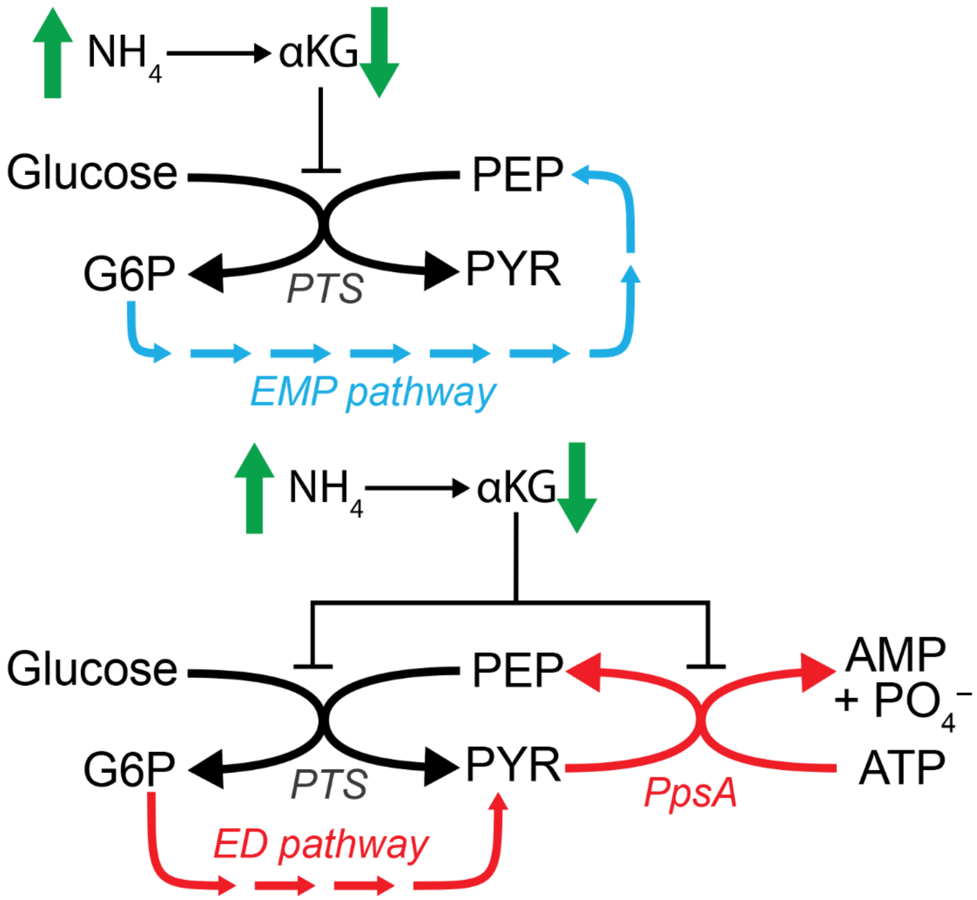


**Supplementary Figure 21. Concerted action of phosphotransferase, phosphoenolpyruvate synthase, and ED pathway streamlines glucose import.** While the EMP pathway takes 8 steps to generate PEP for glucose import via the pts system, the concerted regulation and action of the ED pathway and PpsA can supply PEP in 5 steps.

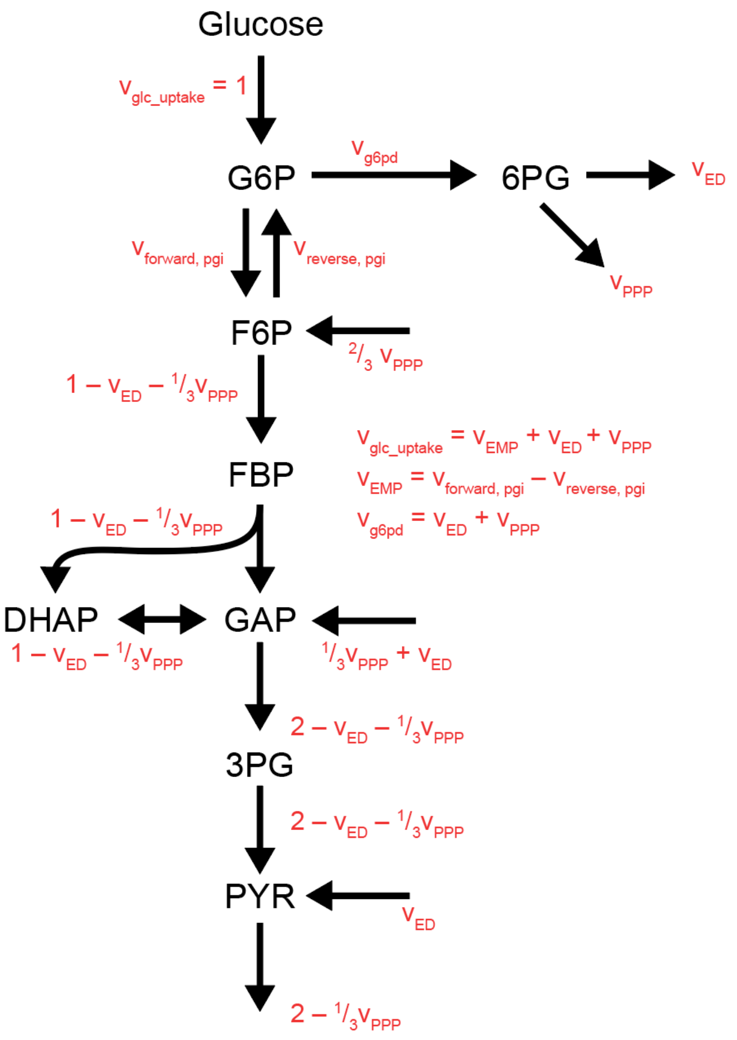


**Supplementary Figure 22. Metabolic model for quantifying fluxes.** Glucose is catabolized through three routes: the EMP pathway, the ED pathway, and the PPP. Fluxes through these pathways are represented by $v_{EMP}$, $v_{ED}$, and $v_{PPP}$. The fluxes were normalized to glucose uptake (i.e., the sum of the three fluxes is 1).

### Supplementary Table 1. Absolute metabolite concentrations of the ED and the EMP glycolytic intermediates under different nutrient conditions

| Metabolite | BiGG ID | KEGG ID | Glucose and NH4 | L.B. | U.B. | Acetate and NH4 | L.B. | U.B. | Glucose and low NH4 | L.B. | U.B. | Glucose and Arg | L.B. | U.B. |
|---|---|---|---|---|---|---|---|---|---|---|---|---|---|---|
| glucose-6-phosphate | g6p | C00092 | 7.88E-03 | 7.59E-03 | 8.17E-03 | 1.06E-03 | 7.92E-04 | 1.42E-03 | 5.18E-03 | 4.60E-03 | 5.84E-03 | 7.87E-03 | 5.15E-03 | 1.20E-02 |
| fructose-6-phosphate | f6p | C00085 | 2.52E-03 | 2.16E-03 | 2.89E-03 | | | | | | | | | |
| fructose-1,6-bisphosphate | fdp | C00354 | 1.52E-02 | 1.40E-02 | 1.64E-02 | 5.38E-04 | 2.89E-04 | 1.00E-03 | 3.82E-03 | 3.64E-03 | 4.01E-03 | 1.13E-02 | 6.80E-03 | 1.88E-02 |
| dihydroxyacetonephosphate | dhap | C00111 | 3.06E-03 | 2.90E-03 | 3.22E-03 | 3.53E-04 | 2.28E-04 | 5.47E-04 | 1.50E-03 | 1.26E-03 | 1.79E-03 | 1.83E-03 | 4.67E-04 | 7.20E-03 |
| 3-phosphoglycerate | 3pg | C00197 | 1.54E-03 | 1.51E-03 | 1.58E-03 | 1.51E-03 | 1.02E-03 | 2.23E-03 | 1.30E-04 | 1.01E-04 | 1.66E-04 | 6.67E-04 | 3.95E-04 | 1.13E-03 |
| phosphoenolpyruvate | pep | C00074 | 1.84E-04 | 1.46E-04 | 2.31E-04 | 8.73E-04 | 5.94E-04 | 1.28E-03 | 6.58E-05 | 6.18E-05 | 7.00E-05 | 6.46E-05 | 5.00E-05 | 8.34E-05 |
| pyruvate | pyr | C00022 | 3.66E-03 | 3.13E-03 | 4.20E-03 | 1.93E-03 | 1.28E-03 | 2.92E-03 | 3.52E-04 | 2.31E-04 | 5.36E-04 | 2.30E-03 | 1.83E-03 | 2.91E-03 |
| 6-phospho-D-gluconate | 6pgc | C00345 | 3.77E-03 | 3.69E-03 | 3.85E-03 | 2.09E-04 | 1.63E-04 | 2.68E-04 | 2.75E-04 | 2.07E-04 | 3.65E-04 | 3.19E-04 | 2.64E-04 | 3.86E-04 |
| 2-keto-3-deoxy-6-phosphogluconate | 2ddg6p | C04442 | 2.18E-04 | 1.89E-04 | 2.52E-04 | 1.28E-06 | 3.84E-07 | 4.27E-06 | 1.83E-03 | 2.24E-03 | 1.69E-03 | 6.59E-05 | 3.46E-05 | 1.25E-04 |
| ATP | atp | C00002 | 9.63E-03 | 8.13E-03 | 1.14E-02 | 6.05E-03 | 3.80E-03 | 9.62E-03 | 4.25E-03 | 3.11E-03 | 5.82E-03 | 4.28E-03 | 3.84E-03 | 4.77E-03 |
| ADP | adp | C00008 | 5.55E-04 | 4.37E-04 | 7.04E-04 | 2.35E-04 | 9.10E-05 | 6.07E-04 | 7.17E-05 | 5.37E-05 | 9.57E-05 | 3.17E-04 | 2.34E-04 | 4.30E-04 |
| AMP | amp | C00020 | 2.81E-04 | 2.32E-04 | 3.41E-04 | 1.01E-03 | 3.87E-04 | 2.62E-03 | 3.63E-05 | 2.01E-05 | 6.57E-05 | 4.43E-04 | 2.86E-04 | 6.86E-04 |
| NAD+ | nad | C00003 | 2.55E-03 | 2.32E-03 | 2.80E-03 | 2.86E-03 | 2.51E-03 | 3.26E-03 | 9.13E-04 | 7.21E-04 | 1.16E-03 | 1.76E-03 | 9.65E-04 | 3.22E-03 |
| NADH | nadh | C00004 | 8.36E-05 | 5.45E-05 | 1.27E-04 | 7.27E-06 | 5.87E-06 | 9.01E-06 | 3.65E-05 | 3.35E-05 | 3.98E-05 | 5.67E-05 | 3.15E-05 | 1.02E-04 |
| NADP+ | nadp | C00006 | 2.08E-06 | 1.40E-07 | 3.11E-05 | 1.62E-06 | 1.31E-06 | 2.00E-06 | 3.44E-07 | 2.83E-07 | 4.18E-07 | 1.06E-06 | 8.19E-07 | 1.38E-06 |
| NADPH | nadph | C00005 | 1.21E-04 | 1.10E-04 | 1.34E-04 | 2.98E-04 | 5.22E-05 | 1.70E-03 | 2.20E-05 | 1.72E-05 | 2.82E-05 | 5.63E-05 | 3.96E-05 | 8.00E-05 |
| phosphate (orthophosphate) | pi | C00009 | 2.39E-02 | 1.60E-02 | 2.40E-02 | 2.39E-02 | 1.60E-02 | 2.40E-02 | 2.39E-02 | 1.60E-02 | 2.40E-02 | 2.39E-02 | 1.60E-02 | 2.40E-02 |

L.B. and U.B. represent the lower and upper bounds of the 95% confidence interval.

### Supplementary Table 2. Isotopic labeling patterns of glycolytic intermediates in WT *E. coli* in nutrient replete medium

| Metabolite | Labeling | [1,2-$^{13}C_2$]glucose | s.e.m. | [5,6-13C2]glucose | s.e.m. |
|---|---|---|---|---|---|
| G6P/F6P | M+0 | 0.1% | 0.0% | 7.5% | 0.3% |
| | M+1 | 0.6% | 0.0% | 0.0% | 0.0% |
| | M+2 | 95.2% | 0.1% | 92.5% | 0.3% |
| | M+3 | 0.6% | 0.0% | 0.0% | 0.0% |
| | M+4 | 3.4% | 0.1% | 0.0% | 0.0% |
| | M+5 | 0.1% | 0.1% | 0.0% | 0.0% |
| | M+6 | 0.0% | 0.0% | 0.0% | 0.0% |
| 6PG | M+0 | 0.0% | 0.0% | 2.4% | 0.2% |
| | M+1 | 0.0% | 0.0% | 0.0% | 0.0% |
| | M+2 | 96.9% | 0.4% | 97.6% | 0.2% |
| | M+3 | 0.5% | 0.1% | 0.0% | 0.0% |
| | M+4 | 1.9% | 0.1% | 0.0% | 0.0% |
| | M+5 | 0.1% | 0.0% | 0.0% | 0.0% |
| | M+6 | 0.5% | 0.3% | 0.0% | 0.0% |
| 3PG | M+0 | 52.7% | 0.2% | 45.2% | 0.4% |
| | M+1 | 1.2% | 0.1% | 0.0% | 0.0% |
| | M+2 | 45.6% | 0.1% | 54.8% | 0.4% |
| | M+3 | 0.5% | 0.3% | 0.0% | 0.0% |
| PYR | M+0 | 51.7% | 0.8% | 45.6% | 0.4% |
| | M+1 | 4.4% | 0.1% | 2.4% | 0.2% |
| | M+2 | 42.8% | 0.6% | 50.9% | 0.3% |
| | M+3 | 1.0% | 0.2% | 1.0% | 0.0% |

G6P denotes glucose-6-phosphate; F6P, fructose-6-phosphate; 3PG, 3-phosphoglycerate; 6PG, 6-phosphogluconate; PYR, pyruvate. Values represent the mean and standard error (n=3 biological replicates).

**Supplementary Table 3. Isotopic labeling patterns of valine its $MS^2$ fragments in WT *E. coli* using [1,2-$^{13}C_2$]glucose in nutrient replete medium**

| Parent valine | mean | s.e.m. | $MS^2$ fragment | mean | s.e.m. |
|---|---|---|---|---|---|
| M+0 | 28.5% | 0.6% | M+0 | 100.0% | 0.0% |
| | | | M+1 | 0.0% | 0.0% |
| | | | M+2 | 0.0% | 0.0% |
| | | | M+3 | 0.0% | 0.0% |
| | | | M+4 | 0.0% | 0.0% |
| M+1 | 5.3% | 0.1% | M+0 | 9.0% | 3.6% |
| | | | M+1 | 91.0% | 3.6% |
| | | | M+2 | 0.0% | 0.0% |
| | | | M+3 | 0.0% | 0.0% |
| | | | M+4 | 0.0% | 0.0% |
| M+2 | 41.3% | 0.3% | M+0 | 0.0% | 0.0% |
| | | | M+1 | 3.7% | 0.2% |
| | | | M+2 | 96.3% | 0.2% |
| | | | M+3 | 0.0% | 0.0% |
| | | | M+4 | 0.0% | 0.0% |
| M+3 | 5.1% | 0.1% | M+0 | 0.0% | 0.0% |
| | | | M+1 | 0.0% | 0.0% |
| | | | M+2 | 23.5% | 1.2% |
| | | | M+3 | 76.5% | 1.2% |
| | | | M+4 | 0.0% | 0.0% |
| M+4 | 19.7% | 0.4% | M+0 | 0.0% | 0.0% |
| | | | M+1 | 0.0% | 0.0% |
| | | | M+2 | 0.0% | 0.0% |
| | | | M+3 | 4.9% | 0.2% |
| | | | M+4 | 95.1% | 0.2% |
| M+5 | 0.0% | 0.0% | M+0 | 0.0% | 0.0% |
| | | | M+1 | 0.0% | 0.0% |
| | | | M+2 | 0.0% | 0.0% |
| | | | M+3 | 2.0% | 0.0% |
| | | | M+4 | 98.0% | 0.0% |

Values represent the mean and standard error (n=3 biological replicates).

**Supplementary Table 4. Isotopic labeling patterns of glycolytic intermediates in WT *E. coli* using [U-$^{13}C_6$]glucose in carbon upshift**

| Metabolite | Labeling | 3 minutes post-upshift | s.e.m. | 5 minutes post-upshift | s.e.m. | 10 minutes post-upshift | s.e.m | 30 minutes post-upshift | s.e.m. |
|---|---|---|---|---|---|---|---|---|---|
| G6P/F6P | M+0 | 1.8% | 0.4% | 0.8% | 0.4% | 1.3% | 0.1% | 2.8% | 1.4% |
| | M+1 | 0.0% | 0.0% | 0.0% | 0.0% | 0.0% | 0.0% | 0.4% | 0.1% |
| | M+2 | 0.0% | 0.0% | 0.0% | 0.0% | 0.1% | 0.0% | 0.6% | 0.1% |
| | M+3 | 1.4% | 0.3% | 0.5% | 0.3% | 0.8% | 0.3% | 0.2% | 0.1% |
| | M+4 | 0.2% | 0.0% | 0.1% | 0.0% | 0.1% | 0.1% | 0.0% | 0.0% |
| | M+5 | 0.5% | 0.2% | 0.2% | 0.1% | 0.3% | 0.2% | 0.1% | 0.1% |
| | M+6 | 96.1% | 0.9% | 98.4% | 0.8% | 97.3% | 0.4% | 95.9% | 1.5% |
| 3PG | M+0 | 27.0% | 1.1% | 16.1% | 1.2% | 7.8% | 0.9% | 4.4% | 0.2% |
| | M+1 | 0.0% | 0.0% | 0.0% | 0.0% | 0.0% | 0.0% | 0.0% | 0.0% |
| | M+2 | 0.0% | 0.0% | 0.0% | 0.0% | 0.0% | 0.0% | 0.5% | 0.3% |
| | M+3 | 73.0% | 1.1% | 83.9% | 1.2% | 92.2% | 0.9% | 95.0% | 0.4% |
| 6PG | M+0 | 0.0% | 0.0% | 0.0% | 0.0% | 0.0% | 0.0% | 33.3% | 0.0% |
| | M+1 | 0.0% | 0.0% | 0.0% | 0.0% | 0.0% | 0.0% | 0.0% | 0.0% |
| | M+2 | 0.0% | 0.0% | 1.2% | 0.7% | 0.0% | 0.0% | 0.0% | 0.0% |
| | M+3 | 2.1% | 0.7% | 1.4% | 0.5% | 0.9% | 0.4% | 0.0% | 0.0% |
| | M+4 | 0.0% | 0.0% | 0.2% | 0.2% | 0.0% | 0.0% | 0.0% | 0.0% |
| | M+5 | 1.4% | 0.6% | 1.2% | 0.6% | 1.2% | 0.4% | 0.7% | 0.3% |
| | M+6 | 96.4% | 1.1% | 96.0% | 1.9% | 97.8% | 0.7% | 99.2% | 0.3% |

G6P denotes glucose-6-phosphate; F6P, fructose-6-phosphate; 3PG, 3-phosphoglycerate; 6PG, 6-phosphogluconate. Values represent the mean and standard error (n=3 biological replicates).

**Supplementary Table 5. Isotopic labeling patterns of glycolytic intermediates in WT *E. coli* using [1,2-$^{13}C_2$]glucose in carbon upshift**

| Metabolite | Labeling | 3 minutes post-upshift | s.e.m. | 5 minutes post-upshift | s.e.m. | 10 minutes post-upshift | s.e.m | 30 minutes post-upshift | s.e.m. |
|---|---|---|---|---|---|---|---|---|---|
| G6P/F6P | M+0 | 4.8% | 0.5% | 5.8% | 0.3% | 3.9% | 0.2% | 2.4% | 0.3% |
| | M+1 | 5.2% | 0.8% | 5.0% | 0.0% | 3.7% | 0.2% | 1.6% | 0.2% |
| | M+2 | 81.0% | 1.2% | 79.1% | 0.4% | 83.0% | 0.5% | 88.4% | 0.7% |
| | M+3 | 4.0% | 0.4% | 4.1% | 0.3% | 3.2% | 0.2% | 2.3% | 0.2% |
| | M+4 | 4.2% | 0.3% | 4.9% | 0.0% | 5.3% | 0.1% | 4.5% | 0.4% |
| | M+5 | 0.7% | 0.1% | 0.9% | 0.1% | 0.9% | 0.0% | 0.8% | 0.0% |
| | M+6 | 0.1% | 0.0% | 0.1% | 0.0% | 0.0% | 0.0% | 0.1% | 0.0% |
| FBP | M+0 | 26.5% | 0.2% | 25.4% | 0.6% | 21.3% | 0.4% | 13.6% | 0.1% |
| | M+1 | 7.7% | 0.3% | 7.3% | 0.2% | 6.3% | 0.1% | 3.8% | 0.1% |
| | M+2 | 50.2% | 0.3% | 51.2% | 0.7% | 53.7% | 0.6% | 60.6% | 0.5% |
| | M+3 | 5.8% | 0.3% | 5.9% | 0.4% | 5.7% | 0.3% | 4.7% | 0.1% |
| | M+4 | 8.2% | 0.4% | 8.8% | 0.3% | 11.2% | 0.6% | 15.5% | 0.3% |
| | M+5 | 0.7% | 0.2% | 1.2% | 0.1% | 1.4% | 0.1% | 1.6% | 0.0% |
| | M+6 | 0.8% | 0.1% | 0.2% | 0.1% | 0.3% | 0.1% | 0.3% | 0.0% |
| 3PG | M+0 | 77.9% | 0.4% | 71.3% | 0.5% | 62.6% | 0.4% | 59.4% | 0.5% |
| | M+1 | 3.7% | 0.5% | 4.1% | 0.4% | 4.9% | 0.5% | 5.2% | 0.2% |
| | M+2 | 18.4% | 0.3% | 24.6% | 0.3% | 32.5% | 0.8% | 35.4% | 0.2% |
| | M+3 | 0.0% | 0.0% | 0.0% | 0.0% | 0.0% | 0.0% | 0.0% | 0.0% |
| 6PG | M+0 | 1.8% | 0.1% | 2.0% | 0.2% | 1.3% | 0.1% | 0.5% | 0.1% |
| | M+1 | 4.9% | 0.2% | 4.7% | 0.2% | 3.8% | 0.3% | 2.1% | 0.1% |
| | M+2 | 86.0% | 0.1% | 84.9% | 0.3% | 87.0% | 0.5% | 91.5% | 0.2% |
| | M+3 | 2.9% | 0.2% | 3.2% | 0.2% | 2.2% | 0.1% | 1.9% | 0.1% |
| | M+4 | 3.7% | 0.2% | 4.3% | 0.2% | 4.7% | 0.3% | 3.5% | 0.0% |
| | M+5 | 0.5% | 0.1% | 0.8% | 0.1% | 0.9% | 0.1% | 0.6% | 0.2% |
| | M+6 | 0.1% | 0.0% | 0.1% | 0.0% | 0.0% | 0.0% | 0.1% | 0.0% |

G6P denotes glucose-6-phosphate; F6P, fructose-6-phosphate; FBP, fructose-1,6-bisphosphate; 3PG, 3-phosphoglycerate; 6PG, 6-phosphogluconate. Values represent the mean and standard error (n=3 biological replicates).

**Supplementary Table 6. Isotopic labeling patterns of glycolytic intermediates in WT *E. coli* using $[5,6\text{-}^{13}C_2]$glucose in carbon upshift**

| Metabolite | Labeling | 3 minutes post-upshift | s.e.m. | 5 minutes post-upshift | s.e.m. | 10 minutes post-upshift | s.e.m | 30 minutes post-upshift | s.e.m. |
|---|---|---|---|---|---|---|---|---|---|
| G6P/F6P | M+0 | 18.4% | 0.9% | 17.8% | 0.5% | 16.7% | 1.9% | 14.2% | 1.1% |
| | M+1 | 0.0% | 0.0% | 0.0% | 0.0% | 0.0% | 0.0% | 0.0% | 0.0% |
| | M+2 | 81.1% | 0.9% | 81.9% | 0.4% | 82.8% | 1.9% | 85.3% | 1.1% |
| | M+3 | 0.0% | 0.0% | 0.0% | 0.0% | 0.0% | 0.0% | 0.0% | 0.0% |
| | M+4 | 0.4% | 0.0% | 0.3% | 0.1% | 0.5% | 0.1% | 0.5% | 0.1% |
| | M+5 | 0.0% | 0.0% | 0.0% | 0.0% | 0.0% | 0.0% | 0.0% | 0.0% |
| | M+6 | 0.0% | 0.0% | 0.0% | 0.0% | 0.0% | 0.0% | 0.0% | 0.0% |
| FBP | M+0 | 23.5% | 1.6% | 23.0% | 1.1% | 21.7% | 0.6% | 17.8% | 0.8% |
| | M+1 | 0.0% | 0.0% | 0.0% | 0.0% | 0.0% | 0.0% | 0.0% | 0.0% |
| | M+2 | 69.3% | 0.4% | 72.0% | 1.8% | 73.4% | 0.9% | 80.5% | 1.0% |
| | M+3 | 0.3% | 0.3% | 0.1% | 0.1% | 0.0% | 0.0% | 0.0% | 0.0% |
| | M+4 | 5.7% | 0.9% | 4.9% | 0.6% | 4.3% | 0.2% | 1.6% | 0.3% |
| | M+5 | 0.0% | 0.0% | 0.0% | 0.0% | 0.0% | 0.0% | 0.0% | 0.0% |
| | M+6 | 1.3% | 0.6% | 0.0% | 0.0% | 0.6% | 0.5% | 0.1% | 0.1% |
| 3PG | M+0 | 53.1% | 0.2% | 55.4% | 0.7% | 54.0% | 1.0% | 52.7% | 0.7% |
| | M+1 | 0.0% | 0.0% | 0.0% | 0.0% | 0.0% | 0.0% | 0.0% | 0.0% |
| | M+2 | 46.9% | 0.2% | 44.6% | 0.7% | 46.0% | 1.0% | 47.3% | 0.7% |
| | M+3 | 0.0% | 0.0% | 0.0% | 0.0% | 0.0% | 0.0% | 0.0% | 0.0% |
| 6PG | M+0 | 9.8% | 1.0% | 8.7% | 1.2% | 9.0% | 0.5% | 6.4% | 0.7% |
| | M+1 | 0.0% | 0.0% | 0.0% | 0.0% | 0.0% | 0.0% | 0.0% | 0.0% |
| | M+2 | 89.8% | 1.2% | 91.3% | 1.2% | 91.0% | 0.5% | 93.4% | 0.7% |
| | M+3 | 0.3% | 0.3% | 0.0% | 0.0% | 0.0% | 0.0% | 0.0% | 0.0% |
| | M+4 | 0.0% | 0.0% | 0.0% | 0.0% | 0.0% | 0.0% | 0.1% | 0.1% |
| | M+5 | 0.0% | 0.0% | 0.0% | 0.0% | 0.0% | 0.0% | 0.0% | 0.0% |
| | M+6 | 0.0% | 0.0% | 0.0% | 0.0% | 0.0% | 0.0% | 0.1% | 0.1% |

G6P denotes glucose-6-phosphate; F6P, fructose-6-phosphate; FBP, fructose-1,6-bisphosphate; 3PG, 3-phosphoglycerate; 6PG, 6-phosphogluconate. Values represent the mean and standard error (n=3 biological replicates).

**Supplementary Table 7. Isotopic labeling patterns of glycolytic intermediates in WT *E. coli* using [1,2-$^{13}C_2$]glucose in arginine to ammonia upshift**

| Metabolite | Labeling | 0 minutes post-upshift | s.e.m. | 3 minutes post-upshift | s.e.m. | 5 minutes post-upshift | s.e.m | 10 minutes post-upshift | s.e.m. | 30 minutes post-upshift | s.e.m |
|---|---|---|---|---|---|---|---|---|---|---|---|
| G6P/F6P | M+0 | 2.4% | 0.1% | 0.9% | 0.0% | 0.7% | 0.0% | 0.6% | 0.1% | 0.7% | 0.1% |
| | M+1 | 0.6% | 0.0% | 0.9% | 0.2% | 0.3% | 0.1% | 0.9% | 0.1% | 1.1% | 0.3% |
| | M+2 | 81.9% | 0.2% | 91.4% | 0.6% | 92.7% | 0.2% | 90.6% | 0.5% | 91.9% | 0.2% |
| | M+3 | 2.9% | 0.1% | 1.3% | 0.2% | 1.2% | 0.4% | 1.7% | 0.4% | 1.3% | 0.1% |
| | M+4 | 10.2% | 0.1% | 4.6% | 0.2% | 4.5% | 0.1% | 5.5% | 0.1% | 4.4% | 0.1% |
| | M+5 | 1.9% | 0.2% | 0.9% | 0.1% | 0.6% | 0.1% | 0.6% | 0.0% | 0.6% | 0.1% |
| | M+6 | 0.1% | 0.0% | 0.1% | 0.0% | 0.0% | 0.0% | 0.1% | 0.0% | 0.1% | 0.0% |
| FBP | M+0 | 11.7% | 0.3% | 7.4% | 0.4% | 8.0% | 0.5% | 8.6% | 0.2% | 6.9% | 0.5% |
| | M+1 | 1.2% | 0.2% | 1.9% | 0.1% | 1.9% | 0.3% | 1.7% | 0.2% | 1.9% | 0.3% |
| | M+2 | 62.9% | 0.6% | 73.1% | 0.5% | 72.3% | 2.1% | 69.5% | 0.4% | 72.5% | 1.6% |
| | M+3 | 3.3% | 0.0% | 2.6% | 0.1% | 2.2% | 0.4% | 2.5% | 0.2% | 2.2% | 0.2% |
| | M+4 | 18.7% | 0.2% | 14.2% | 0.4% | 15.0% | 1.0% | 16.8% | 0.4% | 15.6% | 0.8% |
| | M+5 | 2.0% | 0.1% | 0.7% | 0.2% | 0.5% | 0.1% | 0.7% | 0.1% | 0.6% | 0.1% |
| | M+6 | 0.3% | 0.0% | 0.1% | 0.0% | 0.1% | 0.0% | 0.3% | 0.1% | 0.2% | 0.0% |
| 3PG | M+0 | 53.3% | 0.3% | 54.0% | 0.2% | 54.2% | 0.3% | 53.6% | 0.3% | 53.5% | 0.1% |
| | M+1 | 0.9% | 0.3% | 1.2% | 0.3% | 2.3% | 0.2% | 1.8% | 0.1% | 1.6% | 0.1% |
| | M+2 | 45.8% | 0.5% | 44.8% | 0.5% | 43.5% | 0.5% | 44.6% | 0.3% | 45.5% | 0.3% |
| | M+3 | 0.0% | 0.0% | 0.0% | 0.0% | 0.0% | 0.0% | 0.0% | 0.0% | 0.0% | 0.0% |
| 6PG | M+0 | 2.8% | 0.9% | 0.3% | 0.3% | 0.4% | 0.2% | 0.4% | 0.1% | 0.2% | 0.1% |
| | M+1 | 1.0% | 1.0% | 1.0% | 0.4% | 1.1% | 0.5% | 2.3% | 0.4% | 0.9% | 0.1% |
| | M+2 | 80.2% | 3.5% | 89.8% | 0.9% | 92.1% | 0.4% | 89.7% | 0.8% | 93.4% | 0.5% |
| | M+3 | 0.0% | 0.0% | 1.3% | 0.6% | 0.3% | 0.2% | 2.1% | 0.9% | 1.3% | 0.2% |
| | M+4 | 11.6% | 1.8% | 6.7% | 0.4% | 4.8% | 0.4% | 5.3% | 0.4% | 3.8% | 0.2% |
| | M+5 | 0.9% | 0.9% | 0.8% | 0.4% | 1.3% | 0.4% | 0.1% | 0.1% | 0.3% | 0.1% |
| | M+6 | 3.4% | 2.1% | 0.0% | 0.0% | 0.0% | 0.0% | 0.1% | 0.1% | 0.0% | 0.0% |

G6P denotes glucose-6-phosphate; F6P, fructose-6-phosphate; FBP, fructose-1,6-bisphosphate; 3PG, 3-phosphoglycerate; 6PG, 6-phosphogluconate. Values represent the mean and standard error (n=3 biological replicates).

**Supplementary Table 8. Isotopic labeling patterns of glycolytic intermediates in WT *E. coli* using [5,6-$^{13}C_2$]glucose in arginine to ammonia upshift**

| Metabolite | Labeling | 0 minutes post-upshift | s.e.m. | 3 minutes post-upshift | s.e.m. | 5 minutes post-upshift | s.e.m | 10 minutes post-upshift | s.e.m. | 30 minutes post-upshift | s.e.m |
|---|---|---|---|---|---|---|---|---|---|---|---|
| G6P/F6P | M+0 | 16.4% | 0.6% | 8.6% | 0.4% | 9.1% | 0.3% | 9.5% | 0.4% | 8.9% | 0.2% |
| | M+1 | 0.0% | 0.0% | 0.0% | 0.0% | 0.0% | 0.0% | 0.0% | 0.0% | 0.0% | 0.0% |
| | M+2 | 82.2% | 0.6% | 91.4% | 0.4% | 90.9% | 0.3% | 90.5% | 0.4% | 91.0% | 0.2% |
| | M+3 | 0.0% | 0.0% | 0.0% | 0.0% | 0.0% | 0.0% | 0.0% | 0.0% | 0.0% | 0.0% |
| | M+4 | 1.4% | 0.1% | 0.0% | 0.0% | 0.0% | 0.0% | 0.0% | 0.0% | 0.0% | 0.0% |
| | M+5 | 0.0% | 0.0% | 0.0% | 0.0% | 0.0% | 0.0% | 0.0% | 0.0% | 0.0% | 0.0% |
| | M+6 | 0.0% | 0.0% | 0.0% | 0.0% | 0.0% | 0.0% | 0.0% | 0.0% | 0.0% | 0.0% |
| FBP | M+0 | 21.9% | 0.5% | 19.6% | 0.3% | 19.1% | 0.7% | 19.1% | 0.6% | 17.3% | 0.9% |
| | M+1 | 0.0% | 0.0% | 0.0% | 0.0% | 0.0% | 0.0% | 0.0% | 0.0% | 0.0% | 0.0% |
| | M+2 | 68.1% | 0.4% | 72.7% | 0.9% | 72.9% | 1.0% | 73.9% | 0.7% | 76.8% | 1.4% |
| | M+3 | 0.0% | 0.0% | 0.0% | 0.0% | 0.0% | 0.0% | 0.0% | 0.0% | 0.0% | 0.0% |
| | M+4 | 9.9% | 0.4% | 7.4% | 0.8% | 8.1% | 0.5% | 7.0% | 1.2% | 5.8% | 0.6% |
| | M+5 | 0.0% | 0.0% | 0.0% | 0.0% | 0.0% | 0.0% | 0.0% | 0.0% | 0.0% | 0.0% |
| | M+6 | 0.1% | 0.1% | 0.2% | 0.2% | 0.0% | 0.0% | 0.0% | 0.0% | 0.0% | 0.0% |
| 3PG | M+0 | 51.9% | 0.6% | 49.0% | 0.3% | 47.1% | 0.6% | 45.0% | 0.0% | 45.9% | 0.4% |
| | M+1 | 0.0% | 0.0% | 0.0% | 0.0% | 0.0% | 0.0% | 0.0% | 0.0% | 0.0% | 0.0% |
| | M+2 | 48.1% | 0.6% | 51.0% | 0.3% | 52.9% | 0.6% | 55.0% | 0.0% | 54.1% | 0.4% |
| | M+3 | 0.0% | 0.0% | 0.0% | 0.0% | 0.0% | 0.0% | 0.0% | 0.0% | 0.0% | 0.0% |
| 6PG | M+0 | 14.1% | 0.3% | 9.4% | 0.8% | 11.2% | 0.2% | 8.8% | 0.7% | 6.8% | 1.0% |
| | M+1 | 0.0% | 0.0% | 0.0% | 0.0% | 0.0% | 0.0% | 0.0% | 0.0% | 0.0% | 0.0% |
| | M+2 | 85.9% | 0.3% | 90.6% | 0.8% | 88.6% | 0.4% | 91.2% | 0.7% | 93.2% | 1.0% |
| | M+3 | 0.0% | 0.0% | 0.0% | 0.0% | 0.2% | 0.2% | 0.0% | 0.0% | 0.0% | 0.0% |
| | M+4 | 0.0% | 0.0% | 0.0% | 0.0% | 0.0% | 0.0% | 0.0% | 0.0% | 0.0% | 0.0% |
| | M+5 | 0.0% | 0.0% | 0.0% | 0.0% | 0.0% | 0.0% | 0.0% | 0.0% | 0.0% | 0.0% |
| | M+6 | 0.0% | 0.0% | 0.0% | 0.0% | 0.0% | 0.0% | 0.0% | 0.0% | 0.0% | 0.0% |

G6P denotes glucose-6-phosphate; F6P, fructose-6-phosphate; FBP, fructose-1,6-bisphosphate; 3PG, 3-phosphoglycerate; 6PG, 6-phosphogluconate. Values represent the mean and standard error (n=3 biological replicates).

**Supplementary Table 9. Isotopic labeling patterns of glycolytic intermediates in WT *E. coli* using [1,2-$^{13}C_2$]glucose in no nitrogen to ammonia upshift**

| Metabolite | Labeling | 0 minutes post-upshift | s.e.m. | 3 minutes post-upshift | s.e.m. | 5 minutes post-upshift | s.e.m | 10 minutes post-upshift | s.e.m. | 30 minutes post-upshift | s.e.m |
|---|---|---|---|---|---|---|---|---|---|---|---|
| G6P/F6P | M+0 | 8.7% | 0.4% | 1.1% | 0.5% | 1.3% | 0.3% | 1.1% | 0.4% | 0.9% | 0.2% |
| | M+1 | 2.4% | 0.1% | 0.4% | 0.1% | 0.1% | 0.1% | 0.1% | 0.0% | 0.2% | 0.1% |
| | M+2 | 62.6% | 0.4% | 92.6% | 0.3% | 92.5% | 0.3% | 93.7% | 0.3% | 94.9% | 0.3% |
| | M+3 | 7.1% | 0.1% | 1.0% | 0.3% | 0.8% | 0.1% | 1.0% | 0.2% | 0.8% | 0.2% |
| | M+4 | 14.6% | 0.4% | 4.2% | 0.1% | 4.6% | 0.2% | 3.6% | 0.1% | 2.8% | 0.1% |
| | M+5 | 4.3% | 0.2% | 0.6% | 0.0% | 0.5% | 0.1% | 0.5% | 0.0% | 0.4% | 0.0% |
| | M+6 | 0.2% | 0.0% | 0.1% | 0.0% | 0.0% | 0.0% | 0.1% | 0.0% | 0.1% | 0.0% |
| FBP | M+0 | 20.3% | 0.6% | 10.3% | 0.2% | 10.9% | 0.5% | 9.6% | 0.2% | 8.9% | 0.8% |
| | M+1 | 2.9% | 0.1% | 1.0% | 0.1% | 0.6% | 0.1% | 0.9% | 0.0% | 1.0% | 0.1% |
| | M+2 | 48.9% | 0.2% | 69.1% | 0.4% | 67.0% | 0.5% | 67.8% | 0.3% | 67.7% | 1.1% |
| | M+3 | 6.7% | 0.3% | 1.3% | 0.4% | 1.3% | 0.0% | 1.5% | 0.1% | 1.5% | 0.1% |
| | M+4 | 17.1% | 0.6% | 17.3% | 0.6% | 19.4% | 0.5% | 19.2% | 0.1% | 19.9% | 0.3% |
| | M+5 | 3.9% | 0.2% | 0.7% | 0.1% | 0.6% | 0.1% | 0.7% | 0.0% | 0.6% | 0.1% |
| | M+6 | 0.2% | 0.1% | 0.2% | 0.0% | 0.3% | 0.0% | 0.3% | 0.0% | 0.3% | 0.0% |
| 3PG | M+0 | 52.8% | 0.1% | 53.7% | 0.5% | 54.7% | 0.2% | 55.2% | 0.4% | 54.3% | 0.1% |
| | M+1 | 2.8% | 0.4% | 0.9% | 0.4% | 0.8% | 0.4% | 1.3% | 0.2% | 1.6% | 0.2% |
| | M+2 | 40.4% | 0.3% | 44.8% | 0.6% | 44.3% | 0.7% | 43.2% | 0.3% | 44.1% | 0.1% |
| | M+3 | 3.9% | 0.5% | 0.5% | 0.5% | 0.1% | 0.1% | 0.4% | 0.1% | 0.0% | 0.0% |
| 6PG | M+0 | 6.5% | 0.5% | 0.5% | 0.1% | 0.5% | 0.2% | 0.6% | 0.1% | 0.4% | 0.1% |
| | M+1 | 3.0% | 0.4% | 0.2% | 0.2% | 0.0% | 0.0% | 0.0% | 0.0% | 0.1% | 0.1% |
| | M+2 | 69.8% | 0.3% | 96.8% | 0.2% | 96.3% | 0.6% | 97.0% | 0.2% | 97.2% | 0.4% |
| | M+3 | 5.5% | 1.3% | 0.0% | 0.0% | 0.0% | 0.0% | 0.2% | 0.2% | 0.1% | 0.1% |
| | M+4 | 12.8% | 1.0% | 2.5% | 0.1% | 3.0% | 0.5% | 2.0% | 0.1% | 2.0% | 0.2% |
| | M+5 | 2.4% | 0.8% | 0.0% | 0.0% | 0.1% | 0.0% | 0.1% | 0.0% | 0.2% | 0.1% |
| | M+6 | 0.0% | 0.0% | 0.0% | 0.0% | 0.0% | 0.0% | 0.0% | 0.0% | 0.0% | 0.0% |

G6P denotes glucose-6-phosphate; F6P, fructose-6-phosphate; FBP, fructose-1,6-bisphosphate; 3PG, 3-phosphoglycerate; 6PG, 6-phosphogluconate. Values represent the mean and standard error (n=3 biological replicates).

**Supplementary Table 10. Isotopic labeling patterns of glycolytic intermediates in WT *E. coli* using [5,6-$^{13}C_2$]glucose in no nitrogen to ammonia upshift**

| Metabolite | Labeling | 0 minutes post-upshift | s.e.m. | 3 minutes post-upshift | s.e.m. | 5 minutes post-upshift | s.e.m | 10 minutes post-upshift | s.e.m. | 30 minutes post-upshift | s.e.m |
|---|---|---|---|---|---|---|---|---|---|---|---|
| G6P/F6P | M+0 | 27.1% | 0.6% | 13.1% | 0.3% | 14.4% | 0.3% | 10.4% | 0.2% | 8.1% | 0.2% |
| | M+1 | 0.8% | 0.1% | 0.0% | 0.0% | 0.0% | 0.0% | 0.0% | 0.0% | 0.0% | 0.0% |
| | M+2 | 63.7% | 0.5% | 86.9% | 0.3% | 85.6% | 0.3% | 89.6% | 0.2% | 91.9% | 0.2% |
| | M+3 | 2.0% | 0.1% | 0.0% | 0.0% | 0.0% | 0.0% | 0.0% | 0.0% | 0.0% | 0.0% |
| | M+4 | 6.2% | 0.1% | 0.0% | 0.0% | 0.0% | 0.0% | 0.0% | 0.0% | 0.0% | 0.0% |
| | M+5 | 0.2% | 0.0% | 0.0% | 0.0% | 0.0% | 0.0% | 0.0% | 0.0% | 0.0% | 0.0% |
| | M+6 | 0.1% | 0.0% | 0.0% | 0.0% | 0.0% | 0.0% | 0.0% | 0.0% | 0.0% | 0.0% |
| FBP | M+0 | 22.7% | 0.3% | 18.7% | 0.2% | 19.2% | 0.4% | 19.0% | 0.1% | 19.1% | 0.1% |
| | M+1 | 1.0% | 0.2% | 0.0% | 0.0% | 0.0% | 0.0% | 0.0% | 0.0% | 0.0% | 0.0% |
| | M+2 | 54.3% | 0.7% | 75.7% | 0.3% | 74.7% | 0.6% | 74.1% | 0.1% | 73.3% | 0.2% |
| | M+3 | 1.6% | 0.2% | 0.0% | 0.0% | 0.0% | 0.0% | 0.0% | 0.0% | 0.0% | 0.0% |
| | M+4 | 19.8% | 0.1% | 5.6% | 0.1% | 6.1% | 0.2% | 6.9% | 0.2% | 7.5% | 0.1% |
| | M+5 | 0.3% | 0.0% | 0.0% | 0.0% | 0.0% | 0.0% | 0.0% | 0.0% | 0.0% | 0.0% |
| | M+6 | 0.3% | 0.0% | 0.0% | 0.0% | 0.0% | 0.0% | 0.0% | 0.0% | 0.0% | 0.0% |
| 3PG | M+0 | 46.7% | 0.6% | 45.3% | 0.6% | 45.3% | 1.5% | 43.9% | 0.5% | 44.8% | 0.5% |
| | M+1 | 1.2% | 0.4% | 0.0% | 0.0% | 0.0% | 0.0% | 0.0% | 0.0% | 0.0% | 0.0% |
| | M+2 | 52.1% | 0.3% | 54.7% | 0.6% | 54.7% | 1.5% | 56.1% | 0.5% | 55.2% | 0.5% |
| | M+3 | 0.0% | 0.0% | 0.0% | 0.0% | 0.0% | 0.0% | 0.0% | 0.0% | 0.0% | 0.0% |
| 6PG | M+0 | 16.6% | 0.3% | 2.8% | 0.1% | 4.0% | 0.1% | 3.3% | 0.1% | 2.6% | 0.1% |
| | M+1 | 0.8% | 0.2% | 0.0% | 0.0% | 0.0% | 0.0% | 0.0% | 0.0% | 0.0% | 0.0% |
| | M+2 | 76.1% | 0.7% | 97.2% | 0.1% | 96.0% | 0.1% | 96.7% | 0.1% | 97.4% | 0.1% |
| | M+3 | 1.6% | 0.2% | 0.0% | 0.0% | 0.0% | 0.0% | 0.0% | 0.0% | 0.0% | 0.0% |
| | M+4 | 4.7% | 0.8% | 0.0% | 0.0% | 0.0% | 0.0% | 0.0% | 0.0% | 0.0% | 0.0% |
| | M+5 | 0.1% | 0.1% | 0.0% | 0.0% | 0.0% | 0.0% | 0.0% | 0.0% | 0.0% | 0.0% |
| | M+6 | 0.0% | 0.0% | 0.0% | 0.0% | 0.0% | 0.0% | 0.0% | 0.0% | 0.0% | 0.0% |

G6P denotes glucose-6-phosphate; F6P, fructose-6-phosphate; FBP, fructose-1,6-bisphosphate; 3PG, 3-phosphoglycerate; 6PG, 6-phosphogluconate. Values represent the mean and standard error (n=3 biological replicates).

**Supplementary Table 11. The ED-to-EMP pathway flux ratios before and after nutrient upshift**

| Starting condition | 0 minutes post-upshift | s.e.m. | 3 minutes post-upshift | s.e.m. | 5 minutes post-upshift | s.e.m | 10 minutes post-upshift | s.e.m. | 30 minutes post-upshift | s.e.m |
|---|---|---|---|---|---|---|---|---|---|---|
| Ace to Glc | - | - | 1.694 | 0.050 | 1.094 | 0.027 | 0.456 | 0.016 | 0.296 | 0.005 |
| Ace to Glc* | - | - | 1.289 | 0.042 | 0.457 | 0.016 | 0.105 | 0.006 | 0.031 | 0.004 |
| Arg to Ammonia | 0.128 | 0.022 | 0.158 | 0.004 | 0.151 | 0.023 | 0.125 | 0.022 | 0.110 | 0.005 |
| No N to Ammonia | 0.180 | 0.023 | 0.160 | 0.026 | 0.201 | 0.008 | 0.225 | 0.023 | 0.165 | 0.011 |

"Ace to Glc" upshfit uses data from the [1,2-$^{13}C_2$]glucose experiment that is corrected for incomplete labeling by the [U-$^{13}C_6$]glucose experiment. "Ace to Glc*" upshfit uses data from the same [1,2-$^{13}C_2$]glucose experiment but corrected both for incomplete labeling using [U-$^{13}C_6$]glucose and possible reverse labeling using [5,6-$^{13}C_2$]glucose. Values represent the mean and standard error (n=3 biological replicates).

**Supplementary Table 12. Gibbs free energies of the ED and the EMP pathways during carbon upshift in WT**

| Pathway | Net reaction | 0 minutes post-upshift | s.e.m. | 3 minutes post-upshift | s.e.m. | 5 minutes post-upshift | s.e.m | 10 minutes post-upshift | s.e.m. | 30 minutes post-upshift | s.e.m |
|---|---|---|---|---|---|---|---|---|---|---|---|
| EMP | G6P + ADP + 2 NAD + 2 Pi = 2 PEP + ATP + 2 NADH + 2 H2O | -45.50 | 1.69 | -54.79 | 2.22 | -51.05 | 2.07 | -51.80 | 1.75 | -46.49 | 1.38 |
| ED | G6P + ADP + NAD + NADP + Pi = PYR + PEP + ATP + NADH + NADPH + H2O | -81.77 | 4.78 | -87.86 | 3.74 | -85.80 | 4.24 | -87.16 | 4.43 | -79.57 | 5.67 |
| GNG | 2 PEP + 2 ATP + 2 NADH + 3 H2O = G6P + 2 ADP + 2 NAD + 3 Pi | -2.27 | 1.41 | 8.55 | 2.44 | 2.83 | 1.85 | 6.84 | 2.04 | -0.30 | 0.49 |

Values represent the mean and standard error (n=3 biological replicates).

**Supplementary Table 13. Gibbs free energies of the ED and the EMP pathways during arginine-to-ammonia nitrogen upshift in WT**

| Pathway | Net reaction | 0 minutes post-upshift | s.e.m. | 3 minutes post-upshift | s.e.m. | 5 minutes post-upshift | s.e.m | 10 minutes post-upshift | s.e.m. | 30 minutes post-upshift | s.e.m |
|---|---|---|---|---|---|---|---|---|---|---|---|
| EMP | G6P + ADP + 2 NAD + 2 Pi = 2 PEP + ATP + 2 NADH + 2 H2O | -53.40 | 0.30 | -49.81 | 0.43 | -49.74 | 1.02 | -49.07 | 0.35 | -47.50 | 0.04 |
| ED | G6P + ADP + NAD + NADP + Pi = PYR + PEP + ATP + NADH + NADPH + H2O | -93.43 | 3.23 | -91.39 | 3.44 | -91.55 | 3.26 | -90.63 | 3.92 | -88.79 | 3.40 |
| ED + PpsA | G6P + ADP + NAD + NADP = 2 PEP + NADH + NADPH + AMP | -120.90 | 0.55 | -118.85 | 0.45 | -119.82 | 0.48 | -117.96 | 1.04 | -117.95 | 1.39 |

Values represent the mean and standard error (n=3 biological replicates).

**Supplementary Table 14. Gibbs free energies of the ED and the EMP pathways during ammonia upshift in WT**

| Pathway | Net reaction | 0 minutes post-upshift | s.e.m. | 3 minutes post-upshift | s.e.m. | 5 minutes post-upshift | s.e.m | 10 minutes post-upshift | s.e.m. | 30 minutes post-upshift | s.e.m |
|---|---|---|---|---|---|---|---|---|---|---|---|
| EMP | G6P + ADP + 2 NAD + 2 Pi = 2 PEP + ATP + 2 NADH + 2 H2O | -43.90 | 0.85 | -42.17 | 0.52 | -41.67 | 0.85 | -42.95 | 0.96 | -43.87 | 0.03 |
| ED | G6P + ADP + NAD + NADP + Pi = PYR + PEP + ATP + NADH + NADPH + H2O | -88.90 | 0.26 | -84.21 | 0.63 | -81.81 | 0.31 | -83.85 | 0.77 | -85.36 | 0.01 |
| ED + PpsA | G6P + ADP + NAD + NADP = 2 PEP + NADH + NADPH + AMP | -121.27 | 0.69 | -118.99 | 0.49 | -118.23 | 0.98 | -119.51 | 0.24 | -117.94 | 0.90 |

Values represent the mean and standard error (n=3 biological replicates).

**Supplementary Table 15. Gibbs free energies of the ED and the EMP pathways during carbon upshift in Δ*edd***

| Pathway | Net reaction | 0 minutes post-upshift | s.e.m. | 3 minutes post-upshift | s.e.m. | 5 minutes post-upshift | s.e.m | 10 minutes post-upshift | s.e.m. | 30 minutes post-upshift | s.e.m |
|---|---|---|---|---|---|---|---|---|---|---|---|
| EMP | G6P + ADP + 2 NAD + 2 Pi = 2 PEP + ATP + 2 NADH + 2 H2O | -50.03 | 1.69 | -49.24 | 1.78 | -49.15 | 5.30 | -49.55 | 2.83 | -47.48 | 0.09 |
| ED | G6P + ADP + NAD + NADP + Pi = PYR + PEP + ATP + NADH + NADPH + H2O | -87.08 | 2.38 | -88.82 | 1.80 | -88.25 | 4.33 | -87.76 | 1.45 | -85.39 | 0.30 |
| GNG | 2 PEP + 2 ATP + 2 NADH + 3 H2O = G6P + 2 ADP + 2 NAD + 3 Pi | -0.33 | 3.23 | 1.73 | 3.07 | 2.34 | 7.16 | 2.27 | 4.54 | -1.26 | 0.01 |

Values represent the mean and standard error (n=3 biological replicates).

**Supplementary Table 16. Gibbs free energies of the ED and the EMP pathways during arginine-to-ammonia nitrogen upshift in Δ*edd***

| Pathway | Net reaction | 0 minutes post-upshift | s.e.m. | 3 minutes post-upshift | s.e.m. | 5 minutes post-upshift | s.e.m | 10 minutes post-upshift | s.e.m. | 30 minutes post-upshift | s.e.m |
|---|---|---|---|---|---|---|---|---|---|---|---|
| EMP | G6P + ADP + 2 NAD + 2 Pi = 2 PEP + ATP + 2 NADH + 2 H2O | -53.06 | 1.83 | -48.24 | 0.22 | -44.73 | 1.39 | -47.26 | 0.98 | - | - |
| ED | G6P + ADP + NAD + NADP + Pi = PYR + PEP + ATP + NADH + NADPH + H2O | -88.55 | 2.06 | -87.79 | 2.03 | -85.88 | 2.47 | -87.22 | 0.75 | - | - |
| ED + PpsA | G6P + ADP + NAD + NADP = 2 PEP + NADH + NADPH + AMP | -132.18 | 1.72 | -128.65 | 3.16 | -125.98 | 0.89 | -64.08 | 0.82 | - | - |

Values represent the mean and standard error (n=3 biological replicates).

**Supplementary Table 17. Gibbs free energies of the ED and the EMP pathways during ammonia upshift in Δ*edd***

| Pathway | Net reaction | 0 minutes post-upshift | s.e.m. | 3 minutes post-upshift | s.e.m. | 5 minutes post-upshift | s.e.m | 10 minutes post-upshift | s.e.m. | 30 minutes post-upshift | s.e.m |
|---|---|---|---|---|---|---|---|---|---|---|---|
| EMP | G6P + ADP + 2 NAD + 2 Pi = 2 PEP + ATP + 2 NADH + 2 H2O | -44.39 | 2.43 | -42.44 | 0.64 | -42.90 | 0.43 | -45.48 | 0.91 | -47.48 | 0.09 |
| ED | G6P + ADP + NAD + NADP + Pi = PYR + PEP + ATP + NADH + NADPH + H2O | -86.16 | 1.38 | -84.31 | 0.97 | -83.13 | 1.36 | -87.05 | 0.61 | -86.15 | 0.68 |
| ED + PpsA | G6P + ADP + NAD + NADP = 2 PEP + NADH + NADPH + AMP | -125.00 | 0.11 | -123.62 | 0.36 | -123.16 | 0.42 | -126.85 | 0.57 | -126.39 | 0.39 |

Values represent the mean and standard error (n=3 biological replicates).
